# Heme Hopping Falls Short: What Explains Anti-Arrhenius Conductivity in a Multi-heme Cytochrome Nanowire?

**DOI:** 10.1101/2022.08.01.502099

**Authors:** Matthew J. Guberman-Pfeffer

**Affiliations:** Department of Molecular Biophysics and Biochemistry, Yale University, New Haven, CT, U.S.A.; Microbial Sciences Institute, Yale University, West Haven, CT, U.S.A

**Keywords:** *anti*-Arrhenius, hopping, conductivity, multi-heme, nanowire, OmcS, redox potential, QM/MM, constant redox molecular dynamics

## Abstract

A helical homopolymer of the outer-membrane cytochrome type S (OmcS) was proposed to electrically connect a common soil bacterium, *Geobacter sulfurreducens*, with minerals and other microbes for biogeochemically important processes. OmcS exhibits a surprising rise in conductivity upon cooling from 300 to 270 K that has recently been attributed to a restructuring of H-bonds, which in turn modulates heme redox potentials. This proposal is more thoroughly examine herein by (1) analyzing H-bonding at 13 temperatures encompassing the entire experimental range; (2) computing redox potentials with quantum mechanics/molecular mechanics for 10-times more (3000) configurations sampled from 3-times longer (2 μs) molecular dynamics, as well as 3 μs of constant redox and pH molecular dynamics; and (3) modeling redox conduction with both single-particle diffusion and multi-particle flux kinetic schemes. Upon cooling by 30 K, the connectivity of the intra-protein H-bonding network was highly (86%) similar. An increase in the density and static dielectric constant of the filament’s hydration shell caused a −0.002 V/K shift in heme redox potentials, and a factor of 2 decrease in charge mobility. Revision of a too-far negative redox potential in prior work (−0.521 V; expected = −0.350 – +0.150 V; new Calc. = −0.214 V vs. SHE) caused the mobility to be greater at high *versus* low temperature, opposite to the original prediction. These solution-phase redox conduction models failed to reproduce the experimental conductivity of electrode-absorbed, partially dehydrated, and possibly aggregated OmcS filaments. Some improvement was seen by neglecting reorganization energy from the solvent to model dehydration. Correct modeling of the physical state is suggested to be a prerequisite for reaching a verdict on the operative charge transport mechanism and the molecular basis of its temperature response.

## 1. Introduction

All respiratory life is powered by the flow of electrons through a chain of reduction-oxidation (redox)-active cofactors.^1, 2^ Electrons enter the chain by oxidation of a primary donor (e.g., organic matter), and exit via reduction of a terminal acceptor (e.g., O_2_),^3^ both of which are ordinarily internalized by a cell. The nearest electron donor/acceptor in some aquatic sediments, however, is either another microbe or a mineral nanoparticle that may be up to a few microns away. Coupling the reduction of microbes or minerals in the extracellular space with intracellular oxidative metabolism constitutes an ancient and widespread respiratory strategy.^4–7^ The electrical circuits established between microbes and minerals, if better understood, can elucidate the biogeochemical evolution of the planet,^8–13^ the role of the microbiome in human health and disease,^14–17^ as well as templates^18–21^ for the design of bioelectronic technologies.^21–34^

Microbes can electrically ‘plug-in’ to their environments through direct contact with cell-surface proteins, outer-membrane vesicles, filamentous appendages, or molecular shuttles. One of the best studied electroactive microorganisms, *Geobacter sulfurreducens*, has been known for nearly twenty years to produce conductive filaments.^35^ Over that time, questions about the identity, composition, and conductivity mechanism of the filaments have^36–41^ and continue^42–54^ to be discussed.

Homopolymers of a mostly α-helical pilin protein^46, 51^ or outer-membrane multi-heme cytochromes^45, 55^ have been proposed as the “nanowires” used by *G. sulfurreducens*. The pilin-based polymer, called a pilus, is a common bacterial structure that usually facilitates adhesion to other cells or surfaces, motility, secretion, horizontal gene transfer, biofilm formation, and can serve as a virulence factor.^56^ Electrical conductivity was proposed as a uniquely evolved function of the *G. sulfurreducens* pilus due to a hypothesized continuous chain of aromatic residues (Figure 1, *top left*).^57, 58^

**Figure 1.**
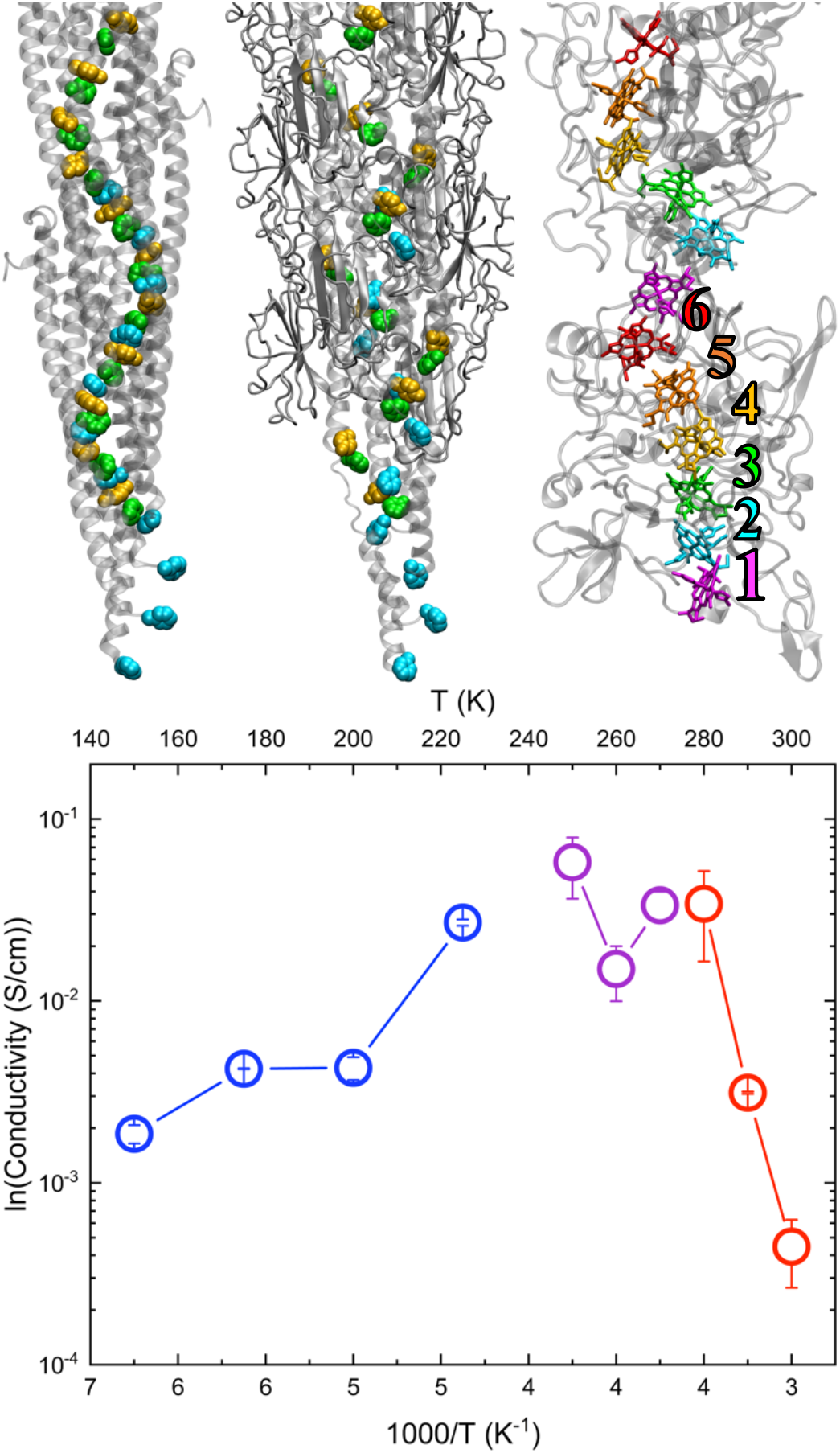
*Top*: Structures of the (*left*) hypothetical *G. sulfurreducens* pilus from Ref. 74, (*middle*) CryoEM model of the pilus from Ref. 50, and (*right*) CryoEM model of the outer-membrane cytochrome type S (OmcS) filament from Ref. 45. In the pilus models, the Phe-1 (cyan), Phe-24 (green) and Tyr-27 (orange) from different pilin subunits in the non-covalent assembly are shown in van der Waals representation. For OmcS, hemes **#1** through **#6** in each subunit are colored magenta, cyan, green, yellow, orange, and red, respectively. (*bottom*) Temperature-dependent electrical conductivity of OmcS. The data is reproduced from Figure 4A in Ref. 83.

In the absence of atomic-scale structural data until very recently (Figure 1, *top middle*),^50^ this suggestion spurred the creation of model systems^59–69^ and theoretical studies based on homology modeling, docking, and molecular dynamics.^70–80^ Computed conductivities for the hypothetical pilus have ranged from orders of magnitude too small^73^ to quantitatively accurate^77^ relative to experimental measurements attributed to the pilus. Proposed mechanisms include multistep hopping between aromatic sidechains,^72^ transport through aromatic clusters with delocalized states,^76^ or aromatic sidechains and cytochromes thought to decorate the pili,^71^ or band-like conductivity through π-π stacked aromatic residues.^74^

Alternatively, the electrically conductive or *e*-pili (Figure 1, *top left*) have been suggested to not even exist as such,^45, 50^ being instead misidentified polymeric cytochromes (e.g., Figure 1, *top right*).^45^ There is a small but growing library of multi-heme cytochromes that assemble into homopolymeric filaments.^45, 55, 81^ Structural and functional characterizations of these cytochrome filaments have given theoretical studies a firmer foundation—in principle--than for the putative *e*-pili. Even so, proposed electrical conductivity mechanisms for the outer-membrane cytochrome type S (OmcS) filament range from incoherent charge hopping under physiological^82^ or solid-state^83^ measurement conditions, to coherence-assisted charge hopping,^84^ and decoherent quantum transport.^85^

In a very recent study led by Dahl *et al*. (including the present author),^83^ one of the incoherent charge hopping models for OmcS was proposed. Charge-hopping is a thermally activated process that should become exponentially less favorable upon cooling. Surprisingly, the conductivity first increased upon cooling down to a transition temperature, and then decreased upon further cooling (Figure 1, *bottom*). The conductivity at 150 K remained 4-times larger than at 300 K. The same behavior was previously reported and ascribed to the *G. sulfurreducens* pilus.^36^

To explain the unexpected initial rise in conductivity upon cooling (*anti*-Arrhenius behavior), a temperature-sensitive massive restructuring of the intra-protein H-bonding network was suggested to modulate the redox potentials of the heme cofactors. The *c*-type bis-histidine ligated hemes of OmcS constituted the charge relaying sites in the conductivity model.

Temperature-induced shifts in redox potentials for cytochromes are known,^86–93^ but the computed shifts were unexpectedly large and of mixed sign (e.g., −0.189 and +0.113 V for hemes in van der Waals contact). A global descriptor was invoked to characterize cooling-induced restructuring of the H-bonding network, but it could not explain the distinct temperature dependencies of the six hemes in a subunit of the filament. Also, the redox potential for one of the hemes was unusually negative at 310 K (−0.521 V *vs*. SHE; expected range = −0.350 to +0.150 V vs. SHE),^94^ which may have skewed the comparison of computed conductivities at different temperatures.

Herein, the temperature dependence of structural transitions and redox conduction in OmcS are more thoroughly examined. H-bonding patterns were now analyzed at temperatures from 100 to 400 K in 25 K increments to encompass the entire range examined experimentally. At 270 and 300 K, which bracket the *anti*-Arrhenius regime, redox potentials were computed as thermal averages over 10-times more (3000) configurations, sampled from 3-times longer (2 μs) molecular dynamics (MD), using quantum mechanics/molecular mechanics (QM/MM) with an extensively benchmarked density functional for iron porphyrins, and compared to results from ~3 μs of constant redox and pH molecular dynamics (C(E,pH)MD). Redox conduction was then modeled using both a single-particle diffusion and a multi-particle flux model, thereby allowing comparisons to all prior theoretical work on OmcS that only used one or the other.

As discussed below, these extensive analyses indicated that *none* of the charge-hopping models described to-date, including the present one, can explain the electrical conductivity of OmcS quantitatively, or qualitatively as a function of temperature. These discrepancies may imply that either the hopping mechanism is inappropriate to describe the electrical conductivity, or that the simulations do not capture important physical aspects of the experimental conditions. The results indicated an important coupling of the electron transfers with solvent structure and dynamics, but the level of hydration in experiments has not been well-characterized. Entropy effects, solvent viscosity, and non-ergodicity have all been used to interpret other *anti*-Arrhenius phenomena and may apply to OmcS.

## 2. Results

### 2.1. Overview

Heme redox potentials at 270 and 300 K are first analyzed (§2.2). These temperatures bracket the range for the observed *anti*-Arrhenius behavior in electrical conductivity (Figure 1, *bottom*). The microscopic origins for the shifts in redox potentials upon cooling are then delineated in terms of altered electrostatic interactions and H-bonding patterns (§2.3). Finally, the temperature dependence of redox conduction is analyzed with multiple kinetic schemes and compared to prior experimental and theoretical work (§2.4). The methods are extensively detailed in the Supporting Information (SI). Discussion of the results is reserved for §3.

### 2.2. Temperature Dependence of Redox Potentials

Figure 2 (Table S13) compares the redox potentials (*E*°) for the heme cofactors in OmcS computed at high (310 or 300 K) and low (270 K) temperatures in the present (*left*) and prior (*right*) works.^83^ The numbering scheme in Figure 1 is used throughout the article with the hemes designated as **#X** (X = 1 – 6).

**Figure 2.**
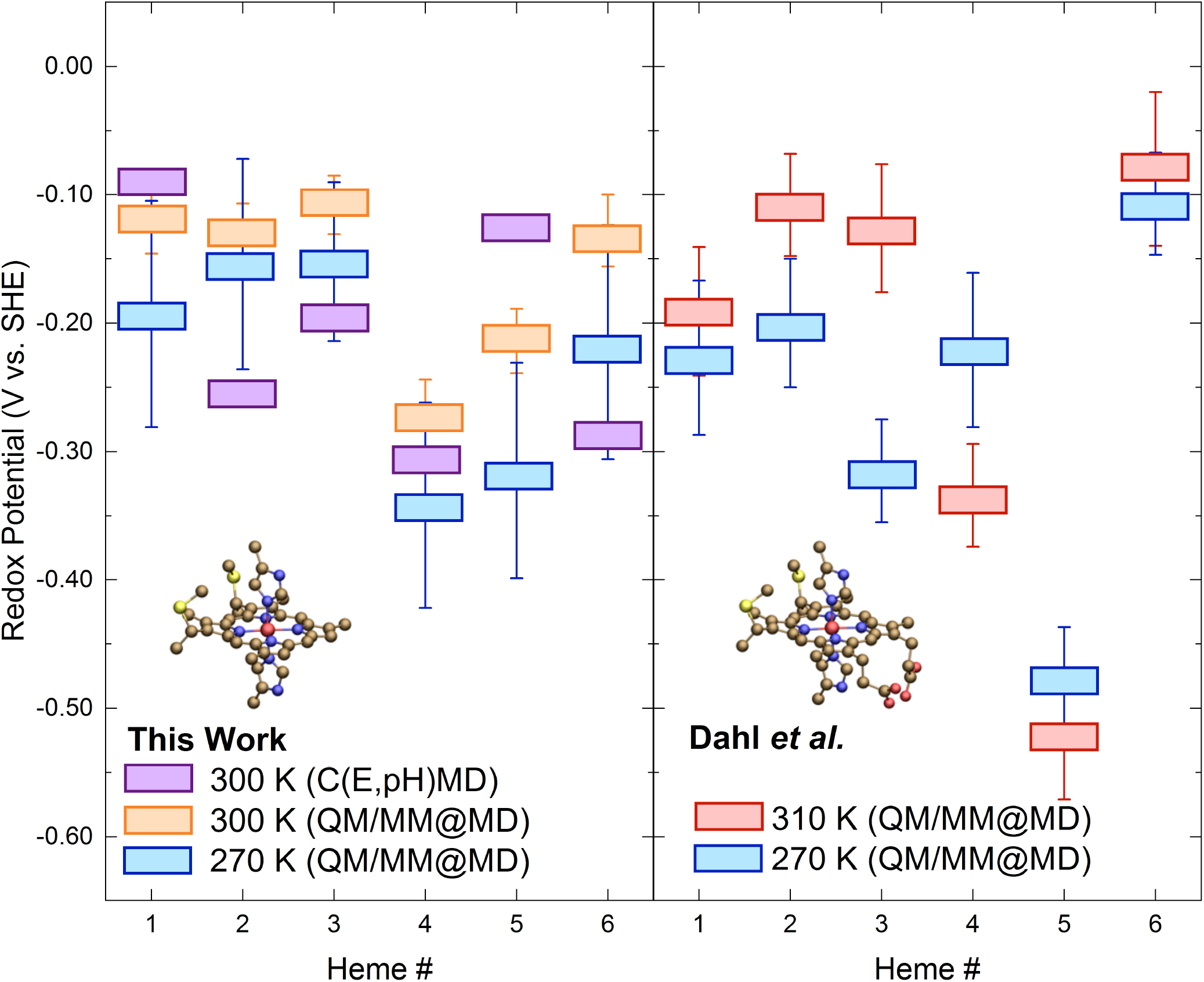
Comparison of redox potentials computed at high (310 or 300 K) and low (270 K) temperatures in (*left*) the present work using constant redox and pH molecular dynamics (C(E,pH)MD) and quantum mechanics/molecular mechanics at MD configurations (QM/MM@MD) with B3LYP/[Fe = LANL2DZ; H, C, N, S = 6-31G(d)]:AMBER99SB, or (*right*) Ref. 83 using QM/MM@MD with ωb97XD/[Fe = LANL2DZ; H, C, N, S, O = cc-pVDZ]:CHAMM36. The inset on each side shows the definition of the QM region, which differed by the (*left*) exclusion or (*right*) inclusion) of the propionic acid groups.

The computed *E*° values in both the present and prior works were obtained under the linear response approximation as the average of the vertical ionization potential and the negative of the vertical electron affinity for each heme (see §S1.3.2 in the SI).^95–97^ These vertical energy gaps were computed from quantum mechanical/molecular mechanical (QM/MM) single point energy evaluations on configurations generated from classical molecular dynamics (MD) trajectories; an approach herein referred to as QM/MM@MD. Table S1 gives a detailed comparison of the implementations in the present and prior^83^ works. The present work is distinguished by:

1. Use of a forcefield specifically parameterized for bis-histidine *c*-type hemes^98, 99^ that has been shown to reasonably estimate *E*° within ~0.120 V from electrostatic free energy differences^100, 101^ and that accurately describes *relative* differences in electric fields.^102^
2. Propagation of separate simulations at 100 to 400 K in 25 K increments, instead of only 270 and 310 K, to characterize temperature-dependent structural transitions. Note that 270 instead of 275 K was used for direct comparison with prior work^83^ at that temperature.
3. Propagation of multiple (3-5) trajectories at 270 K in the oxidized and reduced states for each of the six hemes in OmcS. Each trajectory was spawned from the preceding one and rendered independent by randomizing the velocities.
4. Sampling of 10-times or more (180-370 *versus* 25) configurations for QM/MM energy evaluations in each of two redox states (oxidized and reduced) for every heme to compute *E*° at 300 (Table S4-S6; Figure S2, S3) and 270 K (Tables S7; Figures S3-S6). See §S2 for a discussion on the criteria used to assess the convergence of *E*° at 270 K.
5. Use of an approximate density functional, B3LYP, that has been extensively benchmarked for heme or other iron porphyrin systems^103–111^
6. Use of both double- and triple-ζ basis sets to ensure the QM results were converged with respect to this parameter (Tables S4, S5; Figure S3).
7. Assessment of the approximation made in the prior^83^ work of using heme geometries from MD trajectories for QM calculations without further optimization (Table S9).

In addition to these methodological refinements, *E*°s computed with QM/MM@MD were now compared with those from nearly 3.0 μs of constant redox or constant redox and pH molecular dynamics (CE- or C(E,pH)MD) simulations (Table S3).^101, 112^ Separately, nearly 1.0 μs of constant pH molecular dynamics (CpHMD) was performed in two different redox microstates (i.e., combinations of oxidized and/or reduced hemes) (Table S2). These classical electrostatics-based techniques revealed the influence of heme-heme and heme-acid/base (redox-Bohr) interactions on the computed *E*°s (Table S14; Figure S8) and pK_a_s (Table S29, S30; Figure S11). Doing these tests at the QM/MM@MD level for a multi-cofactor system is currently computationally prohibitive. The influence of both heme-heme and redox-Bohr interactions on *E*° were assumed in the QM/MM@MD computations to be comparable to the ≤0.08 V standard error of the mean obtained with this approach.

#### 2.2.1. High (300-310 K) Temperature

*E*°s at 300 K for the hemes of OmcS were −0.063 – −0.271 V vs. SHE according to QM/MM@MD (Figure 2, Table S13). Use of a triple-instead of double-ζ basis set shifted the values towards the positive end of the range by 0.034 ± 0.013 (mean ± standard deviation) V, a magnitude comparable to the standard error of the mean for each *E*° (Tables S4, S5; Figure S3). The range for *E*° at 300 K was likewise −0.093 – −0.306 V vs. SHE according to C(E,pH)MD simulations (Figure 2, Table S14). These ranges were consistent with the experimental range of −0.350 – +0.150 V vs. SHE reported for bis-histidine ligated *c*-type hemes.^94^ Thus, the computed *E*°s were converged with respect to basis set quality, consistent at multiple levels of theory, and in good agreement with experimental expectations. Results presented below were obtained with the double-ζ basis set unless otherwise noted.

Along the linear heme chain from **#1** → **#2** … **#6** in OmcS (and the x-axis in Figure 2), the *E*°s at 300 K fluctuated within |0.022| V from **#1** to **#3**, plummeted by −0.163 V from **#3** to **#4**, and recovered to within 0.025 V (thermal energy at 290 K) of the starting *E*° from **#4** to **#6**. From the viewpoint of the electron transport function of OmcS, it is remarkable that the protein tunes the *E*°s of six chemically identical heme groups over ~0.2 V, and yet, arranges the hemes so that there is no net change in *E*° through a subunit of the filament. A similar “design” strategy was found for the deca-heme protein MtrF from *Shewanella oneidensis*.^113^ The strategy is also consistent with the need for long-range charge transport through a homopolymer, perhaps supporting the physiological relevance of the OmcS filament. The mechanisms regulating the redox profile of OmcS are delineated in a manuscript under preparation.

C(E,pH)MD simulations captured much of the same behavior for *E*° along the heme chain as QM/MM@MD, except that (1) the swings in *E*° were more accentuated, as is expected for a fix-point charge electrostatics method,^114^ and (2) the *E*° of **#6** was more, not less negative than **#5**. The latter discrepancy may reflect the difficulty of assigning dielectric boundaries for the implicit solvent used in electrostatic free energy evaluations at the interface of subunits in the filament where **#6** resided.

If a dielectric of ε = 80 (for water) was placed in a protein cavity where *ε* should be much less, the oxidized state of the heme would be more stabilized, and *E*° more negative, than it would be otherwise. Effective static dielectric constants (∈_*eff*_) for the heme binding sites were found to be in the expected 3-7 range for a protein interior. ∈_*eff*_ for the binding sites was determined by comparing *E*°s computed in the presence of either the atomistic and heterogeneous protein environment or various implicit homogeneous solvents (Table S15). At the QM level of theory, this analysis revealed a linear correlation (R^2^ = 0.8673) of more negative *E*° with increasing ∈_*eff*_ (Figure S9), which has been observed previously.^115^

The *E*° for **#6** was 0.158 V more negative at the C(E,pH)MD level than with QM/MM@MD. A similar discrepancy (−0.122 V) was found for **#2**, which had the most solvent exposed macrocycle. The differences on *E*° for the other four hemes came in pairs of similar magnitude and opposite sign (±0.03 V for **#1** and **#4** and ±0.09 V for **#3** and **#5**), a hallmark of stochastic (not systematic) errors. The C(E,pH)MD *vs*. QM/MM@MD differences on *E*° were similar in magnitude to differences previously observed between C(E,pH)MD and experimental data on multi-heme proteins.^101^

Notwithstanding the reasonable agreement, the C(E,pH)MD and QM/MM@MD computations respectively included and excluded heme-heme and redox-Bohr interactions. *E*°s were obtained with C(E,pH)MD by simultaneously titrating the redox and protonation states of all 18 hemes in an OmcS trimer for a range of solution potentials at pH 7. *E*°s with QM/MM@MD were determined for each heme separately without permitting redox state changes on other hemes or protonation state changes on any group. When the classical simulations were repeated with either the propionic acid groups locked in the deprotonated state, or the redox state of all hemes except the one being titrated fixed in the oxidized state, the zigzag pattern of *E*°s along the heme chain was qualitatively the same (Figure S8). Thus, redox and pH cooperativities were not predicted to change the ordering of hemes by *E*° at 300 K and pH 7.

Overall, QM/MM@MD and C(E,pH)MD methodologies gave a largely consistent picture of the *E*°s in OmcS. Heme **#4** was the most readily oxidized and **#1** was (or tied with **#3** as) the most readily reduced heme (Tables S13, S14). Between these extremes that spanned ~0.2 V according to both methods, **#2** was more readily oxidized than **#3**.

The *E*°s at 300 K in the present study, and 310 K in a prior work,^83^ both obtained with QM/MM@MD techniques (see Table S1 for an extensive comparison), agreed within |0.072| V for almost every heme (Figure 2, orange on the *left* vs. red on the *right*). The exception was **#5** for which *E*° was too far negative by 0.3 V in the prior work. This level of agreement attested to the robustness of the computed *E*°s in both works; it also indicated that the discrepancy for **#5** was not a systematic error due to different choices in the classical forcefield or approximate density functional. The difference was also likely not due to the 10 K warmer temperature used for MD in the prior work. That supposition would require an average temperature coefficient of 0.03 V/K, which is 10-times larger than experimental values for a variety of cytochromes.^86–92^ ^88, 91, 93^ Given that the *E*° for **#5** in the prior study^83^ was at least ~0.15 V outside the expected experimental range,^94^ and disagreed with both C(E,pH)MD and more rigorous QM/MM@MD computations by an amount at least three-times larger than for any other heme, the data suggests that the *E*° reported here for **#5** is a better estimate of the true value.

The revised *E*° for **#5** significantly reduced the redox potential differences and thereby free energy changes for electron transfer to and from this heme on which Marcus theory rates for heme-to-heme electron transfer exponentially depend. These Marcus rates were previously used to evaluate the charge mobility with Kinetic Monte Carlo (KMC),^83^ The implications of the more positive *E*° for **#5** on charge mobility are discussed in §2.4.

#### 2.2.2. Low (270 K) Temperature

Lowering the temperature by 30 K induced negative and nearly uniform (−0.024 – −0.101 V) shifts in *E*° for each of the six hemes (Figure 2, orange vs. blue on the *left*; Table S13). The average computed temperature coefficient of −0.002 V/K was quantitatively consistent with temperature coefficients experimentally determined for a variety of other cytochormes.^86–92^ ^88, 91, 93^

CEMD simulations found a negligible (~0.023 V) average shift in *E*° upon cooling (Table S14). This result must be treated with care, however, because it assumes—based on some experiments^116^—that the *E*° of the *N*-acetylmicroperoxidase-11 reference compound for the simulations was temperature independent. The result at least suggests that there was no dramatic structural transition within the protein from 300 to 270 K since the electrostatic free energies for heme oxidation were similar at both temperatures. Indeed, the analysis in §2.3.2 below suggests that a change in solvent density was primarily responsible for the largest shifts in *E*° upon cooling, an effect that would be missed by CEMD because the energy evaluations in this approach are performed with an unstructured implicit solvent around the protein.

Because of the uniformity of the cooling-induced shifts at the QM/MM@MD level, the rank ordering of the hemes from most to least negative *E*° at both 300 and 270 K was similar: 4 < 5 < X < Y < Z < 3, where X, Y, and Z = 2, 6, 1 at 300 K and 6, 1, 2 at 270 K. Importantly, the “<“ corresponded to an average increment of 0.035 V, comparable to the standard error of the mean for each *E*°. Thus, the cooling-induced re-ordering of the hemes by *E*° was not statistically meaningful. This is evidenced in Figure 2 *left* by the overall similar zigzag pattern of the *E*°s along the heme **#1** → **#2** … **#6** chain.

These observations regarding shifts in *E*° upon cooling were substantially different from the work by Dahl *et al* (Figure 2, *left* vs. *right*).^83^ The reported rank ordering of the hemes from most to least negative *E*° in that work was **#5** < X < **#1** < Y < **#2** < **#6**, where X and Y were respectively **#4, #3** at 310 K and **#3, #4** at 270 K. The “<“ corresponded to a spacing in *E*° twice as large (0.081 V) as found in the present study. Hemes **#3** and **#4** exchanged positions in the series upon cooling because of nearly equal and opposite shifts in *E*° (−0.189 and +0.113 V) for these two hemes in van der Waals contact. This prior study therefore reported a range of temperature coefficients that was unexpectedly large and of mixed sign (+0.004 to −0.006 V/K). The implications of the differently sized and signed shifts in *E*° found in the present and prior work for charge mobility are analyzed in §2.4

### 2.3. Microscopic origins of Cooling-Induced Redox Potential Shifts

#### 2.3.1. Solvent and Protein electrostatics

An instructive way to look at the temperature dependence of *E*° was in terms of which hemes were most sensitive to the change in this thermodynamic variable. The hemes with the most to least responsive *E*° upon cooling 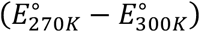 were **#5** > **#6** > **#1** > **#4** > **#3** > **#2**. This ordering was almost perfectly reproduced (Figure 3; Table S13, S16) by the change in electrostatic interaction energy for oxidation of the hemes at 270 *versus* 300 K (Δ*E*_270*K*−300*K*_) (Eq. 1). The only exception was that the ordering of **#1** and **#4**, which were in the middle of the series and had very similar shifts, was inverted for Δ*E*_270*K*−300*K*_.

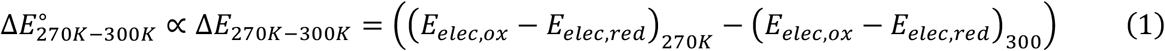

where *E*_*elec,ox*/*red*_ is the electrostatic interaction energy for a given heme in the oxidized/reduced state. Δ*E*_270*K*−300*K*_ was always larger than 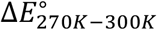 by a factor of 1.4 to 4.2, which likely reflected the greater sensitivity of a fixed point-charge electrostatics model to instantaneous environmental conformations.^114^

**Figure 3.**
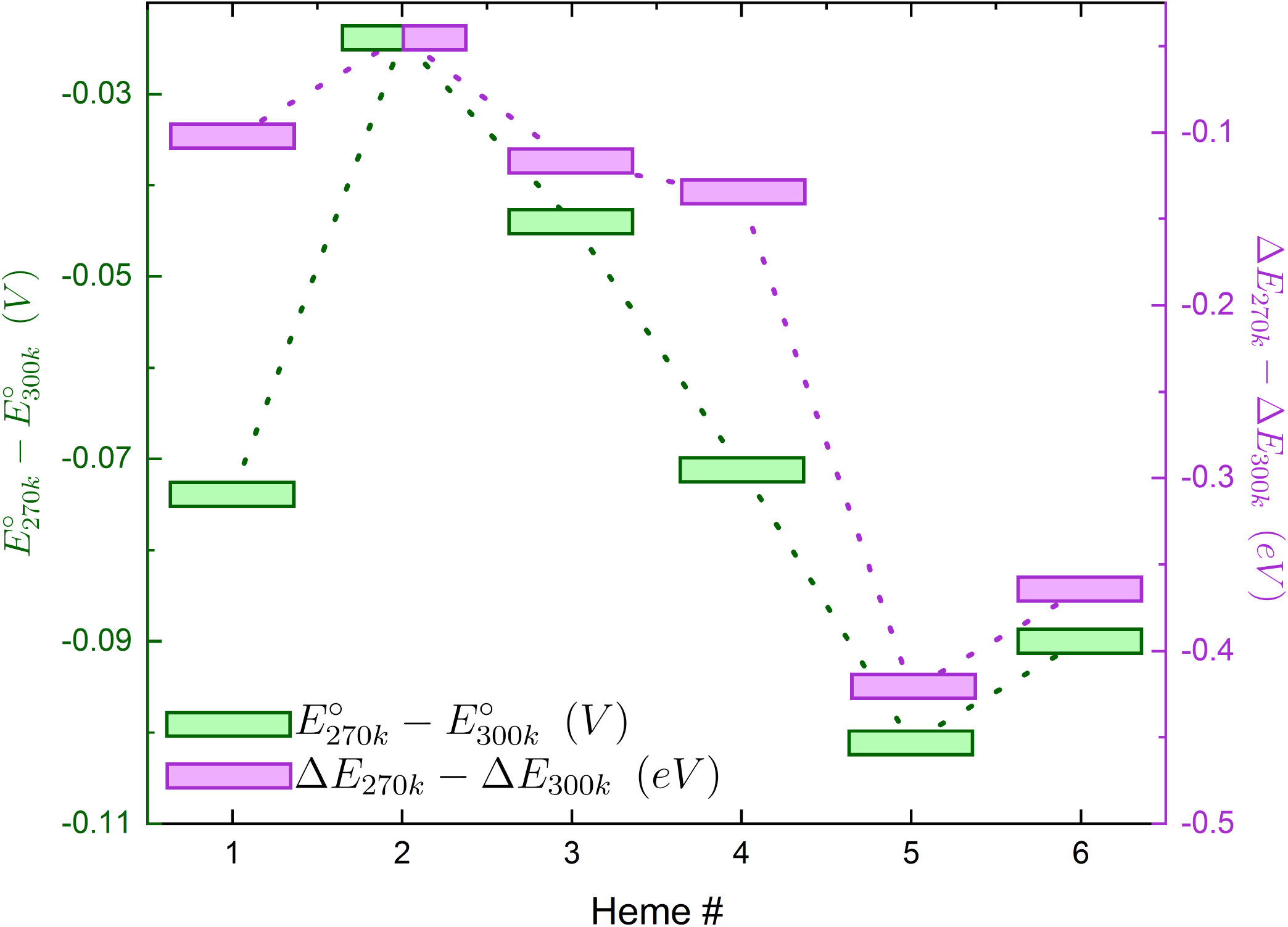
Cooling-induced shifts in redox potential 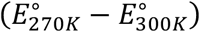 were well matched with shifts in redox-linked electrostatic interaction energies (Δ*E*_270*K*_− Δ*E*_300*K*_), where the Δ*E* at a given temperature for each heme is computed as *E_elec,ox_* − *E_elec,red_*.

Because Coulombic interactions are pairwise decomposable, ΔE_270*K*−300*K*_ was partitioned into contributions from the solvent or the protein. The latter contribution was further divided into influences from non-polar, aromatic, acidic, and basic residues, as well as other heme groups (Figure 4, Table S16). This analysis revealed that the solvent made a dominant contribution to the cooling-induced shifts in electrostatic potential energy (Figure 4), and thereby redox potential (Figure 3) for the most temperature-sensitive hemes (**#5** and **#6**).

**Figure 4.**
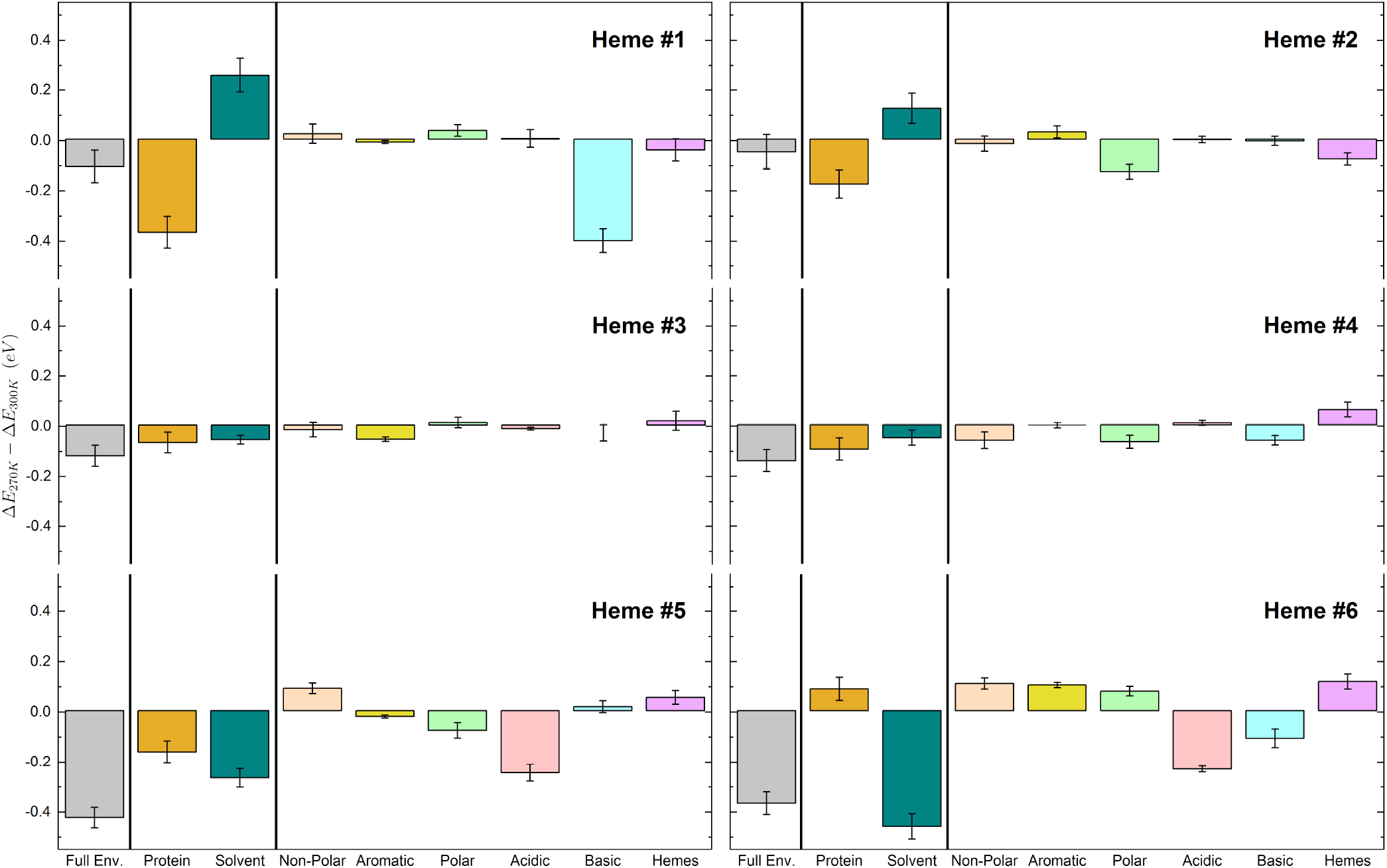
Decomposition of cooling-induced shifts in redox-linked electrostatic interaction energies into contributions from the solvent, protein, or various physicochemical groupings of residues.

The negative-going shift in Δ*E*_270*K*−300*K*_ for heme **#5** resulted from a 62:38 ratio of solvent and protein contributions. The dominant solvent contribution was associated with an increase in the radial distribution function, particularly at distances < 4 Å with respect to heme **#5** in the oxidized state (Figure S10). The smaller protein contribution mainly originated from acidic groups and to a lesser extent polar and aromatic residues (Figure 4). Opposing electrostatic contributions were exerted by non-polar residues, other heme groups, and basic residues.

In the case of heme **#6**, the strong negative-going shift in Δ*E*_270*K*−300*K*_ from the solvent was very weakly opposed by a positive-going contribution from the protein (Figure 4). Again, the solvent contribution was associated with an increase in the radial distribution function, particularly at distances > 6 Å, with respect to heme **#6** in the oxidized state (Figure S10). The protein contribution reflected, as for the other heme, the net result of counter-balancing factors from different groups of residues.

To summarize, all hemes experienced a negative-going shift in *E*° upon cooling that correlated with more negative electrostatic energies for heme oxidation (Figure 3). The largest shifts were experienced by **#5** and **#6**, and primarily originated from altered electrostatics with the solvent (Figure 4). These altered interactions were likely related to an increase in density of the hydrating water layers within a 10 Å radius of **#5** and **#6** (Figure S10). Experimentally, the density of water (above the normal freezing point), and static dielectric constant^117^ increase upon cooling. Increases in the static dielectric constant were found above (§2.2.1) to be linearly correlated with more negative *E*° (Figure S9). Thus, the increased density and static dielectric constant of water upon cooling explain the shift to more negative redox potentials for the most temperature sensitive hemes in OmcS.

Figure 3 directly related 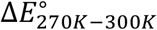 computed with QM/MM@MD to Δ*E*_270*K*−300*K*_ computed with classical electrostatics. Figure 4 followed-up on this analysis to provide a microscopic explanation for the distinct temperature sensitivities of the hemes. Both the approaches and insights drawn from these figures mark an advance over prior work. Previously, 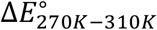 was only weakly correlated (R^2^ = 0.51-0.65) with changes in the planarity of the heme macrocycles or the electric field exerted on the Fe centers.^83^ Both effects were attributed to a cooling-induced restructuring of the H-bonding network; a phenomenon examined in the next subsection.

#### 2.3.2. Intra-protein H-bonding networks

Inspired by the proposed temperature sensitivity of H-bonding in OmcS and its controlling role over the electronic structure of the hemes,^83^ the number, persistence, and composition of H-bonds as a function of temperature from 100 to 400 K were analyzed.

Over this temperature range, the number of H-bonds varied linearly (R^2^ = 0.9778) with temperature according to the equation (Table S17): # of H − bounds = 420(±6) − 0.50 (±0.02)T. This behavior was reflected in a linear dependence (R^2^ = 0.9969) of the characteristic H-bonding frequency (CHF; Figure 5, *top*; Table S18): CHF = 19.1(±0.1) − 0.0273(±0.0004)T. The CHF is a unitless quantity defined as the norm of a matrix of per-residue donor-acceptor H-bonding occupancies. This result was in good agreement with the decrease in the CHF by 1.0 previously reported for OmcS from 270 to 310 K.^83^ The analysis in Figure 5 adds that the temperature dependence of the CHF is linear—at least for OmcS—and extends over a much wider temperature range.

**Figure 5.**
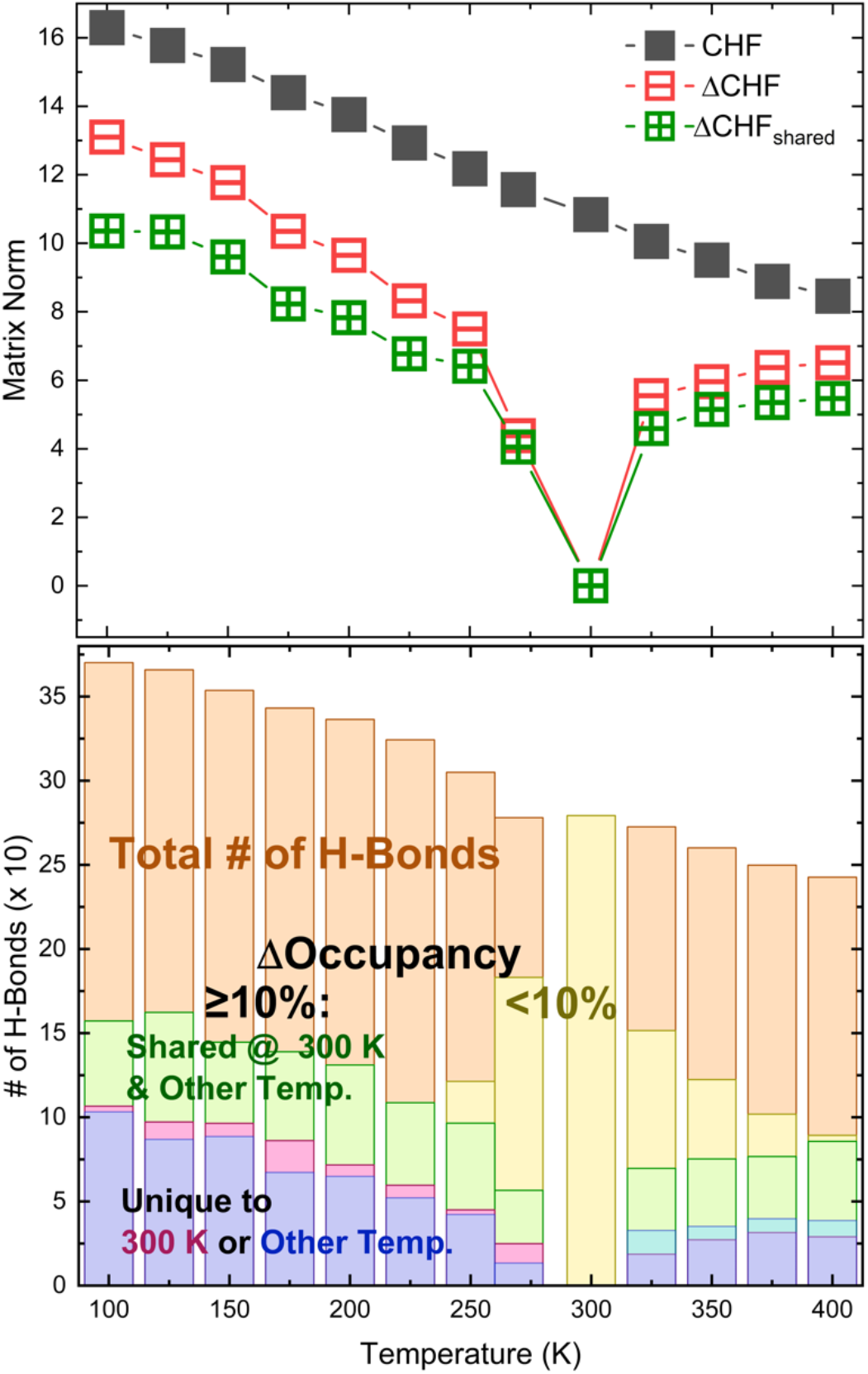
(*top*) Variation with temperature in the characteristic H-bonding frequency (CHF), the difference (ΔCHF), and the ΔCHF only considering H-bonds present at 300 K and some other temperature (ΔCHF_shared_). See text for the definitions of these terms. (*bottom*) Change in the number and occupancy of intra-protein H-bonds as a function of temperature. Both the top and bottom panels are on a common x-axis scale.

To complement the analysis of the CHF as a function of temperature, the difference CHF (ΔCHF) was also computed. ΔCHF is a unitless quantity defined as the norm of a matrix of differences in per-residue donor-acceptor H-bonding occupancies between two temperatures (Figure 5, top; Table S18). The ΔCHF was computed at each temperature relative to 300 K, where it therefore had a value of zero (no difference). Interestingly, the ΔCHF and the ΔCHF:CHF ratio increased as the temperature either decreased or increased from 300 K (Figure 5, *top*). Large (positive or negative) changes in occupancy at cooler or warmer temperatures relative to 300 K both produced large ΔCHFs because these differences were squared when computing the norm of the difference matrix.

A large ΔCHF could reflect a significant change in the connectivity of the H-bonding network if many H-bonds only existed at one of the two temperatures being compared. However, the norm of the difference matrix filtered to only contain elements for H-bonds present at both temperatures (ΔCHF_shared_) nearly equaled the full ΔCHF at a given temperature (Figure 5, *top*; Table S18). This result suggested that H-bonding interactions present at 300 K were a dominant contributor to the ΔCHF at other temperatures. In fact (Figure 5, *bottom*; Table S17), >81% of H-bonds present at 270 and 325 K either also existed at 300 K, or had a change in occupancy (including from zero) < 10% relative to the value at 300 K. Fewer than 12 H-bonds that had a change in occupancy > 10% within the 270-325 K range were within 10 Å of the central six hemes in the filament for which *E*°s were computed (Tables S19-S21). These findings were consistent with the prior result that only 7% of H-bonding interactions were unique to the 270 *versus* 310 K simulation.^83^ Thus, the results of the current study suggest that the intra-protein H-bonding network was highly similar in the 270-325 K range, particularly in the vicinity of the hemes. Other factors apart from, or in addition to changes in H-bonding (Figures 3, 4) likely contributed to the temperature dependence of the redox potentials and electrical conductivity in OmcS.

### 2.4. Temperature Dependence of Redox Conductivity

Roughly ~10^5^ electrons/s flow under physiological conditions out *of a Geobacter sulfurreducens* cell.^118^ The OmcS filament is proposed to carry some fraction of this current. OmcS resembles other redox chains in biology—albeit on a much longer (μm instead of nm) length scale—in that the filament connects discrete electron donors (unidentified) and acceptors (e.g., Fe(III) or Mn(IV) oxide nanoparticles). In-between the primary donor and ultimate acceptor, electrons are thought to non-adiabatically tunnel from heme-to-heme in a multistep process under physiological (not necessarily solid-state) conditions.^119–123^

This “hopping” regime of electron transfer is typically well-described by Marcus theory.^124, 125^ As shown in Eq. 2, the Marcus rate for each electron transfer step (*k*_*nm*_ for donor *m* → acceptor *n*) depends on the free energy change for the reaction (Δ*G*_*mn*_), the reorganization energy for the polarization response of the nuclei (λ_mn_), and the electronic coupling between the electron donor and acceptor (*H*_*mn*_). Note that the convention is to specify the donor and acceptor from right-to-left for the rate constant, but left-to-right for the energetic terms. *k*_7_, *T*, ℏ, and ⟨… ⟩ in Eq. 2 respectively signify the Boltzmann constant, absolute temperature, the reduced Plank constant, and thermal averaging. Δ*G*_*mn*_ in Eq. 2 was determined from the difference in *E*° between the donor and acceptor. Intensive focus was given above to obtain high-quality estimates of *E*° because each *k*_*nm*_ exponentially depends on Δ*G*_*mn*_.

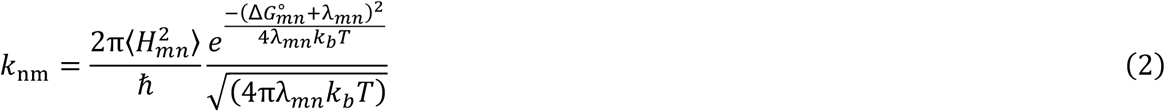

Using the best estimates of *E*° for the hemes in OmcS to determine Δ*G*_*mn*_ (Table S22), along with computed λ_mn_ (Table S23, S24) and ⟨H_mn_⟩ (Table S25) values (see §S1.3-S1.5 for details), Marcus rates were evaluated in the forward (**#1** → **#2** … → **#6**) and reverse (**#6** → **#5** … → **#1**) directions (Table S26). These rates were assembled into kinetic models to simulate the electron current (Tables S27, S28). Similar modeling efforts have already been described for OmcS.^82–84^ To aid comparison with these works, the multi-particle steady-state flux model of Blumberger and co-workers,^126^ as well as the single-particle diffusion model employed by Amdursky and co-workers^84^ and Malvankar and co-workers^83^ were both used in the present study.

To compare the diffusive and flux models with one another as well as experiment, the following procedure was devised (see §S1.6.3 in the SI): The experimentally measured resistance,^83^ and the resistance expressed in terms of the computed charge diffusion constant (Eq. S10 in the SI) were each related to an electrical current at a given applied bias using Ohm’s law. The voltage was simply a multiplicative scaling factor for this experiment *versus* theory comparison, and the conclusion was independent of the chosen voltage (within the ohmic *I*-*V* regime for OmcS). The voltage was chosen to correspond to the electron injection/ejection rate used in the flux model to obtain a protein-limited current. This choice allowed a comparison of the experimental current with both the simulated diffusive and protein-limited currents at the same voltage.

Importantly, the voltage needed to attain the protein-limited current (Figure S7) was 10-times larger (1.1 V) than the voltage used in electrical measurements on OmcS. For this reason, the current computed at this voltage with the experimentally measured resistance is referred to as hypothetical. The currents measured at lower voltages are presumably limited by the protein-electrode contacts and do not reflect the intrinsic conductivity of the filament.^127^ And yet, as discussed below, the protein-limited current severely underestimated both the hypothetical current at the same voltage, as well as measured currents at much lower voltages. The diffusive current was also orders of magnitude too small.

In the following Δ*G*_*mn*_, λ_mn_, ⟨H_mn_⟩, and *k*_*nm*_ are first compared across all available hopping models for OmcS, including the present study (§2.4.1-2.4.3; Tables S22-S26). Simulated currents obtained by parameterizing the diffusive and flux kinetic models with *k*_*nm*_s from all these studies are then compared (§2.4.4; Tables S27, S28).

#### 2.4.1. Reaction Free Energy (ΔG_mn_)

Figure 6 compares the free energy landscapes computed in the present and prior studies.^82, 83^ ΔG_mn_s reported here were −0.128 to +0.190 eV at 270 or 300 K. This range was considerably smaller than the −0.441 to +0.208 eV reported in the prior study^83^ at 270 or 310 K, but in very close agreement with the work of Jiang *et al*.^82^ at 300 K (−0.090 to +0.120 eV). The larger range found before primarily resulted from a too-far negative *E*° for heme **#5** at 310 K in that work^83^ (as discussed in §2.2 above).

**Figure 6.**
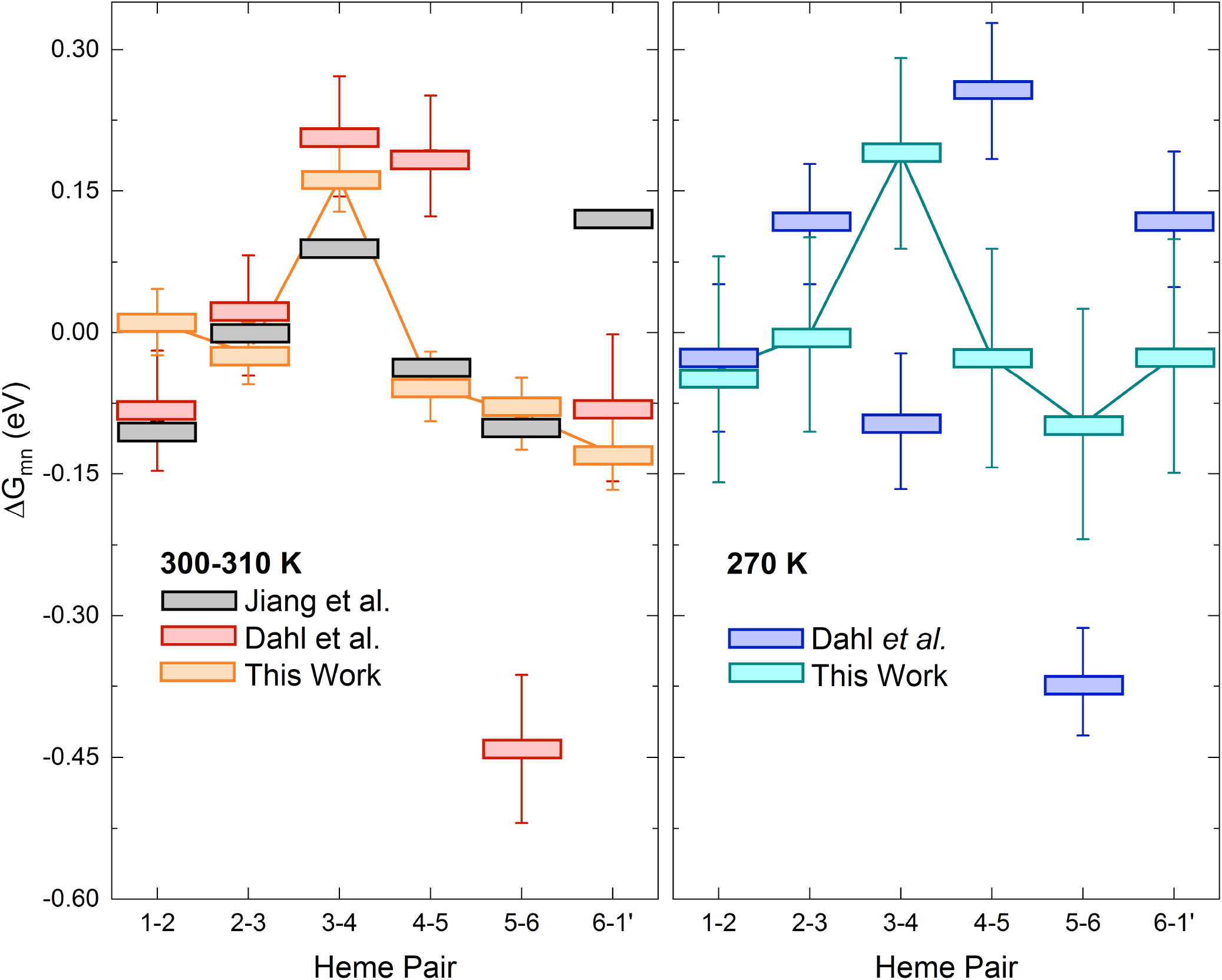
Free energy landscapes at (*left*) high (300 or 310 K) and (*right*) low (270 K) temperatures. At high temperature, the results of the present study and that by Jiang *et al*. pertain to 300 K; the work by Dahl *et al*. pertains to 310 K.

The individual ΔG_mn_s at 300 K in the current study generally differed by ≤ 0.1 eV with respect to all prior work at high temperature (Figure 6, *left*), which is quite reasonable given uncertainties from sampling MD trajectories and systematic errors between different approximate density functionals. The only exception with respect to the work of Jiang *et al*.^82^ was for Δ*G*_6,1_ (|0.25| eV). In that work, there may have been difficulties in assigning dielectric boundaries for the implicit solvent at the interface of subunits where hemes **#6** and **#1** reside. A similar issue was already noted is §2.2 when comparing *E*°s from C(E,pH)MD simulations, which use an implicit solvent for electrostatic free energy evaluations, and explicit solvent QM/MM@MD. Larger deviations were found at high temperature with respect to the work of Dahl *et al*. ^83^ for Δ*G*_4,5_ (|0.25| eV) and Δ*G*_5,6_ (|0.35| eV), likely because the *E*° for heme **#5** was too far negative by 0.3 V in that work.

At 270 K, the ΔG_mn_s in the present study deviated from the report of Dahl et al.^83^ by ~|0.13| eV for Δ*G*_2,3_ and Δ*G*_6,1_, and ~|0.28| eV for Δ*G*_3,4_, Δ*G*_4,5_, and Δ*G*_5,6_. The origin of these large differences is not entirely clear. However, for reference, the mean-unsigned-error of the Δ*G*_*m,n*_ s in the present study was half as large as for Dahl *et al*.^83^ at high temperature relative to the work of Jiang *et al*.^82^

#### 2.4.2. Reaction Reorganization Energy (λ_mn_)

λ_mn_ for each electron transfer step in OmcS, regardless of temperature, was found to be within the expected ~0.5 to 1.0 eV range for a fully solvated protein in the present and prior studies.^82, 83^ Each λ_mn_ reported here agreed within a factor of 0.7 to 1.4 of values in the literature for OmcS.^82, 83^

To consider the impact of some unquantified amount of desolvation for the electrical measurements, λ_mn_ was re-computed after removing either all water molecules and Na^+^ ions, or only those beyond the first solvation shell (≤ 3.4 Å from the filament). λ_mn_—more specifically, the Stokes reorganization energy—decreased by 35-58% in these analyses; meanwhile the variance reorganization energy tended to increase (Table S24). (See §S1.4 in the SI for the definitions of these various types of reorganization energy.) The divergence between Stokes and variance reorganization energies is a hallmark of non-ergodicity. Ergodicity-breaking is one of the possible explanations for the *anti*-Arrhenius temperature dependence of conductivity in OmcS (discussed in §3.4).

#### 2.4.3. Thermally averaged Donor-Acceptor Electronic Coupling (⟨H_mn_⟩)

The ⟨H_mn_⟩ for each electron transfer step was < 0.02 eV in the present and prior studies (Table S25).^82, 83^ The finding that ⟨*H*_*mn*_⟩ ≪ λ_mn_ justified the use of Marcus theory in the non-adiabatic limit (Eq. 2) to model redox conductivity.

Each ⟨H_mn_⟩ agreed within a factor of ≤ 2.0 of those reported by Jiang *et al*.^*82*^ at 300 K, and ≤ 8.0 of those reported by Dahl *et al*.^83^ at 310 K. A similar level of agreement was found for the ⟨H_mn_⟩s at 270 K with the work of Dahl *et al*. at the same temperature..^83^ A negligible (~0.001 eV) change in ⟨H_mn_⟩ was observed upon cooling from 300 to 270 K. Consistent with prior work,^128^ the ⟨H_mn_⟩s were only minimally changed by the inclusion of an electrostatic point-charge environment, or the propionic acid substituents of the hemes in the QM region of the calculations (Table S11).

#### 2.4.4. Marcus Electron Transfer Rates (k_nm_)

Forward (**#1 → #2 … → #6**) and reverse (**#6 → #5** … **→ #1**) rates at 300 K ranged from 2 × 10^6^ to 2 × 10^10^ s^-1^, and generally matched those reported by either Jiang *et al*.^82^ at 300 K or Dahl *et al*.^83^ at 310 K within one order of magnitude (Figure 7, *top*). Relative to Jiang *et al*., only *k*_6,5_ and *k*_3,4_ differed by two orders of magnitude. Relative to Dahl *et al*. there were differences of 2 (*k*_5,4_, *k*_6,5_, *k*_1,2_), 3 (*k*_5,6_), and 4 (*k*_2,1_) orders of magnitude. These differences are not the result of the 10 K warmer temperature used in the simulations by Dahl *et al*.

**Figure 7.**
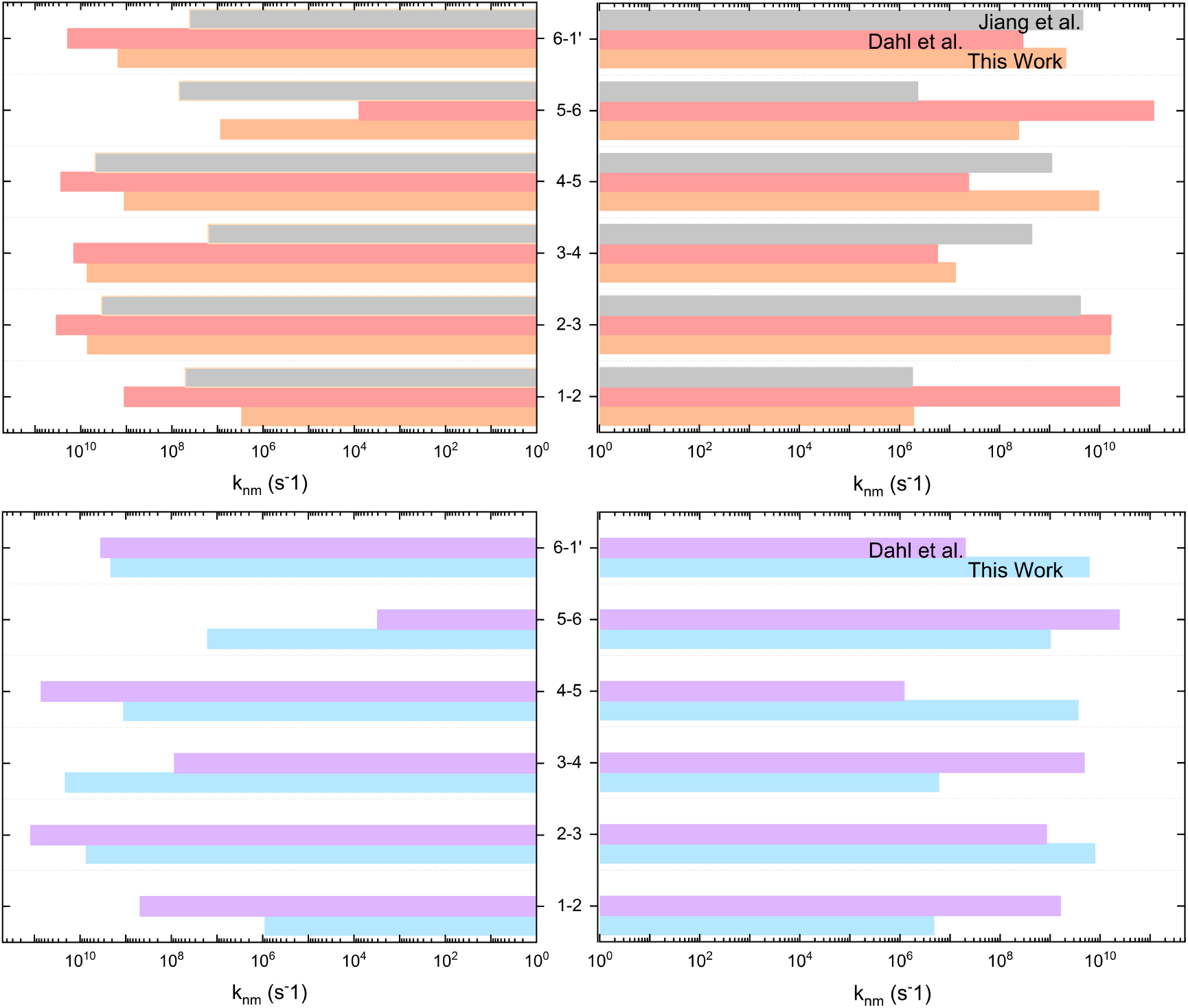
Forward (*right*) and backward (*left*) rates at (*top*) high (300-310 K) and (*bottom*) Low (270 K) temperatures.

At 270 K (Figure 7, *bottom*), the rates presented here also generally agreed with the work of Dahl et al.^83^ within an order of magnitude. Differences of 2 (*k*_2,1_, *k*_1,2_, *k*_4,3_, *k*_3,4_, *k*_1′,6_) and 3 (*k*_5,4_, *k*_5,6_) orders of magnitudes also occurred for some rates.

#### 2.4.5. Simulated Electrical Hopping Currents

Figure 8 shows the electrical currents computed with a single-particle diffusion and a steady-state flux kinetics model parameterized with Marcus rates from Jiang *et al*.^*82*^, Dahl *et al*.,^83^ and the present work.

**Figure 8.**
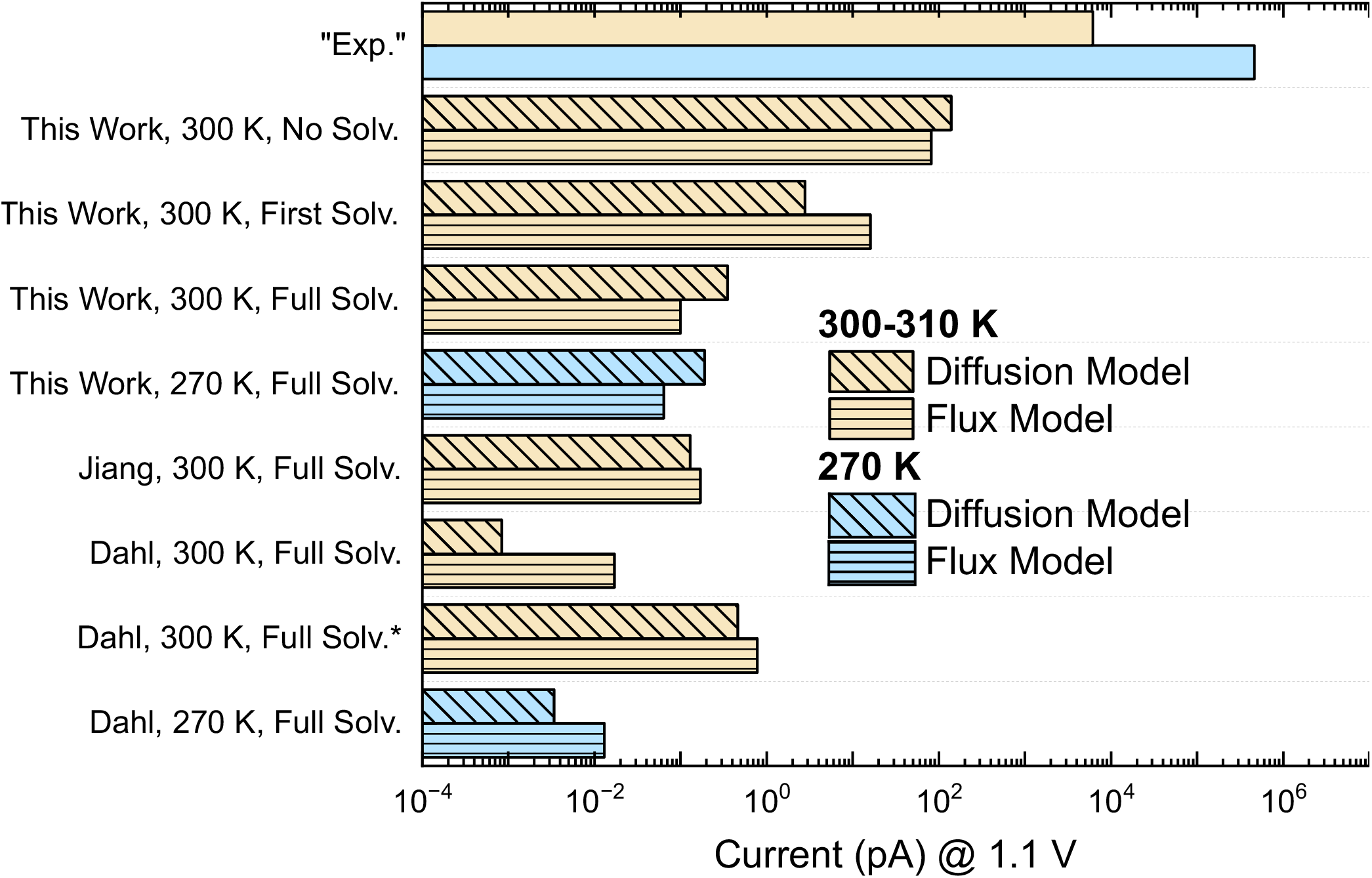
Comparison of “experimental” and simulated hopping currents. The “experimental” (with quotation marks) indicates the current that would be observed based on the measured conductivity of the OmcS filament if it could withstand and maintain Ohmic behavior under a 1.1 V bias. The simulated currents were modeled with a single particle diffusion (diagonal lines) and a multi-particle steady-state flux (horizonal lines) model. “No Solv.”, “First Solv.” and “Full Solv.” Indicate that the contribution of waters/ions to the reorganization energies were entirely neglected, only included for the ~3400 water molecules and ~12 Na^+^ ions within the first solvation shell (3.4 Å of the filament), or the full complement of waters/ions in the simulations. The entry marked with an asterisk (*) indicates that the redox potential for heme **#5** was shifted by +0.3 V to be more consistent with the revised value reported in the present work.

An immediate observation from Figure 8 is that the theoretical hopping currents— regardless of which set of Marcus rates or which kinetic scheme was used— underestimated the hypothetical experimental (“Exp.”) current. All models including the reorganization energy contributed by the aqueous solvent in the fully hydrated state predicted currents that were 3- to 6-fold too small. When none of the reorganization energy contributed from the aqueous solvent, or only the portion due to the first solvation shell (~3400 waters and 12 Na^+^ ions within 3.4 Å of the filament) was included, the predicted currents were 2- to 3-fold closer to the hypothetical current. This result was sensible because the activation energy for each electron transfer ((Δ*G*_mn_ + λ_mn_)^2^/4λ_mn_) in solution-phase simulations of OmcS was dominated by λ_mn_ (λ_*mn*_ ≫ Δ*G*_*mn*_), ~50% of which originated from the solvent (Table S24). The amount of solvent present in the experiments is unknown and may play an important role in the temperature dependence.^129^ At the very least, this result suggests that simulations need to consider the conditions of solid-state electrical measurements in detail before reaching a verdict on the operative charge transport mechanism.

Other Important observations from Figure 8 were: (1) Simulated hopping currents at 300 K in the present study agreed (within a factor of 3) of the work by Jiang *et al*.^*82*^; (2) The currents at 310 K for the work by Dahl *et al*.^*83*^ were one- to two-orders of magnitude smaller than the results presented here; (3) This discrepancy was entirely removed by shifting the previous *E*° for **#5** by +0.3 V, which seemed warranted from the results in §2.2.1; (4) From 310 to 270 K, the currents for Dahl *et al*.^*83*^ decreased by 25% (flux model) or increased by a factor of 4 (diffusion model) depending on the kinetic scheme; (5) From 310 to 270 K, a 60 – 130-times decrease in the current for either kinetics scheme was predicted for Dahl *et al*.^*83*^ with the revised value of *E*° for **#5** at 310 K. Note that shifting the *E*° for any other heme at 310 or 270 K was not warranted, because all the *E*°s agreed with the present work within |0.16| eV, a deviation half as large as for **#5** at 310 K. Even if the *E*° of **#5** was only shifted by +0.15 V, the current at 270 K vs. 310 K for Dahl *et al*. was found to be 10 – 16-times smaller (Table S28). Thus, Figure 8 suggests that the previously simulated increase in conductivity with decreasing temperature may have resulted, in part, from an underestimation of the conductivity at high temperature because of a too-far negative E° for heme **#5**.

Irrespective of this point, the Marcus rates reported by Dahl *et al*.^*83*^ taken at face-value, do not give the previously reported 77-times increase using two different kinetic schemes, one of which is an analytical approach to the diffusion problem that was previously treated with KMC. A reason for this discrepancy is given in §3.3.

## 3. Discussion

### 3.1. What charge transport mechanism is operative in OmcS?

It is currently unclear whether incoherent charge hopping can account for the conductivity of OmcS,^82, 83^ or if coherence-assisted hopping^84^ or decoherent quantum transport^85^ mechanisms need to be invoked. Incoherent hopping was consistent with a measured linear-dependence of conductance on filament length,^83^ but this is not an unambiguous metric.^130^ Temperature is another factor that is usually varied to diagnose a particular charge transport mechanism. But the complex dependence found for OmcS (Figure 1, *bottom*) does not conform neatly to Landauer-Buttiker (coherent tunneling) or Marcus (incoherent tunneling) theory, or a generalization of both theoretical frameworks.^131, 132^

In the present study, redox conduction through a freely floating and isolated OmcS oligomer in electrolytic solution severely underestimated the experimental solid-state electrical conductivity (Figure 8). This conclusion was independent of whether Marcus rate constants reported by Jiang *et al*.^*82*^ or Dahl *et al*.^*83*^ were substituted for those computed here when parametrizing both diffusive and steady-state flux kinetic models.

The discrepancy, which was already foreseen by Amdursky and co-workers,^84^ may reflect the inadequacy of the incoherent hopping mechanism. These authors proposed that either the reorganization energy accompanying electron transfer was unusually low for biological systems (~0.15 eV), or that rigidification of the protein scaffold preserved coherences among blocks of heme groups to assist the incoherent charge hopping process. Since loops and turns comprise 81% of the 2° structure of OmcS,^133^ and the inter-heme electronic coupling is less than thermal energy at 298 K (Table S27), it is difficult to envision how coherence over a 20 nm stretch of hemes, as proposed,^84^ could be maintained.

The other (unusually low reorganization energy) hypothesis may be realized in the partially dehydrated state of OmcS under experimental conditions. Under this hypothesis, the incoherent hopping mechanism appears insufficient only because the physical state of the system was inadequately modeled. Beratan and co-workers,^134^ noted that dehydration of the protein nanowire likely removes a significant fraction^135^—perhaps as much as 50%^136^--of the outer-sphere reorganization energy.^135^ Indeed, the activation energies for electron transfer in solution-phase simulations of OmcS were dominated by λ_*mn*_, a large fraction of which came from the aqueous solvent (Table S26). Simulated hopping currents reported in Figure 8 became a few orders of magnitude closer to the hypothetical experimental value when most or all the reorganization energy contributed by the aqueous solvent was neglected.

The level of hydration has not always been rigorously controlled in temperature-dependent conductivity measurements on pili/OmcS. Variation in experimental conditions, including hydration, have been implicated in the observation of different temperature dependences.^129, 137, 138^ The bell-shaped pattern reported by Dahl *et al*.^83^ (Figure 1, *bottom*) was reported earlier—albeit ascribed to pili at the time—under similar conditions.^36^ At temperatures above 270 K, however, a much weaker increase^139^ or even a decrease^140^ in conductivity upon cooling was reported under different conditions. The moisture/water content was shown in the context of *G. sulfurreducens* biofilms to determine whether an increase or decrease in conductivity was observed with decreasing temperature.^129^ In kinetic isotope experiments at different temperatures,^83^ a sharp change in conductance at 220 K was ascribed to the dynamical protein transition, which is intimately related to the hydration shell.^141^

Better experimental characterization is needed because deficiencies in modeling conductivity may result just as much from missing key structural aspects as applying the wrong charge transport theory. At the same time, more attention in modeling these experimental details is needed because simulations on a fully hydrated and freely floating protein oligomer in electrolytic solution are likely of little relevance for modeling charge transport through an electrode-adsorbed, (partially or fully) dehydrated, and (possibly) aggregated protein.

### 3.2. Is there a temperature-sensitive ‘switch’ for H-bonding in OmcS that controls electrical conductivity?

Cooling-induced restructuring of intra-protein H-bonds was previously attributed with shifting the *E*°s of the hemes and thereby producing the *anti*-Arrhenius conductivity behavior. The restructuring was quantified in terms of the norm of a H-bonding occupancy matrix (dubbed the Characteristic H-bonding Frequency, CHF) at a given temperature, and the norm of a difference matrix (dubbed ΔCHF) between two temperatures.

From 300 to 270 K, where OmcS filaments exhibit *anti*-Arrhenius conductivity, the CHF and ΔCHF changed respectively by 0.8 and 4.4 in the present study. Similar changes of 1.0 and 6.2 were found in the prior work between 310 and 270 K.^83^ In fact, a linear dependence of the CHF on temperature was found (§2.3.2) over the entire 100 to 400 K range. This analysis suggested that temperature was more like a ‘dial’ than a ‘switch’ for the overall occupancy of the H-bonding network. However, a straightforward connection between the monotonic and bell-shaped dependencies of H-bonding and electrical connectivity on temperature was not obvious.

The larger magnitude of ΔCHF compared to the change in CHFs between two temperatures (i.e., the norm of a difference matrix *versus* the difference in norms of individual matrices) was interpreted in the prior work as a “massive restructuring” of the H-bonding network.^83^ This terminology, however, is potentially misleading. Both the prior and present studies found < 10% of H-bonds to be unique to either high (300 or 310 K) or low (270 K) temperature simulations.

In the simulations reported here, 66% of H-bonds had a < 10% change in occupancy (including from zero), and another 20% of H-bonds were common to both temperatures but experienced larger changes in occupancy (Table S17). Thus, the data suggested that the connectivity of the H-bonding network was largely intact, not restructured, between 300 and 270 K.

Only 4% and 8% of the remaining H-bonds were unique to 270 or 300 K, respectively, and had an occupancy > 10%. Of these unique H-bonds, fewer than a dozen occurred within 10 Å of the hemes at the center of the filament for which *E*°s were computed. Given this observation, it seems unlikely that restructuring in H-bonds, taken alone, can account for significant shifts in the *E*°*s* of the hemes upon cooling.

With virtually identical metrics for the change in H-bonding upon cooling as found in the present work, the prior study^83^ reported a maximal change in *E*° that was twice as large as found here, and one-fourth the entire biological range of heme redox potentials.^94^ This comparison suggests that changes in intra-protein H-bonding were not the controlling factor for the temperature sensitivity of the *E*°s. Other factors mentioned in the prior study—changes in heme macrocycle planarity or the electric field exerted on the Fe centers of the hemes—were only weakly correlated (R^2^ = ~0.6) with the change in Δ*E*°, and proposed to be caused by H-bond restructuring.^83^

A different interpretation of Δ*E*° emerged in the present work from Figures 3 and 4 (§2.3.1). These analyses demonstrated that cooling-induced changes in electrostatic energy for heme oxidation and the redox potentials of the hemes were qualitatively matched. By partitioning the change in electrostatic interactions into contributions from various groups, the solvent was found to play a dominant role in the negative-going shift in electrostatic energy, and thereby redox potential for the most temperature-sensitive hemes. The prior analysis^83^ did not consider the influence of the solvent.

Thus, the present results suggest: (1) Only modest restructuring of the H-bonding network, particularly near the hemes, occurs over the 30 K range of *anti*-Arrhenius conductivity. (2) Identical metrics for changes in H-bonding (e.g., CHF and ΔCHF) can occur in simulations that find differently signed and sized shifts in *E*°, suggesting other factors have a more commanding role over the temperature dependence of this property. (3) Changes in electrostatic interactions, particularly with the solvent as its structure changes with cooling, may play a dominant role in the temperature dependence of *E*°.

Findings (1) and (2) further suggest that an interpretation different from H-bond restructuring may apply to a previously reported kinetic isotopic effect (KIE).^83^ A 3- to 300-fold reduction in conductivity was reported upon deuteration of films of OmcS filaments. KIEs ≥ 130 for OmcS in the 230 – 260 K range were reminiscent of those ascribed to concerted electron-proton transfer in wild-type soybean lipoxygenase (KIE = 81), ^23^ and proton coupled electron transfer (PCET) in a benzoquinone and Os(IV)-hydrazido complex (KIE = 455). ^24^ In the context of PCET, large KIEs reflect the fact that the two-times greater mass of the deuteron *versus* proton causes more localized vibrational wavefunctions, decreased wavefunction overlaps, and thereby reduced tunneling probabilities.^22^

On a classical electrostatics level, constant pH molecular dynamics simulations in the present work indicated that changes in pK_a_s were coupled to the redox processes so as to compensate for the change in charge.^142^ The net charge of the protein was found to change by only ~50% of the expected −6*e* in the physiologically relevant 5 to 7 pH range (Table S30; Figure S11) upon reduction of the six central hemes in the filament. This charge-compensation effect was determined by computing the pK_a_s for all 82 titratable residues within the central subunit of the filament before and after a 6-electron reduction of the hemes in that subunit (Table S29) and applying the Henderson-Hasselbalch equation at different pHs. Charge compensation was also used to rationalize^39^ the increased electrical conductivity observed for pili^36^/OmcS^55^ filaments as the pH was lowered

### 3.3. Can temperature-dependent energetics explain *anti*-Arrhenius electrical conductivity in OmcS?

The ⟨*H*_*mn*_⟩ and λ_*mn*_ changed by only |0.001| and |0.1| eV, respectively, from 300/310 to 270 K in the present and prior^83^ studies. Δ*G*_*mn*_ changed by |0.05| eV from 300 to 270 K in the present work, which was well within the errors of the computations. ΔΔ*G*_*mn*_s 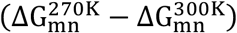 were relatively small because changes in the redox potential differences between hemes were small. All the *E*° in the present study shifted in the same (negative) direction by a nearly uniform amount: −0.07±0.03 (average ± standard deviation). By contrast, ΔΔ*G*_*mn*_ in the prior^83^ study ranged from −0.3 to +0.1 eV because the underlying *E*°s shifted by −0.189 to +0.113 V upon cooling from 310 to 270 K. Thus, a more dramatic resculpting of the free energy landscape upon cooling was found in the prior work.

Irrespective of which temperature dependence for Δ*G*_*mn*_ is more reasonable, the first question is: Can either result account for the *anti*-Arrhenius temperature dependence? “No” is the answer from the analysis in Figure 8. Only a 4-, not 77-times increase in hopping conductivity was simulated using an analytical version of the diffusion model implemented with variable-timestep KMC in the prior work.^83^ The flux kinetic model even found a decrease. What can account for this discrepancy?

The answer seems to lie in methodological details. Dahl *et al*.^83^ implemented a KMC scheme to compute the diffusion constant for a periodic one-dimensional hopping chain. The present work used the analytical diffusion formula derived by Derrida and implemented by Jansson^143, 144^ (see §S1.6.1 in the SI) for the same physical system. In principle these two approaches should agree in the limit of converged statistics for the KMC approach.

The diffusion constant was computed in the prior work from the slope of a least-squares fit to a mean-squared-displacement *versus* time plot for the diffusing particle. The squared displacement at each MC step was averaged over the number of MC steps; a procedure I, as a co-author, discovered after publication. Problematically, the MC step count is not a physical observable and corresponded to a variable time increment in the KMC simulations.

Proposed strategies for computing mean-squared-displacements as a function of time have been described for variable time-step KMC simulations.^145, 146^ These methods involve either (1) running KMC simulations for a few hundred-million steps (only 10^5^ were used by Dahl *et al*.^83^) and then filtering the data to create a trajectory with nearly equidistant time points, or (2) constructing histograms of time intervals and squared displacements computed at each step with *N* previous steps. These strategies suggest that more care is needed in averaging squared displacements from variable time-step KMC simulations than dividing by the number of unequally sized (in time) MC steps. Note that the publication from which the KMC methodology was adapted did not average over the number of MC steps.^147^

Notwithstanding the implementation issue in the prior work,^83^ an increase in conductivity upon cooling was still found (Figure 8) using the same energetic parameters with the diffusion model. Unfortunately, the simulated *anti*-Arrhenius behavior that remains resulted entirely from an abnormally negative *E*° for heme **#5** at 310 K in the prior work. If the *E*° for that heme is shifted by +0.3 V, as suggested by the multiple and more rigorous methods reported here (§2.2), and all other energetic parameters from the prior work are retained, a 1- to 2-fold *decrease* in the simulated conductivity was found upon cooling from 310 to 270 K (Figure 8, compare the 310 K entry marked with an asterisk to the result at 270 K for Dahl *et al*.). A decrease in conductivity was found even if *E*° is shifted only by half as much (Table S27, S28).

Note that the redox potentials for *all other hemes* at 310 and 270 K in the prior work deviated from the ones reported here at 300 and 270 K by half as much as for heme **#5**; these deviations were within or slightly above the standard error of the computations (< 0.160 V). Also note that the 0.3 V more negative *E*° in the prior work cannot be ascribed to the 10 K higher temperature used there. Doing so would suppose a 0.03 V/K shift, which is 10-times larger than experimentally determined temperature coefficients for a variety of cytochromes.^86–92^ ^88, 91, 93^ Some special explanation peculiar to the simulations reported by Dahl *et al*.^83^ seems to be involved.

The far-too-negative *E*° for heme **#5** in the prior work dramatically changed the rates to and from this heme: The ratios 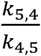 and 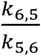 were 10^4^ smaller and 10^5^ larger, respectively, than those found in the present work or reported by Jiang *et al*.^82^ (Table S26). The resulting diffusion constant was 551-times smaller than it would have been with *E*° shifted by +0.3 V (2.7 × 10^−11^ *versus* 1.5 × 10^−8^ cm^2^/s). This underestimation of the diffusion constant at 310 K may have caused the diffusion constant at 270 K (1.1 × 10^−10^ cm^2^/s) to appear larger.

Overall, the prior reproduction of the *anti*-Arrhenius temperature dependence seems to have been the result of (1) an unrecommended way to compute the diffusion constant from variable time-step KMC simulations, and (2) a large underestimation of the charge mobility at 310 K because of an abnormally negative *E*° that is not supported by the work of Jiang *et al*.^82^, the more rigorous calculations reported here, or experimental expectations.^94^

Answering the question that titles this section: The temperature dependence of electron transfer energetics, at least within the framework of ‘vanilla’ Marcus theory,^148^ do not account for the *anti*-Arrhenius conductivity in OmcS. This failure may reflect the mismatch between the real and simulated physical conditions for the filaments, or a breakdown of approximations behind the Marcus expression in Eq. 2. Some of the assumptions that may breakdown with cooling include the high-temperature classical limit for thermal population of vibrations and a relaxation time of the environment that exceeds the electron transfer rate.^148, 149^ These issues are discussed more in the next section.

### 3.4. What additional factors may contribute to *anti*-Arrhenius electrical conductivity?

An *anti*-Arrhenius temperature dependence, as found for electrical conductivity in OmcS between 270 and 300 K (Figure 1, *bottom*), reflects a negative (apparent) activation enthalpy (Δ*H*^‡^). Marcus demonstrated (Eq. 3) that a negative Δ*H*^‡^ can result if the enthalpic change for a reaction (Δ*H*°) is sufficiently negative, and/or if the entropic change for the reaction (Δ*S*°) is sufficiently large (positive or negative).^150, 151^

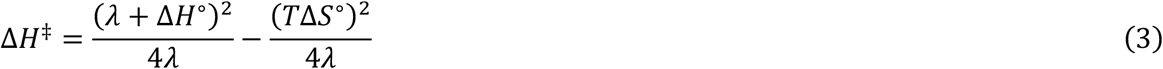

Eq. 3 is a consequence of the fact that parabolic enthalpy curves for the reactant and product states are displaced from the corresponding free energy curves when Δ*S*° ≠ 0. The equation was derived assuming equal-curvature parabolas, weak donor-acceptor electronic couplings, and temperature independent reorganization energies—all of which reasonably apply to OmcS based on the presented computations (Figures S2, S4; Tables S23, S25).

Marcus used Eq. 3 to explain^150, 151^ the *anti*-Arrhenius temperature dependence of electron transfer between Fe(H_2_O)_6_^2+^ and Fe^3+^ or Ru^3+^ polypyridine complexes.^152, 153^ The measured negative Δ*H*° and Δ*S*° indicated that the product quantum states were shifted to lower energy and were more widely spaced than those of the reactants (see Figure 1 in Ref ^151^). If these shifts were large enough, the Boltzmann-averaged enthalpy of the activated state could fall below the Boltzmann-averaged enthalpy of the reactant state. In this scenario, the lower-energy reactant quantum states would react more readily than the higher energy ones, giving an *anti*-Arrhenius temperature dependence.

Large (positive or negative) entropic changes upon electron transfer may originate in several ways. A change in the vibrational density of states, for example, was invoked to explain the *anti*-Arrhenius behavior of electron transfer in bacterial photosynthetic reaction cneters.^154, 155^ An increase in rate upon cooling for these activationless reactions was explained in terms of a transfer of population from higher to lower vibrational energy levels in the reactant and product states. The lower energy vibrations had larger Frank-Condon overlap factors, and Marcus theory rates are directly proportional to the Frank-Condon weighted density of states. Changes in frequency for hydrational, porphyrin skeletal, or protein vibrations coupled to the electron transfer were also needed to account for the experimental data. Similarly, high-frequency intra-molecular vibrations that need to be treated quantum mechanically were found to couple to electron transfer in an organic semi-conductor and to result in an *anti*-Arrhenius temperature dependence.^156^

Changes in entropy may also result from differences in the motions of solvent molecules surrounding the reactants and products. Dipolar orientations and translations are two important collective modes of a polar medium that respectively make enthalpic and entropic contributions to the donor-acceptor energy gap.^157^ Entropic dipolar translations manifest as density fluctuations. The analysis in §2.3.1 indicated that an increase in the density of hydration layers within 10 Å of **#5** and **#6** drove the *E*°s of these hemes in the negative direction.

Matyushov and co-workers^157, 158^ showed that dipolar orientations and translations of the solvent impart hyperbolic temperature dependencies, with opposite signs, to the reaction and reorganization free energies of electron transfer. These dependences become manifest when the solvent response is not quasi-macroscopic,^159^ which may describe the case of a partially dried protein sample as in the temperature-dependent experiments on OmcS. The result is a bell-shaped curve in the Arrhenius (e.g., ln(*k*_*nm*_) vs. 1/T) coordinates, analogous in some respects to the shape of the plot in Figure 1. This phenomenology was used to explain the bell-shaped temperature dependence of charge recombination in a porphyrin-bridge-fullerene complex.^157^

Another way the solvent can produce a bell-shaped Arrhenius plot is if the solvent differentially wets one of the two charge states of the redox protein. In the case of ferredoxin, the temperature dependence of the reorganization energy produced a bell-shaped dependence of the activation energy on inverse temperature.^160^

Comparisons to metal-molecule-metal or molecular junctions are perhaps more relevant for interpreting electrical measurements of OmcS than these solution-phase examples. Marcus showed that inclusion of entropic effects results in a temperature-dependent shift in the position of the molecular energy level relative to the electronic states of the electrodes.^132^ The result in an *anti*-Arrhenius temperature dependence in the off-resonant and resonant regimes for electron transport through the device.

In the same type of molecular junction, Nitzan and co-workers^161^ found that the average current for hopping conduction under low applied bias (0.1 V) can switch from a direct to inverse proportionality on the solvent relaxation time (*γ*) as *γ* increases. Given that *γ*, or the viscosity of the medium increases with decreasing temperature, this finding predicts a switch from an *anti*-Arrhenius to an Arrhenius dependence of the current as the temperature is lowered.^162^

Nitzan and co-workers dubbed this phenomenon a “Kramers-like turnover” in recognition of Kramers’ rate theory,^163^ which describes solvent-controlled dynamics when the relaxation time of the medium is comparable to the passage time through the transition state region along the reaction coordinate. Kramers’ turnovers have been used to explain the *anti*-Arrhenius-to-Arrhenius temperature dependence of *trans*-to-*cis* photo-isomerization of stilbene, the positronium quenching rate constant for formation of a positronium-acceptor complex,^162^ and may contribute to the *anti*-Arrhenius-to-Arrhenius crossover observed in Figure 1 *bottom* for OmcS.

Still in the context of electrical measurements, Matyushov and co-workers pointed out that thermally induced oscillations of the protein-electrode distance introduce temperature-dependent contributions to the activation enthalpy and entropy.^164^ At high enough temperatures, the entropic term in an equation analogous to Eq. 3 (Eq. 65 in Ref. ^164^), which originates from the mean-squared-displacement of the protein atoms, is anticipated to cause a negative activation energy, or *anti*-Arrhenius behavior. Further, Murgida and co-workers^165^ showed experimentally that the temperature dependence of medium relaxation modes influenced electron transfer rates under molecular crowding conditions, a conclusion probably relevant to electrochemistry on films of OmcS.

Lowering of the temperature may cause, as already noted, the electron transfer rate to be sensitive to solvent friction by making nuclear modes coupled to the reaction coordinate sluggish. If the temperature is further lowered, some of these modes can become dynamically arrested. Below a crossover temperature, the dynamics of the protein are non-ergodic, and the reorganization energy decreases (instead of increases as expected) upon further cooling.^166^ This effect was able to explain the *anti*-Arrhenius temperature dependence of self-exchange electron transfer in mixed-valence complexes as the temperatures neared the point of solvent crystallization.^166^

Finally, apart from entropic or non-ergodicity effects, or solvent-friction controlled reaction dynamics, a highly speculative proposal is that the *anti*-Arrhenius-to-Arrhenius temperature dependence of conductivity in OmcS reflects a crossover from thermally activated to tunneling transport upon cooling.^167^ Thermally activated over-barrier crossings dominate at high temperature, whereas through-barrier tunneling dominates at low temperature. The crossover temperature (*T*_*c*_) from over- to through-barrier mechanisms depends on the specific reaction; for hydrogen atom transfers, *T*_*c*_ is commonly near room temperature.^168^ The overall reaction rate in the intermediate range between the high and low temperature extremes often exceeds the (extrapolated) thermal rate and the low-temperature tunneling limit combined; a phenomenon dubbed temperature-assisted tunneling or vibrationally activated tunneling.^168, 169^ A consequence of this effect is typically a convex, instead of a linear line in an Arrhenius ln(*k*_*nm*_) vs. 1/T plot, which is called “sub-Arrhenius” behavior.^170^ However, if the increase in through-barrier tunneling upon cooling can over-compensate for the decrease in over-barrier transitions, an *anti*-Arrhenius temperature dependence may be observed down to *T*_*c*_. Below *T*_*c*_, the conductivity may appear to decrease because one of the transport channels (over-barrier transitions) has now been completely shut down.

An analogous explanation was advanced for the *anti*-Arrhenius-to-Arrhenius temperature dependence of heterogeneous ammonia synthesis in the presence of an electric field.^171^ Associative and dissociative mechanisms were proposed to be operative below 373 K and above 573 K, respectively, and both exhibit Arrhenius-type dependencies on temperature. Between 373 and 573 K, both mechanisms were proposed to be operative to some extent to explain the *anti*-Arrhenius temperature dependence.

Taken together, there are a plethora of ways in which changes in entropy and medium relaxation times relative to the rate of electron transfer can contribute to the observed anti-Arrhenius—and more generally, bell-shaped—dependence of electrical conductivity on temperature in OmcS. Virtually all the examples reviewed in this section involve simpler systems (e.g., a single-step electron transfer or a single energy level in theoretical work), so a quantitative explanation for the behavior in OmcS cannot be expected. The hope, however, is that this recitation of interpretations for *anti*-Arrhenius phenomena will inspire new experiments in *silico* and *reapse*.

## 4. Conclusion

To summarize, QM/MM@MD-computed *E*°s in OmcS were −0.063 – −0.271 V vs. SHE at 300 K, consistent with experiments on bis-histidine-ligated *c*-type hemes.^94^ Cooling by 30 K induced negative and nearly uniform (−0.07 ± 0.03 V) shifts that were consistent with the magnitude of temperature coefficients measured for a variety of cytochromes.^86–92^ ^88, 91, 93^ Much of the results with QM/MM@MD were corroborated by C(E,pH)MD: *E*° range = −0.093 – −0.306 V vs. SHE and mean unsigned error = ~0.09 V relative to QM/MM@MD.

Changes in *E*° were well-matched by negative shifts in electrostatic energy for heme oxidation. A dominant contributor to the largest of these shifts, which were experienced by hemes **#5** and **#6**, originated from the aqueous solvent. Hydrating layers within 10 Å of hemes **#5** and **#6** became denser at lower temperature (above the normal freezing point). The static dielectric constant of water is known experimentally to increase upon cooling,^117^ and in the present study, the *E*° of the hemes was shown to be inversely correlated with the static dielectric constant of the medium. Thus, the data suggests that an increase in the density of hydration layers around the filament upon cooling increased the static dielectric constant, and in turn, shifted the *E*°s of the hemes to more negative values.

Changes in descriptors of the H-bonding network were also observed and virtually reproduced those reported earlier.^83^ However, the sign and size of the cooling-induced shifts found for *E*° were very different from that study, suggesting H-bond restructuring was not the controlling factor for this effect.

The uniformity of the cooling-induced shifts in *E*° found here with QM/MM@MD resulted in mean-unsigned changes in Δ*G*_*mn*_ of ~0.04 eV, which were not statistically meaningful. Cooling induced mean-unsigned shifts in λ_*mn*_ (0.05 eV) and ⟨*H*_*mn*_⟩ (0.001 eV) were similarly small. The 30 K drop in temperature was found to only be a weak perturbation on these energetic terms in the present study.

As a result, all heme-to-heme Marcus theory electron transfer rates differed by a factor of ≤ 4 at 270 *vs*. 300 K. The charge diffusion constant and protein-limited flux (averaged over forward and reverse directions) was only ~2-times smaller at the lower temperature. Thus, the data suggests that the cooling-induced changes in energetic terms within standard Marcus theory (Eq. 2) predicts a slight Arrhenius, not a dramatic *anti*-Arrhenius temperature dependence for OmcS within the 270-300 K range. Entropic effects^150, 151^ from changes in vibrations^154, 155^ (including the oscillations of the protein-electrode distance),^164^ solvent density^157, 159^ or redox-linked active site wetting,^160^ and renormalization of molecular energy levels in the electrical device;^132^ a Kramers’-like turnover from the under to over-damped regime in solvent fraction-controlled reaction dynamics;^161^ or non-ergodicity effects^166^ may be involved in the *anti*-Arrhenius behavior.

A similarly small cooling-induced effect on charge mobility was also observed with the much larger resculpting of the free energy landscape reported by Dahl *et al*.^83^ This prior work reported shifts in *E*° from 310 to 270 K of mixed sign that were as large as 40% (0.189 V) of the experimental range for bis-histidine ligated *c*-type heme.^94^ The shifts were also, at most, twice as large as the shifts reported in the present study for a smaller temperature range. The resulting changes to Δ*G*_*mn*_ were −0.3 to +0.2 eV, or as a mean unsigned average, > 3-times the effect found in the present work.

Using these Δ*G*_*mn*_s, combined with very similar λ_*mn*_s and ⟨*H*_*mn*_⟩s in both studies, the diffusion constant increased by a factor of 4, whereas the protein-limited flux decreased slightly. This seemed reasonable since most of the Marcus rates reported by Dahl *et al*. decreased on average by one order of magnitude from 310 to 270 K; the largest exception being the forward and reverse rates between hemes **#3** and **#4**, which increased by two orders of magnitude.

The simulated *anti*-Arrhenius charge mobility using the rates computed by Dahl *et al*. was much smaller than the 77-times increase reported before. A variable time-step KMC implementation was used in that work to treat the same one-dimensional diffusion problem solved here with Derrida’s analytical formula. The discrepancy likely resulted from an averaging of squared displacements from the KMC simulations by the number of MC steps, which represented variable and unequal time increments. Different averaging procedures have been recommended for this purpose.^145, 146^

Regardless of this technical issue, Arrhenius, instead of *anti*-Arrhenius behavior is predicted with the same energetic parameters published by Dahl *et al*.^83^ if only the *E*° of heme **#5** is shifted by either +0.15 or +0.3 V. These shift seemed warranted because the reported *E*° (−0.521 V vs. SHE) fell ~0.15 V outside the expected experimental range,^94^ and disagreed by ≥0.3 eV from the value obtained with using C(E,pH)MD and more rigorous QM/MM@MD techniques in the present work—a deviation 4-times larger than for any other heme at the same temperature. The *E*°s reported here are also in much better agreement with spectroelectrochemical experiments (manuscript under preparation). Thus, the data suggests that the present and prior theoretical studies *do not* capture the *anti*-Arrhenius conductivity observed experimentally.

The failure in modeling the temperature dependence of electrical conductivity in OmcS is instructive and motivates two hypotheses: Either the hopping mechanism is inappropriate to describe charge transport under solid-state measurement conditions, or the inadequacy of modeling those experimental conditions make the hopping mechanism appear inappropriate. These hypotheses have been discussed before,^84, 134^ but the second of them is not always appreciated, including by the present author at the outset of this research.

Simulations on a fully hydrated and freely floating protein oligomer in electrolytic solution are likely of little relevance for modeling charge transport through an electrode-adsorbed, (partially or fully) dehydrated, and (possibly) aggregated protein. Simply by neglecting some portion of the reorganization energy of the aqueous solvent and leaving all other energetic parameters the same, a few orders of magnitude improvement in the predicted hopping current was found relative to what the experimental value would be (hypothetically) at the same voltage.

In closing, better experimental and theoretical characterizations of microbial filaments under experimental conditions are essential before a verdict can be reached on the operative mechanism of charge transport and its response to changes in physical state. Key questions relate to the structural and electronic consequences of, for example, surface adsorption, hydration state, and protein-protein contacts.

## Notes

I have no conflicts of interest to declare.

## Acknowledgements

This research was supported by the National Institute of General Medical Sciences of the National Institutes of Health under Award Number 1F32GM142247-01A1. The research was started while I was funded by Nikhil S. Malvankar via NSF CAREER award no. 1749662 and the Career Award at the Scientific Interfaces from Burroughs Welcome Fund. All calculations were performed using resources at the Yale High Performance Computing Center. Implementations of the diffusion and flux kinetic models were kindly provided, respectively, by Fredrick Jansson and Xiuyun Jiang. I wish to deeply thank Yangqi Gu, Clorice Reinhardt, Vinicius Cruzeiro, Peng Zhang, Andrea Amadei, Laura Zanettipolzi, Michael Buehl, Jonathan Colburn, Tom Langford, and José Gascón for useful conversations related to this research.

## References

1. Torres, C. I.; Marcus, A. K.; Lee, H.-S.; Parameswaran, P.; Krajmalnik-Brown, R.; Rittmann, B. E., A kinetic perspective on extracellular electron transfer by anode-respiring bacteria. FEMS microbiology reviews 2010, 34 (1), 3–17.

2. Matyushov, D. V., Protein electron transfer: is biology (thermo) dynamic? Journal of Physics: Condensed Matter 2015, 27 (47), 473001.

3. Nealson, K. H.; Conrad, P. G., Life: past, present and future. Philosophical Transactions of the Royal Society of London. Series B: Biological Sciences 1999, 354 (1392), 1923–1939.

4. Vargas, M.; Kashefi, K.; Blunt-Harris, E. L.; Lovley, D. R., Microbiological evidence for Fe (III) reduction on early Earth. Nature 1998, 395 (6697), 65–67.

5. Ilbert, M.; Bonnefoy, V., Insight into the evolution of the iron oxidation pathways. Biochimica et Biophysica Acta (BBA)-Bioenergetics 2013, 1827 (2), 161–175.

6. Zhao, J.; Li, F.; Cao, Y.; Zhang, X.; Chen, T.; Song, H.; Wang, Z., Microbial extracellular electron transfer and strategies for engineering electroactive microorganisms. Biotechnology Advances 2021, 53, 107682.

7. Lovley, D. R.; Holmes, D. E., Electromicrobiology: the ecophysiology of phylogenetically diverse electroactive microorganisms. Nature Reviews Microbiology 2022, 20 (1), 5–19.

8. Nealson, K. H.; Saffarini, D., Iron and manganese in anaerobic respiration: environmental significance, physiology, and regulation. Annual review of microbiology 1994, 48, 311–344.

9. Jelen, B. I.; Giovannelli, D.; Falkowski, P. G., The role of microbial electron transfer in the coevolution of the biosphere and geosphere. Annual review of microbiology 2016, 70, 45–62.

10. Jiang, Y.; Shi, M.; Shi, L., Molecular underpinnings for microbial extracellular electron transfer during biogeochemical cycling of earth elements. Science China Life Sciences 2019, 62 (10), 1275–1286.

11. Zhang, X.; Yuan, Z.; Hu, S., Anaerobic oxidation of methane mediated by microbial extracellular respiration. Environmental Microbiology Reports 2021, 13 (6), 790–804.

12. Reyes, C.; Meister, P., The Role of Microorganisms in Iron Reduction in Marine Sediments. Systems Biogeochemistry of Major Marine Biomes 2022, 41–60.

13. Lovley, D. R., Syntrophy goes electric: direct interspecies electron transfer. Annual review of microbiology 2017, 71, 643–664.

14. Wang, W.; Du, Y.; Yang, S.; Du, X.; Li, M.; Lin, B.; Zhou, J.; Lin, L.; Song, Y.; Li, J., Bacterial extracellular electron transfer occurs in mammalian gut. Analytical chemistry 2019, 91 (19), 12138–12141.

15. Schwab, L.; Rago, L.; Koch, C.; Harnisch, F., Identification of Clostridium cochlearium as an electroactive microorganism from the mouse gut microbiome. Bioelectrochemistry 2019, 130, 107334.

16. Khan, M. T.; Duncan, S. H.; Stams, A. J.; Van Dijl, J. M.; Flint, H. J.; Harmsen, H. J., The gut anaerobe Faecalibacterium prausnitzii uses an extracellular electron shuttle to grow at oxic– anoxic interphases. The ISME journal 2012, 6 (8), 1578–1585.

17. Light, S. H.; Su, L.; Rivera-Lugo, R.; Cornejo, J. A.; Louie, A.; Iavarone, A. T.; Ajo-Franklin, C. M.; Portnoy, D. A., A flavin-based extracellular electron transfer mechanism in diverse Gram-positive bacteria. Nature 2018, 562 (7725), 140–144.

18. Bostick, C. D.; Mukhopadhyay, S.; Pecht, I.; Sheves, M.; Cahen, D.; Lederman, D., Protein bioelectronics: A review of what we do and do not know. Reports on Progress in Physics 2018, 81 (2), 026601.

19. Wellman, S. M.; Eles, J. R.; Ludwig, K. A.; Seymour, J. P.; Michelson, N. J.; McFadden, W. E.; Vazquez, A. L.; Kozai, T. D., A materials roadmap to functional neural interface design. Advanced functional materials 2018, 28 (12), 1701269.

20. Ha, T. Q.; Planje, I. J.; White, J. R.; Aragonès, A. C.; Díez-Pérez, I., Charge transport at the protein–electrode interface in the emerging field of BioMolecular Electronics. Current Opinion in Electrochemistry 2021, 28, 100734.

21. Zhang, Y.; Hsu, L. H.-H.; Jiang, X., Living electronics. Nano Research 2020, 13 (5), 1205–1213.

22. Sanjuan-Alberte, P.; Alexander, M. R.; Hague, R. J.; Rawson, F. J., Electrochemically stimulating developments in bioelectronic medicine. Bioelectronic Medicine 2018, 4 (1), 1–7.

23. Sanjuan-Alberte, P.; Rawson, F. J., Engineering the spark into bioelectronic medicine. Future Science: 2019; Vol. 10, pp 139–142.

24. Cohen-Karni, T.; Langer, R.; Kohane, D. S., The smartest materials: the future of nanoelectronics in medicine. ACS nano 2012, 6 (8), 6541–6545.

25. Alfonta, L., Bioelectrochemistry and the Singularity Point “I Robot”? Israel Journal of Chemistry 2021, 61 (1-2), 60–67.

26. Bird, L. J.; Kundu, B. B.; Tschirhart, T.; Corts, A. D.; Su, L.; Gralnick, J. A.; Ajo-Franklin, C. M.; Glaven, S. M., Engineering Wired Life: Synthetic Biology for Electroactive Bacteria. ACS Synthetic Biology 2021, 10 (11), 2808–2823.

27. Lovley, D. R., e-Biologics: fabrication of sustainable electronics with “green” biological materials. MBio 2017, 8 (3), e00695–17.

28. Lovley, D. R.; Yao, J., Intrinsically conductive microbial nanowires for ‘green’electronics with novel functions. Trends in Biotechnology 2021, 39 (9), 940–952.

29. Harnisch, F.; Holtmann, D., Bioelectrosynthesis. Springer: 2019; Vol. 490.

30. Zou, L.; Zhu, F.; Long, Z.-e.; Huang, Y., Bacterial extracellular electron transfer: a powerful route to the green biosynthesis of inorganic nanomaterials for multifunctional applications. Journal of Nanobiotechnology 2021, 19 (1), 1–27.

31. Sun, Y. L.; Tang, H. Y.; Ribbe, A.; Duzhko, V.; Woodard, T. L.; Ward, J. E.; Bai, Y.; Nevin, K. P.; Nonnenmann, S. S.; Russell, T., Conductive composite materials fabricated from microbially produced protein nanowires. Small 2018, 14 (44), 1802624.

32. Gu, T.; Wang, D.; Lekbach, Y.; Xu, D., Extracellular electron transfer in microbial biocorrosion. Current Opinion in Electrochemistry 2021, 29, 100763.

33. Kumar, A.; Hsu, L. H.-H.; Kavanagh, P.; Barrière, F.; Lens, P. N.; Lapinsonnière, L.; Lienhard V J. H.; Schroeder, U.; Jiang, X.; Leech, D., The ins and outs of microorganism–electrode electron transfer reactions. Nature Reviews Chemistry 2017, 1 (3), 1–13.

34. Baek, G.; Kim, J.; Kim, J.; Lee, C., Role and potential of direct interspecies electron transfer in anaerobic digestion. Energies 2018, 11 (1), 107.

35. Reguera, G.; McCarthy, K. D.; Mehta, T.; Nicoll, J. S.; Tuominen, M. T.; Lovley, D. R., Extracellular electron transfer via microbial nanowires. Nature 2005, 435 (7045), 1098–1101.

36. Madeline, V.; Ching, L.; Byoung-Chan, K.; Kengo, I.; Tünde, M., Tunable metallic-like conductivity in microbial nanowire networks. Nature Nanotechnology 2011, 6 (9).

37. Strycharz-Glaven, S. M.; Snider, R. M.; Guiseppi-Elie, A.; Tender, L. M., On the electrical conductivity of microbial nanowires and biofilms. Energy & Environmental Science 2011, 4 (11), 4366–4379.

38. Malvankar, N. S.; Tuominen, M. T.; Lovley, D. R., Comment on “on electrical conductivity of microbial nanowires and biofilms” by SM Strycharz-Glaven, RM Snider, A. Guiseppi-Elie and LM Tender, energy environ. sci., 2011, 4, 4366. Energy & Environmental Science 2012, 5 (3), 6247–6249.

39. Strycharz-Glaven, S. M.; Tender, L. M., Reply to the ‘Comment on “On electrical conductivity of microbial nanowires and biofilms”‘by NS Malvankar, MT Tuominen and DR Lovley, Energy Environ. Sci., 2012, 5. Energy & Environmental Science 2012, 5 (3), 6250–6255.

40. Yates, M. D.; Strycharz-Glaven, S. M.; Golden, J. P.; Roy, J.; Tsoi, S.; Erickson, J. S.; El-Naggar, M. Y.; Barton, S. C.; Tender, L. M., Measuring conductivity of living Geobacter sulfurreducens biofilms. Nature nanotechnology 2016, 11 (11), 910–913.

41. Malvankar, N. S.; Rotello, V. M.; Tuominen, M. T.; Lovley, D. R., Reply to’Measuring conductivity of living Geobacter sulfurreducens biofilms’. Nature Nanotechnology 2016, 11 (11), 913–914.

42. Nealson, K. H.; Myers, C. R., Microbial reduction of manganese and iron: new approaches to carbon cycling. Applied and environmental microbiology 1992, 58 (2), 439–443.

43. Reguera, G.; Kashefi, K., The electrifying physiology of Geobacter bacteria, 30 years on. Advances in Microbial Physiology 2019, 74, 1–96.

44. Boesen, T.; Nielsen, L. P.; Schramm, A., Pili for nanowires. Nature Microbiology 2021, 6 (11), 1347–1348.

45. Wang, F.; Gu, Y.; O’Brien, J. P.; Yi, S. M.; Yalcin, S. E.; Srikanth, V.; Shen, C.; Vu, D.; Ing, N. L.; Hochbaum, A. I.; Egelman, E. H.; Malvankar, N. S., Structure of Microbial Nanowires Reveals Stacked Hemes that Transport Electrons over Micrometers. Cell 2019, 177 (2), 361-369.e10.

46. Lovley, D. R.; Walker, D. J., Geobacter protein nanowires. Frontiers in microbiology 2019, 10, 2078.

47. Yalcin, S. E.; Malvankar, N. S., The blind men and the filament: understanding structures and functions of microbial nanowires. Current opinion in chemical biology 2020, 59, 193–201.

48. Lovley, D. R.; Holmes, D. E., Protein nanowires: the electrification of the microbial world and maybe our own. Journal of bacteriology 2020, 202 (20), e00331–20.

49. Clark, M. M.; Reguera, G., Biology and biotechnology of microbial pilus nanowires. Journal of Industrial Microbiology & Biotechnology: Official Journal of the Society for Industrial Microbiology and Biotechnology 2020, 47 (9-10), 897–907.

50. Gu, Y.; Srikanth, V.; Salazar-Morales, A. I.; Jain, R.; O’Brien, J. P.; Yi, S. M.; Soni, R. K.; Samatey, F. A.; Yalcin, S. E.; Malvankar, N. S., Structure of Geobacter pili reveals secretory rather than nanowire behaviour. Nature 2021, 597 (7876), 430–434.

51. Lovley, D. R., On the Existence of Pilin-Based Microbial Nanowires. Frontiers in Microbiology, 2069.

52. Ye, Y.; Liu, X.; Nealson, K. H.; Rensing, C.; Qin, S.; Zhou, S., Dissecting the Structural and Conductive Functions of Nanowires in Geobacter sulfurreducens Electroactive Biofilms. Mbio 2022, 13 (1), e03822–21.

53. Lovley, D. R., Untangling Geobacter sulfurreducens Nanowires. mBio 2022, e00850-22.

54. Liu, X.; Nealson, K. H.; Zhou, S.; Rensing, C., Reply to Lovley,”Untangling Geobacter sulfurreducens Nanowires”. mBio 2022, e01041–22.

55. Yalcin, S. E.; O’Brien, J. P.; Gu, Y.; Reiss, K.; Yi, S. M.; Jain, R.; Srikanth, V.; Dahl, P. J.; Huynh, W.; Vu, D., Electric field stimulates production of highly conductive microbial OmcZ nanowires. Nature chemical biology 2020, 16 (10), 1136–1142.

56. Hospenthal, M. K.; Costa, T. R.; Waksman, G., A comprehensive guide to pilus biogenesis in Gram-negative bacteria. Nature Reviews Microbiology 2017, 15 (6), 365.

57. Lovley, D. R., Electrically conductive pili: biological function and potential applications in electronics. Current Opinion in Electrochemistry 2017, 4 (1), 190–198.

58. Creasey, R. C.; Mostert, A. B.; Nguyen, T. A.; Virdis, B.; Freguia, S.; Laycock, B., Microbial nanowires–electron transport and the role of synthetic analogues. Acta Biomaterialia 2018, 69, 1–30.

59. Nathanael, J. G.; Gamon, L. F.; Cordes, M.; Rablen, P. R.; Bally, T.; Fromm, K. M.; Giese, B.; Wille, U., Amide Neighbouring-Group Effects in Peptides: Phenylalanine as Relay Amino Acid in Long-Distance Electron Transfer. ChemBioChem 2018, 19 (9), 922–926.

60. Guterman, T.; Gazit, E., Toward peptide-based bioelectronics: reductionist design of conductive pili mimetics. Bioelectronics in medicine 2018, 1 (2), 131–137.

61. Ing, N. L.; Spencer, R. K.; Luong, S. H.; Nguyen, H. D.; Hochbaum, A. I., Electronic conductivity in biomimetic α-helical peptide nanofibers and gels. ACS nano 2018, 12 (3), 2652–2661.

62. Shipps, C.; Kelly, H. R.; Dahl, P. J.; Yi, S. M.; Vu, D.; Boyer, D.; Glynn, C.; Sawaya, M. R.; Eisenberg, D.; Batista, V. S., Intrinsic electronic conductivity of individual atomically resolved amyloid crystals reveals micrometer-long hole hopping via tyrosines. Proceedings of the National Academy of Sciences 2021, 118 (2), e2014139118.

63. Wang, Y.; Pu, J.; An, B.; Lu, T. K.; Zhong, C., Emerging paradigms for synthetic design of functional amyloids. Journal of molecular biology 2018, 430 (20), 3720–3734.

64. Kalyoncu, E.; Ahan, R. E.; Olmez, T. T.; Seker, U. O. S., Genetically encoded conductive protein nanofibers secreted by engineered cells. RSC advances 2017, 7 (52), 32543–32551.

65. Shapiro, D. M.; Mandava, G.; Yalcin, S. E.; Arranz-Gibert, P.; Dahl, P. J.; Shipps, C.; Gu, Y.; Srikanth, V.; Salazar-Morales, A. I.; O’Brien, J. P., Protein nanowires with tunable functionality and programmable self-assembly using sequence-controlled synthesis. Nature communications 2022, 13 (1), 1–10.

66. Creasey, R. C.; Shingaya, Y.; Nakayama, T., Improved electrical conductance through self-assembly of bioinspired peptides into nanoscale fibers. Materials Chemistry and Physics 2015, 158, 52–59.

67. Creasey, R. C.; Mostert, A. B.; Solemanifar, A.; Nguyen, T. A.; Virdis, B.; Freguia, S.; Laycock, B., Biomimetic peptide nanowires designed for conductivity. ACS Omega 2019, 4 (1), 1748–1756.

68. Ivnitski, D.; Amit, M.; Silberbush, O.; Atsmon-Raz, Y.; Nanda, J.; Cohen-Luria, R.; Miller, Y.; Ashkenasy, G.; Ashkenasy, N., The strong influence of structure polymorphism on the conductivity of peptide fibrils. Angewandte Chemie International Edition 2016, 55 (34), 9988–9992.

69. Lewis, D. K.; Oh, Y.; Mohanam, L. N.; Wierzbicki, M.; Ing, N. L.; Gu, L.; Hochbaum, A.; Wu, R.; Cui, Q.; Sharifzadeh, S., Electronic Structure of de Novo Peptide ACC-Hex from First Principles. The Journal of Physical Chemistry B 2022.

70. Reardon, P. N.; Mueller, K. T., Structure of the type IVa major pilin from the electrically conductive bacterial nanowires of Geobacter sulfurreducens. Journal of Biological Chemistry 2013, 288 (41), 29260–29266.

71. Bonanni, P. S.; Massazza, D.; Busalmen, J. P., Stepping stones in the electron transport from cells to electrodes in Geobacter sulfurreducens biofilms. Physical Chemistry Chemical Physics 2013, 15 (25), 10300–10306.

72. Feliciano, G.; Steidl, R.; Reguera, G., Structural and functional insights into the conductive pili of Geobacter sulfurreducens revealed in molecular dynamics simulations. Physical Chemistry Chemical Physics 2015, 17 (34), 22217–22226.

73. Yan, H.; Chuang, C.; Zhugayevych, A.; Tretiak, S.; Dahlquist, F. W.; Bazan, G. C., Inter-Aromatic Distances in Geobacter Sulfurreducens Pili Relevant to Biofilm Charge Transport. Advanced Materials 2015, 27 (11), 1908–1911.

74. Xiao, K.; Malvankar, N. S.; Shu, C.; Martz, E.; Lovley, D. R.; Sun, X., Low energy atomic models suggesting a pilus structure that could account for electrical conductivity of Geobacter sulfurreducens pili. Scientific reports 2016, 6 (1), 1–9.

75. Shu, C.; Xiao, K.; Sun, X., Structural Basis for the Influence of A1, 5A, and W51W57 Mutations on the Conductivity of the Geobacter sulfurreducens Pili. Crystals 2018, 8 (1), 10.

76. Ru, X.; Zhang, P.; Beratan, D. N., Assessing Possible Mechanisms of Micrometer-Scale Electron Transfer in Heme-Free Geobacter sulfurreducens Pili. The Journal of Physical Chemistry B 2019, 123 (24), 5035–5047.

77. Shu, C.; Zhu, Q.; Xiao, K.; Hou, Y.; Ma, H.; Ma, J.; Sun, X., Direct Extracellular Electron Transfer of the Geobacter sulfurreducens Pili Relevant to Interaromatic Distances. BioMed Research International 2019, 2019.

78. Shu, C.; Xiao, K.; Sun, X., Structural Basis for the High Conductivity of Microbial Pili as Potential Nanowires. Journal of nanoscience and nanotechnology 2020, 20 (1), 64–80.

79. Tan, Y.; Adhikari, R. Y.; Malvankar, N. S.; Ward, J. E.; Woodard, T. L.; Nevin, K. P.; Lovley, D. R., Expressing the Geobacter metallireducens PilA in Geobacter sulfurreducens yields pili with exceptional conductivity. MBio 2017, 8 (1), e02203–16.

80. Mezzina Freitas, L. P.; Feliciano, G. T., Atomic and Electronic Structure of Pilus from Geobacter sulfurreducens through QM/MM Calculations: Evidence for Hole Transfer in Aromatic Residues. The Journal of Physical Chemistry B 2021, 125 (30), 8305–8312.

81. Wang, F.; Mustafa, K.; Suciu, V.; Joshi, K.; Chan, C. H.; Choi, S.; Su, Z.; Si, D.; Hochbaum, A. I.; Egelman, E. H., Cryo-EM structure of an extracellular Geobacter OmcE cytochrome filament reveals tetrahaem packing. Nature Microbiology 2022, 1-10.

82. Jiang, X.; van Wonderen, J. H.; Butt, J. N.; Edwards, M. J.; Clarke, T. A.; Blumberger, J., Which multi-heme protein complex transfers electrons more efficiently? Comparing MtrCAB from Shewanella with OmcS from Geobacter. The Journal of Physical Chemistry Letters 2020, 11 (21), 9421–9425.

83. Dahl, P. J.; Yi, S. M.; Gu, Y.; Acharya, A.; Shipps, C.; Neu, J.; O’Brien, J. P.; Morzan, U. N.; Chaudhuri, S.; Guberman-Pfeffer, M. J., A 300-fold conductivity increase in microbial cytochrome nanowires due to temperature-induced restructuring of hydrogen bonding networks. Science Advances 2022, 8 (19), eabm7193.

84. Eshel, Y.; Peskin, U.; Amdursky, N., Coherence-assisted electron diffusion across the multi-heme protein-based bacterial nanowire. Nanotechnology 2020, 31 (31), 314002.

85. Livernois, W.; Anantram, M. In Quantum Transport in Conductive Bacterial Nanowires, 2021 IEEE 16th Nanotechnology Materials and Devices Conference (NMDC), IEEE: 2021; pp 1–5.

86. Bertrand, P.; Mbarki, O.; Asso, M.; Blanchard, L.; Guerlesquin, F.; Tegoni, M., Control of the redox potential in c-type cytochromes: importance of the entropic contribution. Biochemistry 1995, 34 (35), 11071–11079.

87. Verhagen, M. F.; Wolbert, R. B.; Hagen, W. R., Cytochrome c553 from Desulfovibrio vulgaris (Hildenborough) Electrochemical properties and electron transfer with hydrogenase. European journal of biochemistry 1994, 221 (2), 821–829.

88. Battistuzzi, G.; Bellei, M.; Borsari, M.; Di Rocco, G.; Ranieri, A.; Sola, M., Axial ligation and polypeptide matrix effects on the reduction potential of heme proteins probed on their cyanide adducts. JBIC Journal of Biological Inorganic Chemistry 2005, 10 (6), 643–651.

89. Larroque, C.; Maurel, P.; Douzou, P., Redox potentials in hydro-organic media at normal and subzero temperatures Ferro-ferricyanide and cytochrome c as models. Biochimica et Biophysica Acta (BBA)-Bioenergetics 1978, 501 (1), 20–32.

90. Anderson, C. W.; Halsall, H. B.; Heineman, W. R.; Kreishman, G. P., The temperature dependence of the redox potential of horse heart cytochrome c in sodium chloride solutions. Biochemical and Biophysical Research Communications 1977, 76 (2), 339–344.

91. Battistuzzi, G.; Borsari, M.; Cowan, J. A.; Ranieri, A.; Sola, M., Control of cytochrome c redox potential: axial ligation and protein environment effects. Journal of the American Chemical Society 2002, 124 (19), 5315–5324.

92. Reid, L. S.; Taniguchi, V. T.; Gray, H. B.; Mauk, A. G., Oxidation-reduction equilibrium of cytochrome b5. Journal of the American Chemical Society 1982, 104 (26), 7516–7519.

93. Battistuzzi, G.; Borsari, M.; Ranieri, A.; Sola, M., Redox thermodynamics of the Fe3+/Fe2+ couple in horseradish peroxidase and its cyanide complex. Journal of the American Chemical Society 2002, 124 (1), 26–27.

94. Zheng, Z.; Gunner, M., Analysis of the electrochemistry of hemes with Ems spanning 800 mV. Proteins: Structure, Function, and Bioinformatics 2009, 75 (3), 719–734.

95. Guerard, J. J.; Tentscher, P. R.; Seijo, M.; Arey, J. S., Explicit solvent simulations of the aqueous oxidation potential and reorganization energy for neutral molecules: gas phase, linear solvent response, and non-linear response contributions. Physical Chemistry Chemical Physics 2015, 17 (22), 14811–14826.

96. Ghosh, D.; Roy, A.; Seidel, R.; Winter, B.; Bradforth, S.; Krylov, A. I., First-principle protocol for calculating ionization energies and redox potentials of solvated molecules and ions: theory and application to aqueous phenol and phenolate. The Journal of Physical Chemistry B 2012, 116 (24), 7269–7280.

97. Karnaukh, E. A.; Bravaya, K. B., The redox potential of a heme cofactor in Nitrosomonas europaea cytochrome c peroxidase: A polarizable QM/MM study. Physical Chemistry Chemical Physics 2021, 23 (31), 16506–16515.

98. Crespo, A.; Martí, M. A.; Kalko, S. G.; Morreale, A.; Orozco, M.; Gelpi, J. L.; Luque, F. J.; Estrin, D. A., Theoretical study of the truncated hemoglobin HbN: exploring the molecular basis of the NO detoxification mechanism. Journal of the American Chemical Society 2005, 127 (12), 4433–4444.

99. Henriques, J.; Costa, P. J.; Calhorda, M. J.; Machuqueiro, M., Charge parametrization of the DvH-c3 heme group: validation using constant-(pH,E) molecular dynamics simulations. J Phys Chem B 2013, 117 (1), 70–82.

100. Cruzeiro, V. W. D.; Amaral, M. S.; Roitberg, A. E., Redox potential replica exchange molecular dynamics at constant pH in AMBER: Implementation and validation. J Chem Phys 2018, 149 (7), 072338.

101. Cruzeiro, V. W. D.; Feliciano, G. T.; Roitberg, A. E., Exploring Coupled Redox and pH Processes with a Force-Field-Based Approach: Applications to Five Different Systems. J Am Chem Soc 2020, 142 (8), 3823–3835.

102. Wang, X.; He, X., An Ab Initio QM/MM Study of the Electrostatic Contribution to Catalysis in the Active Site of Ketosteroid Isomerase. Molecules 2018, 23 (10), 2410.

103. Smith, D. M.; Dupuis, M.; Vorpagel, E. R.; Straatsma, T., Characterization of electronic structure and properties of a bis (histidine) heme model complex. Journal of the American Chemical Society 2003, 125 (9), 2711–2717.

104. Rovira, C.; Kunc, K.; Hutter, J.; Ballone, P.; Parrinello, M., Equilibrium geometries and electronic structure of iron− porphyrin complexes: A density functional study. The Journal of Physical Chemistry A 1997, 101 (47), 8914–8925.

105. Kozlowski, P. M.; Spiro, T. G.; Bérces, A.; Zgierski, M. Z., Low-lying spin states of iron (II) porphine. The Journal of Physical Chemistry B 1998, 102 (14), 2603–2608.

106. Smith, D. M.; Rosso, K. M.; Dupuis, M.; Valiev, M.; Straatsma, T., Electronic coupling between heme electron-transfer centers and its decay with distance depends strongly on relative orientation. The Journal of Physical Chemistry B 2006, 110 (31), 15582–15588.

107. Johansson, M. P.; Sundholm, D.; Gerfen, G.; Wikström, M., The spin distribution in low-spin iron porphyrins. Journal of the American Chemical Society 2002, 124 (39), 11771–11780.

108. Johansson, M. P.; Blomberg, M. R.; Sundholm, D.; Wikström, M., Change in electron and spin density upon electron transfer to haem. Biochimica et Biophysica Acta (BBA)-Bioenergetics 2002, 1553 (3), 183–187.

109. McMahon, M. T.; DeDios, A. C.; Godbout, N.; Salzmann, R.; Laws, D. D.; Le, H.; Havlin, R. H.; Oldfield, E., An experimental and quantum chemical investigation of CO binding to heme proteins and model systems: a unified model based on 13C, 17O, and 57Fe nuclear magnetic resonance and 57Fe Mössbauer and infrared spectroscopies. Journal of the American Chemical Society 1998, 120 (19), 4784–4797.

110. Zhang, Y.; Mao, J.; Oldfield, E., 57Fe Mössbauer isomer shifts of heme protein model systems: electronic structure calculations. Journal of the American Chemical Society 2002, 124 (26), 7829–7839.

111. Havlin, R. H.; Godbout, N.; Salzmann, R.; Wojdelski, M.; Arnold, W.; Schulz, C. E.; Oldfield, E., An experimental and density functional theoretical investigation of iron-57 Mössbauer quadrupole splittings in organometallic and heme-model compounds: applications to carbonmonoxy-heme protein structure. Journal of the American Chemical Society 1998, 120 (13), 3144–3151.

112. Cruzeiro, V. W. D.; Amaral, M. S.; Roitberg, A. E., Redox potential replica exchange molecular dynamics at constant pH in AMBER: Implementation and validation. The Journal of chemical physics 2018, 149 (7), 072338.

113. Breuer, M.; Rosso, K. M.; Blumberger, J., Electron flow in multiheme bacterial cytochromes is a balancing act between heme electronic interaction and redox potentials. Proceedings of the National Academy of Sciences 2014, 111 (2), 611–616.

114. Bradshaw, R. T.; Dziedzic, J.; Skylaris, C.-K.; Essex, J. W., The role of electrostatics in enzymes: do biomolecular force fields reflect protein electric fields? Journal of chemical information and modeling 2020, 60 (6), 3131–3144.

115. Matsui, T.; Song, J.-W., A Density Functional Theory-Based Scheme to Compute the Redox Potential of a Transition Metal Complex: Applications to Heme Compound. Molecules 2019, 24 (4), 819.

116. O’Donoghue, D.; Magner, E., The redox thermodynamics of microperoxidase are dependent on the solvent medium. Chemical communications 2003, (3), 438–439.

117. Fernandez, D. P.; Mulev, Y.; Goodwin, A.; Sengers, J. L., A database for the static dielectric constant of water and steam. Journal of Physical and Chemical Reference Data 1995, 24 (1), 33–70.

118. Karamash, M.; Stumpe, M.; Dengjel, J.; Salgueiro, C. A.; Giese, B.; Fromm, K. M., Reduction Kinetic of Water Soluble Metal Salts by Geobacter sulfurreducens: Fe2+/Hemes Stabilize and Regulate Electron Flux Rates. Frontiers in microbiology 2022, 2135.

119. Amdursky, N.; Pecht, I.; Sheves, M.; Cahen, D., Electron transport via cytochrome C on Si–H surfaces: Roles of Fe and Heme. Journal of the American Chemical Society 2013, 135 (16), 6300–6306.

120. Amdursky, N.; Ferber, D.; Pecht, I.; Sheves, M.; Cahen, D., Redox activity distinguishes solid-state electron transport from solution-based electron transfer in a natural and artificial protein: Cytochrome C and hemin-doped human serum albumin. Physical Chemistry Chemical Physics 2013, 15 (40), 17142–17149.

121. Agam, Y.; Nandi, R.; Kaushansky, A.; Peskin, U.; Amdursky, N., The porphyrin ring rather than the metal ion dictates long-range electron transport across proteins suggesting coherence-assisted mechanism. Proceedings of the National Academy of Sciences 2020, 117 (51), 32260–32266.

122. Blumberger, J., Electron transfer and transport through multi-heme proteins: Recent progress and future directions. Current Opinion in Chemical Biology 2018, 47, 24–31.

123. Futera, Z.; Ide, I.; Kayser, B.; Garg, K.; Jiang, X.; Van Wonderen, J. H.; Butt, J. N.; Ishii, H.; Pecht, I.; Sheves, M., Coherent electron transport across a 3 nm bioelectronic junction made of multi-heme proteins. The journal of physical chemistry letters 2020, 11 (22), 9766–9774.

124. Marcus, R. A., On the theory of oxidation-reduction reactions involving electron transfer. I. The Journal of chemical physics 1956, 24 (5), 966–978.

125. Marcus, R., Discussion comment on mixed reaction-diffusion controlled rates. Discuss Faraday Soc 1960, 29, 21–31.

126. Jiang, X.; Futera, Z.; Ali, M. E.; Gajdos, F.; von Rudorff, G. F.; Carof, A.; Breuer, M.; Blumberger, J., Cysteine linkages accelerate electron flow through tetra-heme protein STC. Journal of the American Chemical Society 2017, 139 (48), 17237–17240.

127. Garg, K.; Ghosh, M.; Eliash, T.; van Wonderen, J. H.; Butt, J. N.; Shi, L.; Jiang, X.; Zdenek, F.; Blumberger, J.; Pecht, I., Direct evidence for heme-assisted solid-state electronic conduction in multi-heme c-type cytochromes. Chemical science 2018, 9 (37), 7304–7310.

128. Blumberger, J., Free energies for biological electron transfer from QM/MM calculation: method, application and critical assessment. Physical Chemistry Chemical Physics 2008, 10 (37), 5651–5667.

129. Phan, H.; Yates, M. D.; Kirchhofer, N. D.; Bazan, G. C.; Tender, L. M.; Nguyen, T.-Q., Biofilm as a redox conductor: a systematic study of the moisture and temperature dependence of its electrical properties. Physical Chemistry Chemical Physics 2016, 18 (27), 17815–17821.

130. Mejía, L.; Kleinekathöfer, U.; Franco, I., Coherent and incoherent contributions to molecular electron transport. The Journal of Chemical Physics 2022, 156 (9), 094302.

131. Alami, F. A.; Soni, S.; Borrini, A.; Nijhuis, C. A., Perspective—Temperature Dependencies and Charge Transport Mechanisms in Molecular Tunneling Junctions Induced by Redox-Reactions. ECS Journal of Solid State Science and Technology 2022, 11 (5), 055005.

132. Sowa, J. K.; Marcus, R. A., On the theory of charge transport and entropic effects in solvated molecular junctions. The Journal of Chemical Physics 2021, 154 (3), 034110.

133. Filman, D. J.; Marino, S. F.; Ward, J. E.; Yang, L.; Mester, Z.; Bullitt, E.; Lovley, D. R.; Strauss, M., Cryo-EM reveals the structural basis of long-range electron transport in a cytochrome-based bacterial nanowire. Communications biology 2019, 2 (1), 1–6.

134. Ru, X. Nanometer to Micrometer Electron Transfer: Incoherent Hopping in Biomolecular Systems. Duke University, 2020.

135. Bortolotti, C. A.; Siwko, M. E.; Castellini, E.; Ranieri, A.; Sola, M.; Corni, S., The reorganization energy in cytochrome c is controlled by the accessibility of the heme to the solvent. The Journal of Physical Chemistry Letters 2011, 2 (14), 1761–1765.

136. Tipmanee, V.; Oberhofer, H.; Park, M.; Kim, K. S.; Blumberger, J., Prediction of reorganization free energies for biological electron transfer: A comparative study of Ru-modified cytochromes and a 4-helix bundle protein. Journal of the American Chemical Society 2010, 132 (47), 17032–17040.

137. Ing, N. L.; El-Naggar, M. Y.; Hochbaum, A. I., Going the distance: long-range conductivity in protein and peptide bioelectronic materials. The Journal of Physical Chemistry B 2018, 122 (46), 10403–10423.

138. Roy, S.; Xie, O.; Dorval Courchesne, N. M., Challenges in engineering conductive protein fibres: Disentangling the knowledge. The Canadian Journal of Chemical Engineering 2020, 98 (10), 2081–2095.

139. Ing, N. L.; Nusca, T. D.; Hochbaum, A. I., Geobacter sulfurreducens pili support ohmic electronic conduction in aqueous solution. Physical Chemistry Chemical Physics 2017, 19 (32), 21791–21799.

140. Yates, M. D.; Golden, J. P.; Roy, J.; Strycharz-Glaven, S. M.; Tsoi, S.; Erickson, J. S.; El-Naggar, M. Y.; Barton, S. C.; Tender, L. M., Thermally activated long range electron transport in living biofilms. Physical Chemistry Chemical Physics 2015, 17 (48), 32564–32570.

141. Bellissent-Funel, M.-C.; Hassanali, A.; Havenith, M.; Henchman, R.; Pohl, P.; Sterpone, F.; Van Der Spoel, D.; Xu, Y.; Garcia, A. E., Water determines the structure and dynamics of proteins. Chemical reviews 2016, 116 (13), 7673–7697.

142. Zahler, C. T.; Shaw, B. F., What are we missing by not measuring the net charge of proteins? Chemistry–A European Journal 2019, 25 (32), 7581–7590.

143. Nenashev, A.; Jansson, F.; Baranovskii, S.; Österbacka, R.; Dvurechenskii, A.; Gebhard, F., Effect of electric field on diffusion in disordered materials. I. One-dimensional hopping transport. Physical Review B 2010, 81 (11), 115203.

144. Jansson, F., Charge transport in disordered materials: simulations, theory, and numerical modeling of hopping transport and electron-hole recombination. 2011.

145. Ghosh, C.; Kara, A.; Rahman, T. S., Usage of pattern recognition scheme in kinetic Monte Carlo simulations: Application to cluster diffusion on Cu (1 1 1). Surface science 2007, 601 (15), 3159–3168.

146. Leetmaa, M.; Skorodumova, N. V., Mean square displacements with error estimates from non-equidistant time-step kinetic Monte Carlo simulations. Computer Physics Communications 2015, 191, 119–124.

147. Byun, H. S.; Pirbadian, S.; Nakano, A.; Shi, L.; El-Naggar, M. Y., Kinetic Monte Carlo simulations and molecular conductance measurements of the bacterial decaheme cytochrome MtrF. ChemElectroChem 2014, 1 (11), 1932–1939.

148. de la Lande, A. Cailliez, F.; Salahub, D. R., Electron transfer reactions in enzymes: seven things that might break down in vanilla marcus theory and how to fix them if they do. Simulating Enzyme Reactivity: Computational Methods in Enzyme Catalysis; Moliner, V., Tunon, I., Eds 2016, 89.

149. Heitele, H., Dynamic solvent effects on electron-transfer reactions. Angewandte Chemie International Edition in English 1993, 32 (3), 359–377.

150. Marcus, R.; Sutin, N., The relation between the barriers for thermal and optical electron transfer reactions in solution. Comments on Inorganic Chemistry 1986, 5 (3), 119–133.

151. Marcus, R.; Sutin, N., Electron-transfer reactions with unusual activation parameters. Treatment of reactions accompanied by large entropy decreases. Inorganic Chemistry 1975, 14 (1), 213–216.

152. Braddock, J. N.; Meyer, T. J., Kinetics of the oxidation of hexaaquoiron (2+) by polypyridine complexes of ruthenium (III). Negative enthalpies of activation. Journal of the American Chemical Society 1973, 95 (10), 3158–3162.

153. Cramer, J. L.; Meyer, T. J., Unusual activation parameters in the oxidation of hexaaquairon (2+) ion by polypyridine complexes of iron (III). Evidence for multiple paths for outer-sphere electron transfer. Inorganic Chemistry 1974, 13 (5), 1250–1252.

154. Kirmaier, C.; Holten, D.; Parson, W. W., Temperature and detection-wavelength dependence of the picosecond electron-transfer kinetics measured in Rhodopseudomonas sphaeroides reaction centers. Resolution of new spectral and kinetic components in the primary charge-separation process. Biochimica et Biophysica Acta (BBA)-Bioenergetics 1985, 810 (1), 33–48.

155. Fleming, G.; Martin, J.; Breton, J., Rates of primary electron transfer in photosynthetic reaction centres and their mechanistic implications. Nature 1988, 333 (6169), 190–192.

156. Shuai, Z.; Geng, H.; Xu, W.; Liao, Y.; André, J.-M., From charge transport parameters to charge mobility in organic semiconductors through multiscale simulation. Chemical Society Reviews 2014, 43 (8), 2662–2679.

157. Waskasi, M. M.; Kodis, G.; Moore, A. L.; Moore, T. A.; Gust, D.; Matyushov, D. V., Marcus bell-shaped electron transfer kinetics observed in an Arrhenius plot. Journal of the American Chemical Society 2016, 138 (29), 9251–9257.

158. Waskasi, M. M.; Newton, M. D.; Matyushov, D. V., Impact of Temperature and Non-Gaussian Statistics on Electron Transfer in Donor–Bridge–Acceptor Molecules. The Journal of Physical Chemistry B 2017, 121 (12), 2665–2676.

159. Matyushov, D. V., Energetics of electron-transfer reactions in soft condensed media. Accounts of chemical research 2007, 40 (4), 294–301.

160. Waskasi, M. M.; Martin, D. R.; Matyushov, D. V., Wetting of the protein active site leads to non-Marcusian reaction kinetics. The Journal of Physical Chemistry B 2018, 122 (46), 10490–10495.

161. Kirchberg, H.; Thorwart, M.; Nitzan, A., Charge Transfer through Redox Molecular Junctions in Nonequilibrated Solvents. The Journal of Physical Chemistry Letters 2020, 11 (5), 1729–1737.

162. Gangopadhyay, D.; Ganguly, B.; Mukherjee, T.; Dutta-Roy, B., Anti-Arrhenius behaviour in positronium chemistry: a Kramers’ turnover? Chemical Physics Letters 2000, 318 (1-3), 161–167.

163. Kramers, H. A., Brownian motion in a field of force and the diffusion model of chemical reactions. Physica 1940, 7 (4), 284–304.

164. Matyushov, D. V., Dynamical effects in protein electrochemistry. The Journal of Physical Chemistry B 2019, 123 (34), 7290–7301.

165. Zitare, U. A.; Szuster, J.; Scocozza, M. F.; Espinoza-Cara, A.; Leguto, A. J.; Morgada, M. N.; Vila, A. J.; Murgida, D. H., The role of molecular crowding in long-range metalloprotein electron transfer: Dissection into site-and scaffold-specific contributions. Electrochimica Acta 2019, 294, 117–125.

166. Matyushov, D. V., Non-Ergodic Electron Transfer in Mixed-Valence Charge-Transfer Complexes. The Journal of Physical Chemistry Letters 2012, 3 (12), 1644–1648.

167. Benderskii, V.; Goldanskii, V.; Makarov, D. E., Quantum dynamics in low-temperature chemistry. Physics reports 1993, 233 (4-5), 195–339.

168. Meisner, J.; Kästner, J., Atom tunneling in chemistry. Angewandte Chemie International Edition 2016, 55 (18), 5400–5413.

169. Dewar, M. J.; Merz, K. M.; Stewart, J. J., Vibrationally assisted tunnelling (VAT) in a 1, 5 hydrogen shift? Journal of the Chemical Society, Chemical Communications 1985, (3), 166–168.

170. Carvalho-Silva, V. H.; Coutinho, N. D.; Aquilanti, V. In Description of deviations from Arrhenius behavior in chemical kinetics and materials science, AIP conference proceedings, AIP Publishing LLC: 2016; p 020006.

171. Murakami, K.; Tanaka, Y.; Sakai, R.; Hisai, Y.; Hayashi, S.; Mizutani, Y.; Higo, T.; Ogo, S.; Seo, J. G.; Tsuneki, H., Key factor for the anti-Arrhenius low-temperature heterogeneous catalysis induced by H+ migration: H+ coverage over support. Chemical Communications 2020, 56 (23), 3365–3368.

