## Supporting Information for "Heme Hopping Falls Short: What Explains Anti-Arrhenius Conductivity in a Multi-heme Cytochrome Nanowire?"

### S1. Methods

#### S1.1. Overview

A general overview of the computations is presented in Figure S1.

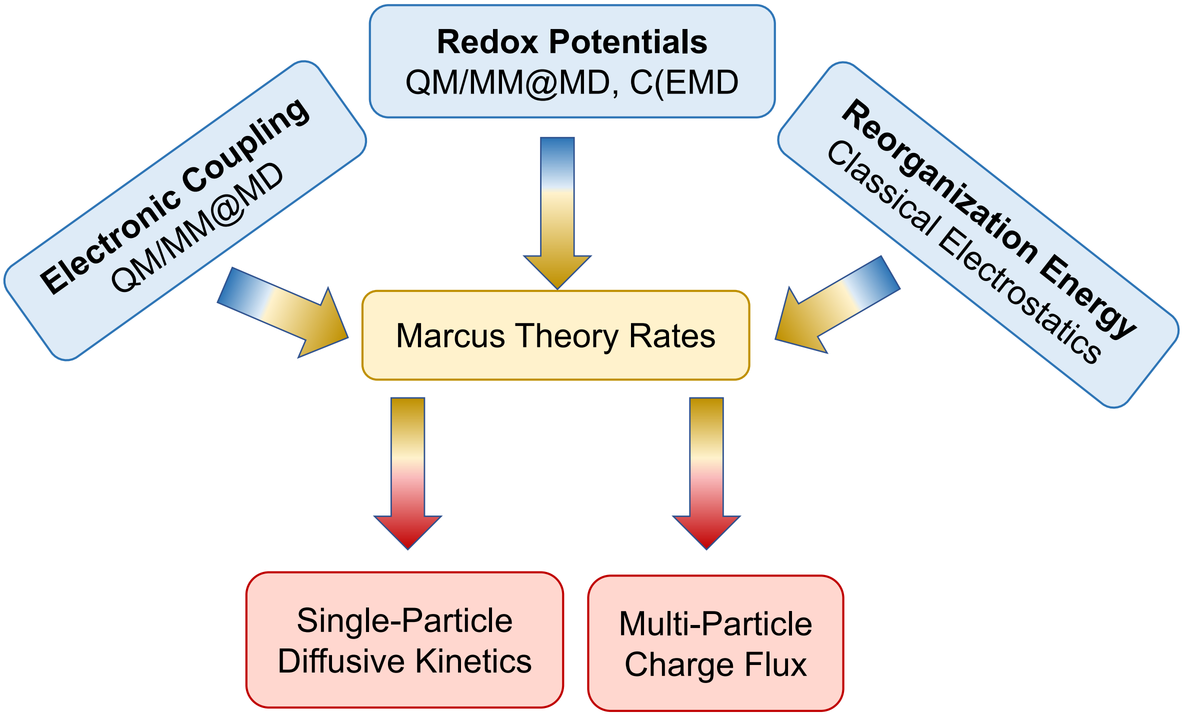

Figure S1. High-level overview of the reported computational Procedure.

To compute the redox potentials, reorganization energies, and electronic couplings, molecular dynamics (MD) simulations were performed. We therefore detail the methodology of the MD simulations first, and then describe the other techniques used in this workflow. The methods used here are compared to those in our prior work[1] in Table S1.

### S1.2. Molecular Dynamics

#### S1.2.1. System setup and parameterization

Starting from the cryo-EM structure of an OmcS filament (PDB accession code 6EF8),[2] a trimeric (chains A, B, and C) assembly was constructed. A segment composed of the first 20 residues of chain D was also included to provide the distal axial histidine (His-16) coordination for heme **#5** (PDB designation HEC-501 in chain B) that is present in the full-length filament. Adopting the chain designations from the PDB, the linear sequence of chains in the trimer structure was C→A→B→D, where the "→" indicates the N- to C-terminal direction.

Each OmcS subunit has 19 Asp, 11 Glu, 5 His (not counting those coordinated to the hemes), 14 Lys, 13 Arg, and 22 Tyr titratable residues. The capping peptide fragment from chain D contributed 2 Glu, 3 His, and 2 Cys residues that were also titratable; in the actual filament context, the His and Cys residues of chain D would be bonded to a heme group. Unless otherwise indicated, the titratable residues were assigned standard protonation states (e.g., deprotonated Asp and Glu; singly protonated His on $N_\epsilon$; protonated Lys, Arg, Cys, and Tyr) to simulate circumneutral pH conditions. The propionic acid groups on the 18 hemes were assumed to be deprotonated, except in simulations where these groups were titrated.

Using tLEaP in the AmberTools20 package[3, 4] hydrogen atoms were added to the OmcS trimer assuming standard protonation of titratable residues. The structure was placed at the center of a box of explicit water with at least a 15 Å buffer region to the boundary of the box. A sufficient number of counterions (9 $Na^+$ per subunit for the fully oxidized state of the heme cofactors with deprotonated propionic acid groups) was added to achieve charge neutrality. Standard proteinogenic residues were modelled with the AMBER FF99SB forcefield.[5] Parameters for the heme cofactor were adopted from Crespo *et al*.[6] and Henriques *et al*.,[7] and used as in previous studies for single- and multi-heme systems.[8, 9] The TIP3P water model[10] and the monovalent ion parameters of Joung and Cheatham[11] were used to model the solution state.

#### S1.2.2. Pre-production simulations

Each solvated structure was subjected to 10,000 steps of steepest descent, followed by 40,000 steps of conjugate gradient minimization. A 10 kcal/(mol Å$^2$) restraint was applied to the heavy atoms of the protein backbone, as well as selected atoms (PDB names FE, NA, NB, NC, ND, C3D, C2A, C3B, C2C, CA, CB) of each heme group during the minimization.

The system was subsequently heated at a rate of 0.3 K/ps to 100, 125, 150, 175, 200, 225, 250, 270, 300, 325, 350, 375, and 400 K in the NVT ensemble and held at the final temperature for 1.0 ns. The restraints on the protein backbone and heme groups were reduced to 1.0 kcal/(mol Å$^2$) for the heating stage. These restraints were further reduced to 0.1 kcal/(mol Å$^2$) for a subsequent 8 ns simulation in the NPT ensemble to equilibrate the density at 1.0 bar and the target temperature. Finally, a 4 ns simulation at the target temperature in the NVT ensemble was performed.

All NVT and NPT simulations (including the production-stage trajectories described below) employed periodic boundary conditions, the Particle Mesh Ewald[12] treatment of electrostatic interactions with a direct sum cut-off of 10.0 Å, the SHAKE algorithm[13, 14] to rigidify bonds to hydrogen atoms, a Langevin thermostat with a collision frequency of 2 $ps^{-1}$, and an integration timestep for the Langevin equation of motion of 2.0 fs. Pressure in NPT simulations was regulated with a Monte Carlo barostat having a relaxation time of 1.0 ps. PMEMD in its CPU and GPU[15] implementations in the Amber20 package[4] was used to perform the minimization and dynamical simulations, respectively.

#### S1.2.3. Production-stage simulations

The dynamics of OmcS were propagated in the NVT ensemble for various redox microstates (i.e., combinations of oxidized and/or reduced heme groups) and finite temperatures (Table S1). In the fully oxidized state of the hemes , the simulations were propagated for 73, 92, 92, 73, 83, 83, 83, 252, 143, 108, 108, 108, and 108 ns at temperatures of 100, 125, 150, 175, 200, 225, 250, 270, 300, 325, 350, 375, and 400 K, respectively. Simulations at 270 K with one heme at a time reduced were propagated for 180 ns for hemes **#1**, **#3**, **#4**, **#5**, and **#6**, and 268 ns for heme **#2**. Simulations at 300 K with one heme at a time reduced were each propagated for 72 ns.

A separate set of simulations were propagated at 300 K in which the hemes in each adjacent pair were assigned a charge distribution halfway between the forcefield definitions for the oxidized and reduced states to model the transition state region of the electron transfer reaction.[16] The simulations for heme pairs **#1-#2**, **#2-#3**, **#3-#4**, **#4-#5**, **#5-#6**, and **#6-#1’** (prime denotes the next subunit in the filament) were propagated for 1.0 ns each, and sampled every 100 fs.

Simulations in which pH- and/or redox-active residues were titrated were also performed (Tables S2, S3). Constant pH simulations were performed at a variety of solution pHs with either all 18 hemes oxidized, or with the 6 central hemes of the trimeric assembly reduced and all other hemes oxidized. Constant redox simulations were performed at a variety of solution potentials in which the redox state of all 18 hemes were simultaneous titrated. Constant redox and pH simulations were performed at pH 7 and a variety of solution potentials in which either the redox state of each heme in the central subunit as well as the protonation state of the propionic acid groups in that subunit were titrated, or all 18 hemes in the trimeric assembly and their propionic acid groups were titrated.

Altogether, 3.0, 0.8, 0.4, and 2.4 μs of production-stage conventional, constant pH, constant redox, and constant pH and redox molecular dynamics, respectively, were performed.

#### S1.3. Quantum Mechanical/Molecular Mechanical Computations at Classical Molecular Dynamics-Generated Configurations (QM/MM@MD)

##### S1.3.1. General Setup

The QM/MM interface to SANDER in the AmberTools20 package[3, 4] was used to generate either Gaussian or Q-Chem input files for selected frames from MD trajectories. Gaussian 16 Rev. A.03[17] was employed for redox potential computations, whereas Q-Chem version 5.3.1[18] was used for an electronic couplings method only implemented in that software package (see below).

The QM region in these calculations (except where indicated) comprised a heme macrocycle, the bonded Cys and His residues up to and including the $C_\beta$ atoms (saturated with a capping hydrogen atom in place of the linkage to $C_\alpha$), and all peripheral substituents, except the propionic acid groups. Capping hydrogen atoms satisfied the valances of the C2A and C3D ring atoms to which the CAA and CAD atoms of the propionic acid groups would have been attached, respectively. When the environment was included, the propionic acid groups were treated with molecular mechanics to avoid having to include their H-bonding partners in the QM region for a balanced description of the system.[19] This definition of the QM region has been used previously.[19] The rest of the simulation cell from the MD simulation, including the protein, water molecules, and counterions were treated with the AMBER FF99SB forcefield, which has been shown to accurately describes *relative* differences in electric fields.[20]

#### S1.3.2. Redox Potentials

#### S1.3.2.1. Background.

The standard redox potential ($E°$) is related to the free energy of reduction ($\Delta G_{red}$) by the thermodynamic relationship

$$\Delta G_{\text{red}} = \text{nF}(\text{E} - E°) \tag{S1}$$

where $\text{n}$ is the number of transferred electrons ($\text{n} = 1$ in this work), $\text{F}$ is Faraday's constant (1 eV/V), $\text{E}$ is the solution potential $\left(\frac{-k_bT}{F} ln[e^-]\right)$, and $E°$ is the standard redox potential $\left(\frac{k_bT}{nF} ln[k_e]\right)$. $k_e$ is the equilibrium constant for the reduction reaction $A_{ox} + ne^- \rightarrow A_{red}$.

Several computational strategies are available for computing $\Delta G_{red}$.[21] In this work we employed the linear response approximation (LRA)[22] in the framework of quantum mechanical/molecular mechanical computations performed at geometries from molecular dynamics (QM/MM@MD).[23] We also used a classical electrostatics-based approach termed Constant Redox Molecular Dynamics (CEMD).[24] The key equations for these previously described approaches are given below for completeness.

#### S1.3.2.1.1. Redox potentials via the linear response approximation

$E°$ was computed as:

$$E° = \frac{-\Delta G_{red}}{nF} + E_{SHE} + E_{EC-FD} = \frac{\Delta G_{ox}}{nF} + E_{SHE} + E_{EC-FD} \quad (S2)$$

where the last two terms are needed to compare the computed potential to an experimental measurement. These terms set the computed potential relative to the standard hydrogen electrode ($E_{SHE} = 4.28\,V$) and correct for the integrated heat capacity and entropy of the electron according to the electron convention and Fermi-Dirac statistics ($E_{EC-FD} = 0.038\,V$).[36, 37]

Under the assumption that the polarization of the solvent (and therefore the free energy of solvation) is a linear function of the charge on the solute, $\Delta G_{ox}$ is given by

$$\Delta G_{ox} = \frac{1}{2}(\langle E_{VertIP}\rangle_{red} + \langle E_{-VertEA}\rangle_{ox}) \quad (S3)$$

where $\langle E_{VertIP}\rangle_{red}$ is the vertical ionization potential thermally averaged over configurations with the solute in the reduced state. $\langle E_{-VertEA}\rangle_{ox}$ is the negative of the vertical electron affinity computed over thermally averaged configurations with the solute in the oxidized state. Each of these terms is given by $E_{ox}(r_X) - E_{red}(r_X)$, which are the energies of the oxidized and reduced solute, respectively, at the geometries of the $X = ox$ state for $\langle E_{-VertEA}\rangle_{ox}$ and $X = red$ state for $\langle E_{VertIP}\rangle_{red}$. The LRA is valid if the distributions for $\langle E_{-VertEA} \rangle_{ox}$ and $\langle E_{VertIP} \rangle_{red}$ are symmetric, Gaussian-shaped, and of equal width (Tables S4-S7, Figures S2-S4).[22]

In practical computations, a differential free energy of solvation term is needed in principle to correct vertical energies obtained from finite-sized simulations.[25] However, we found that replacement of the finite-sized explicit solvent region with a bulk implicit solvent only caused a mean unsigned shift in the computed redox potentials of 0.040 or 0.048 (Table S8), depending on whether a 15 or 150 mM salt concentration was specified in computations using the Poisson-Boltzmann Solvation Area model[26] implemented in AmberTools20.[3, 4].

##### S1.3.2.1.2. Redox Potentials from classical electrostatics

$\Delta G_{\text{red}}$ can be split into contributions from electrostatic ($\Delta G_{elec}$) and non-electrostatic ($\Delta G_{non-elec}$) terms. If $\Delta G_{\text{red}}$ is computed relative to a reference compound for which $\Delta G_{\text{red}}$ is known, and it is assumed that $\Delta G_{non-elec}$ is the same for both the target and reference compounds, then

$$\Delta G_{red} = \text{nF}\left(E - E^{\circ}_{ref}\right) + \Delta G_{elec} - \Delta G_{elec,ref} \qquad (S4)$$

where $E^{\circ}_{ref}$ is the standard redox potential of the reference compound and $\Delta G_{elec,ref}$ is pre-computed to reproduce $E^{\circ}_{ref}$. Thus, $\Delta G_{red}$ is determined from the free energy change in electrostatic interactions for the two redox states ($\Delta G_{elec}$).

The implementation of this approach in AMBER is dubbed Constant Redox Molecular Dynamics and uses *N*-acetomicroperoxidase-11 as the reference compound for a *c*-type heme.[24] The methodology, paired with the analogous constant pH approach (i.e., Constant Redox and pH Molecular Dynamics, C(E,pH)MD) was used extensively in the present work.

**S1.3.2.2. Computational Details**

Vertical energy gaps were assessed for configurations sampled from MD trajectories of each heme in the oxidized or reduced state as detailed in Tables S4-S7 and Figures S2-S5). Figure S6 shows the redox potentials computed systematic from every possible pairing of vertical ionization and electron affinities obtained from multiple trajectories at 270 K for a given heme.

The SCF energies used to compute the vertical energy gaps were obtained with the B3LYP approximate density functional[27-29] and a mixed basis set. The LANL2DZ effective core potential and valance basis sets[30] were used to describe the Fe center of the heme cofactor, whereas the 6-31G(D) basis set[31-35] was applied to all first and second row (H, C, N and S) atoms. This mixed basis set was referred to as the double-ζ basis in the article (Tables S5-S7, Figures S2-S5). For a subset of calculations, the valance part of the mixed basis set was improved from double- to triplet-ζ quality (i.e., LANL2TZ[30, 36] and 6-311G(d)[33, 37, 38]), and this combination was denoted as the triple-ζ basis (Table S4, Figure S3). With respect to redox potentials, the mean unsigned difference between B3LYP using the double- or triple-ζ basis sets was 0.034 V (Table S4, S5), which was well within the standard error of the mean for either set of calculations.

To estimate the effective static dielectric constant of each protein binding site, the redox potential for each heme was computed from a representative sub-ensemble of frames in the protein environment (Table S6, Figure S3) and in a variety of homogeneous solvents using a polarizable continuum model.[39] B3LYP/double-ζ was used for these computations.

The B3LYP functional used for all these computations has been extensively validated for the description of the geometrical,[40-42] electronic,[43-45] reactivity,[41, 46] and electromagnetic response[40, 46-48] properties of iron porphyrins and heme cofactors. We nonetheless evaluated the ability of several functional and basis set combinations to reproduce the experimental gas-phase vertical ionization energy for iron(II) octaethylporphyrin (Fe(II)OEP).

The first vertical ionization energy for Fe(II)OEP was measured by photoelectron spectroscopy to be 6.06 eV.[49] B3LYP with the double- or triple-ζ basis sets well reproduced this value, predicting it to be 5.99 or 5.89 eV, respectively. To compute this quantity, the ethyl substituents were truncated to methyl groups and the resulting Fe(III)OMP (M = methyl) was optimized in the low spin (singlet) or high spin (triplet) states. Consistent with experimental and theoretical work,[50] B3LYP with either basis set found the triplet to be the ground state. The ionized product was optimized in the low spin (doublet) or high spin (quartet) states, and the latter was found to be of lower energy; again, in agreement with prior work.[50] The vertical ionization energy was finally computed as the transformation from the triplet ground state of Fe(II)OMP to the quartet ground state of Fe(III)OMP at the geometry of the triplet Fe(II)OMP.

The level of agreement to experiment achieved with B3LYP and either the double- (< 0.07) or triple-ζ (< 0.17 eV) basis set is consistent with a recent benchmarking study on the electronic component of computed redox potentials for iron complexes,[51] which found B3LYP to be among the top ten best-performing functionals. The computed value is also consistent with prior theoretical work on Fe(II) porphyrins that found first ionization energies at 6.29, 5.97, and 5.50 eV for Fe(II)P, Fe(II)TPP, and Fe(II)OEP, respectively.[50, 52] We note that repetition of the computation with CAM-B3LYP (5.87 eV) or ωB97XD (5.88 eV) with the double-ζ basis set showed no substantial improvement over B3LYP.

Interestingly, we found that the 0.07 eV underestimation of the vertical ionization energy for Fe(II)OMP by B3LYP/double-ζ was partly compensated for by the overestimation of the vertical ionization energy for the heme cofactor when the geometry was directly taken from MD trajectories without QM relaxation (Table S9). Upon partial geometry optimization with all dihedral angles fixed to the values obtained from MD, the average vertical ionization energy for each heme—whether computed for configurations from the fully oxidized or single-heme-reduced trajectories—was lowered by 0.005 to 0.030 eV. A larger decrease in the vertical ionization energies may be obtained if a full geometry optimization was performed, but the relaxation would have to take place within the protein binding site to prevent collapse of the various conformations sampled during the dynamics into a handful of local minima. For the calculations reported in Table S9, the optimizations needed to be torsionally constrained because they were performed in vacuum.

To summarize, the unoptimized geometries from the classical dynamics using the AMBER parameterization for the heme group led to a systematic overestimation of the vertical ionization energies at the B3LYP/double-ζ level of theory that was partially offset by the tendency of the B3LYP/double-ζ methodology to underestimate the vertical ionization energy, as shown by for the related system Fe(II)OMP.

#### S1.3.3. Electronic Couplings

Electronic couplings for every adjacent pair of heme groups in OmcS were computed with the diabatic state based Absolutely Localized Molecular Orbital Multi-State Density Functional Theory 2 (ALMO(MSDFT2))[53] approach, where the "2" indicates the second-generation scheme for evaluating the off-diagonal elements of the diabatic Hamiltonian. The ALMO(MSDFT2) method, as implemented in Q-Chem[18] version 5.3.1, was used with the Perdew-Burke-Ernzerhof (PBE) approximate density functional[54] and the def2-SVP basis set.[55]

Benchmark calculations with this methodology on cationic dimers of ethylene, cyclopropene, cyclobutadiene, cyclopentadiene, furane, pyrrole, thiophene, and imidazole, each at inter-molecular separations of 3.5, 4.0, 4.5, and 5.0 Å gave couplings linearly correlated ($R^2$ = 0.9946 with the high level *ab initio* results of the Hab11 dataset (Table S10).[56] The ALMO(MSDFT2) couplings captured at least 82% of the reported *ab initio* values for inter-molecular separations ≥ 4.0 Å. Curiously, the linear correlation and agreement with the Hab11 dataset was significantly worse in the case of an acetylene dimer at all inter-molecular separations (Table S10). Underestimation in the couplings was also worse at distances < 4.0 Å for all dimer pairs.

The average ratio of the reference Hab11 dataset couplings to ALMO(MSDFT2) couplings for all examined dimer pairs except acetylene at inter-molecular separations ≥ 4.0 Å was 1.11, which is comparable to scaling factors of 0.72 to 1.54 reported for fragment-orbital (FO-) and constrained (C-) DFT methods.[56] Because of convergence issues with the charge flux model discussed below, this scaling factor was not used in the kinetic modeling. Our conclusions are not affected by this choice.

The electronic couplings were averaged over 50 snapshots sampled at a rate of 100 fs/frame from production-stage MD trajectories. In these trajectories, the partial atomic charges of the donor and acceptor hemes were scaled to be halfway between the forcefield definitions for the reduced and oxidized states,[16] while all other hemes were in the oxidized state. A separate trajectory mimicking the transition state region for the heme-to-heme electron transfer in this fashion was conducted for every unique donor-acceptor heme pair. The influence of including an electrostatic environment or the heme propionic acid groups in the QM region was assessed (Table S11), and both factors were found to have a minimal effect. The temporal coherence was also assessed by comparing root mean-squared and root squared-mean electronic couplings (Table S12).[57]

#### S1.4. Reorganization Energy

#### S1.4.1. Background.

The reorganization energy for electron transfer from site $m$ to $n$ was decomposed into additive inner- ($\lambda_{inner}$) and outer- ($\lambda_{outer}$) sphere contributions.[58] $\lambda_{inner}$ was defined as the energy needed to rearrange the nuclei to be consistent with a different charge state while remaining in the original electronic state. $\lambda_{outer}$ reflected instead the energy penalty for rearranging nuclei in the environment in response to altered electrostatic interactions with the electron donor and acceptor. Prior theoretical work[58, 59] found the inner sphere component for models of a bis-histidine ligated iron porphyrin to be 0.05-0.085 eV, which is at least 5 times less than the smallest outer-sphere component computed in the present work.

$\lambda_{outer}$ was computed according to linear response theory from thermal averages of the vertical energy gaps in the initial ($\langle \Delta E_{\mathrm{outer}} \rangle_i$) and final $\left(\langle \Delta E_{\mathrm{outer}} \rangle_f\right)$ states (Eq. S5).[58]

$$\lambda_{\mathrm{outer}}^{st} = \frac{1}{2}\left(\langle \Delta E_{\mathrm{outer}} \rangle_i - \langle \Delta E_{\mathrm{outer}} \rangle_f\right) \qquad (S5)$$

The superscript "st" indicates that this definition of $\lambda_{outer}$ is also known as the Stokes shift reorganization energy. The name derives from the fact that $\lambda_{outer}^{st}$ is one-half the horizontal displacement between the minima on parabolic free energy curves for the initial and final states. From an optical spectroscopy point of view, those lowest energy configurations correspond to absorption and emission maxima of charge-transfer bands.[60] The spectroscopic Stokes shift measures the difference between these peak maxima.

The linear response approximation to $\lambda_{outer}$ given in Eq. S5 is rigorous when the fluctuations in the vertical energy gaps (see below definition) are Gaussian distributed and ergodically sampled according to Boltzmann statistics on the timescale of the electron transfer.[19] In this case, the variances of the vertical energy gap in the initial $(\sigma_i^2)$ and final $(\sigma_f^2)$ states are equal and give an equivalent definition for $\lambda_{outer}$ given in Eq. S6.

$$\lambda_{outer}^{var} = \frac{\sigma_i^2 + \sigma_f^2}{4k_bT} \qquad (S6)$$

The "var" superscript emphases that this definition is based on the variance of the vertical energy gaps. $\lambda_{outer}^{var}$ describes the curvature of the parabolic free energy curves for the initial and final states, and from the spectroscopic point of view, it is related to the inhomogeneous broadening of charge transfer bands.[60]

The assumption of ergodic sampling on the electron transfer timescale was assessed by comparing $\lambda_{outer}^{var}$ to $\lambda_{outer}^{st}$. If $\frac{\lambda_{outer}^{var}}{\lambda_{outer}^{st}} \approx 1$, then ergodic sampling was observed.[19]

The vertical energy gaps needed for both $\lambda_{outer}^{st}$ and $\lambda_{outer}^{var}$ were computed by Eq. S7

$$\Delta E_{outer} = E_f(R^N) - E_i(R^N) - \left(\mathrm{I}E_{Donor} + \mathrm{I}E_{Acceptor}\right) \qquad (S7)$$

where $E_f(R^N)$ and $E_i(R^N)$ are the total electrostatic potential energies at the same nuclear configuration $(R^N)$ for the final and initial charge states, respectively. The $\mathrm{I}E$ terms (defined in Eq. S8) are the self-energy contributions of the donor or acceptor to the change in total electrostatic potential energy $\left(E_f(R^N) - E_i(R^N)\right)$, which belong to $\lambda_{inner}$ and are therefore subtracted out in the computation of $\Delta E_{outer}$. Note that the electrostatic interaction energy between the donor and acceptor is still included.

$$\mathrm{IE} = \sum_{j \neq k} \frac{q_{j,O} q_{k,O} - q_{j,R} q_{k,R}}{R_{jk}} \qquad (S8)$$

In Eq. S8, the $q$ terms are the atomic partial charges on the $j$th and $\mathrm{k}$th atoms of the heme cofactor in the oxidized (O) or reduced (R) charge states, and $R_{jk}$ is the distance between these atoms.

The $E(R^N)$ and $IE$ terms of Eq. S7, and thereby the two definitions of $\lambda_{outer}$ in Eqs. S5 and S6 were computed at the classical (molecular) mechanics level with the ESANDER facility in CPPTRAJ[61] of the AmberTools20 package.[3, 4]

#### S1.4.2. Computational Details

An example input script to compute reorganization energy using CPPTRAJ is provided below.

**Example CPPTRAJ script for the computation of electron transfer reorganization energies**

```
#Definition of Constants ========================================
kb= 0.00008617333262145
T= 300
#================================================================

# Load topology (prmtop) and trajecotyr (mdcrd) for the reduced state of
# Heme #1 (resid 1280) with all other hemes oxidized
parm   ../../../1280/omcs_r1280.prmtop
trajin ../../../1280/prod_2.mdcrd 1 1800 1 parmindex 0

# Compute total electrostatic energy of the system with Heme #1 reduced and
# all other hemes oxidized
esander I1280 out I1280.dat cut 10.0 igb 0 ntb 1 ntf 2 ntc 2
run

# Now, strip the trajectory of everything except either the donating (Heme
# #1; resid 1280) or the accepting (Heme #2; resid 1274) atoms, and then re-
# compute the electrostatic energy
strip !(:HEH&:1280) outprefix ../../../1280/DAsubsysA1280
esander DADAsubsysA1280 out DAsubsysA1280.dat cut 10.0 igb 0 ntb 1 ntf 2 ntc
2
run

unstrap

strip !(:HEH&:1274) outprefix ../../1280/DAsubsysA1274
esander DADAsubsysA1274 out DAsubsysA1274.dat cut 10.0 igb 0 ntb 1 ntf 2 ntc
2
run

unstrap

# Compute the electrostatic interaction energy between the donor-acceptor
# pair and the environment
sysA = I1280[elec]
subAA = DADAsubsysA1280[elec]
subAB = DADAsubsysA1274[elec]
PairIntA = (sysA[elec] - subAA[elec] - subAB[elec])
```

```
writedata If.dat sysA[elec] subAA[elec] subAB[elec] PairIntA
run
#----------------------------------------------------------------------

# Reload the same trajectory, but now with the topology for Heme #2 (resid
# 1274) reduced and all other hemes oxidized
clear trajin
parm   ../../../1274/omcs_r1274.prmtop
trajin ../../../1280/prod_2.mdcrd 1 1800 1 parmindex 1

# In analogy to the above commands: Compute the electrostatic energy for the
# full system in this redox microstate, then strip all but the donor or
# acceptor atoms from the trajectory and re-compute the energy, and finally,
# compute the electrostatic interaction energy between the donor-acceptor
# pair and the environment
esander F1274 out F1274.dat cut 10.0 igb 0 ntb 1 ntf 2 ntc 2
run

strip !(:HEH&:1280) outprefix ../../../1280/DAsubsysB1280
esander DADAsubsysB1280 out DAsubsysB1280.dat cut 10.0 igb 0 ntb 1 ntf 2 ntc
2
run

unstrap

strip !(:HEH&:1274) outprefix ../../1280/DAsubsysB1274
esander DADAsubsysB1274 out DAsubsysB1274.dat cut 10.0 igb 0 ntb 1 ntf 2 ntc
2
run

unstrap

# Compute the electrostatic interaction energy between the donor-acceptor
# pair and the environment
sysB = F1274[elec]
subBA = DADAsubsysB1280[elec]
subBB = DADAsubsysB1274[elec]
PairIntB = (sysB[elec] - subBA[elec] - subBB[elec])
writedata Ib.dat sysB[elec] subBA[elec] subBB[elec] PairIntB
run
```

```
# Compute the vertical energy gap as the difference in the electrostatic
# interaction energy of the donor-acceptor pair with the environment in the
# OR and RO (R = reduced; O = oxidized) charge states. The obtained vertical
# energy gap is for the “forward” RO → OR reaction. The factor “0.043”
# converts energies in kcal/mol to eV.
VEGf = (PairIntB - PairIntA) * 0.0434
writedata VEGf.dat PairIntA PairIntB VEGf
#======================================================================
========

# Repeat all of the above steps for the “backward” OR → RO reaction.
clear trajin
trajin ../../../1274/prod_2.mdcrd 1 1800 1 parmindex 1
esander I1274 out I1274.dat cut 10.0 igb 0 ntb 1 ntf 2 ntc 2
run

strip !(:HEH&:1280) outprefix ../../../1274/DAsubsysC1280
esander DADAsubsysC1280 out DAsubsysC1280.dat cut 10.0 igb 0 ntb 1 ntf 2 ntc
2
run

unstrap

strip !(:HEH&:1274) outprefix ../../1274/DAsubsysC1274
esander DADAsubsysC1274 out DAsubsysC1274.dat cut 10.0 igb 0 ntb 1 ntf 2 ntc
2
run

unstrap

# Compute the electrostatic interaction energy between the donor-acceptor
# pair and the environment
sysC = I1274[elec]
subCA = DADAsubsysC1280[elec]
subCB = DADAsubsysC1274[elec]
PairIntC = (sysC[elec] - subCA[elec] - subCB[elec])
writedata Ff.dat sysC[elec] subCA[elec] subCB[elec] PairIntC
run
#----------------------------------------------------------------------

clear trajin
```

```
trajin ../../1274/prod_2.mdcrd 1 1800 1 parmindex 0
esander F1280 out F1280.dat cut 10.0 igb 0 ntb 1 ntf 2 ntc 2
run

strip !(:HEH&:1280) outprefix ../../../1274/DAsubsysD1280
esander DADAsubsysD1280 out DAsubsysD1280.dat cut 10.0 igb 0 ntb 1 ntf 2 ntc
2
run

unstrap

strip !(:HEH&:1274) outprefix ../../1274/DAsubsysD1274
esander DADAsubsysD1274 out DAsubsysD1274.dat cut 10.0 igb 0 ntb 1 ntf 2 ntc
2
run

unstrap

# Compute the electrostatic interaction energy between the donor-acceptor
# pair and the environment
sysD = I1274[elec]
subDA = DADAsubsysD1280[elec]
subDB = DADAsubsysD1274[elec]
PairIntD = (sysD[elec] - subDA[elec] - subDB[elec)
writedata Ff.dat sysD[elec] subDA[elec] subDB[elec] PairIntD
run
.
VEGb = (PairIntD - PairIntC) * 0.0434
writedata VEGb.dat PairIntC PairIntD VEGb
#============================================================
========

# Compute the thermal averages of the “forward” and “backward” reaction
# vertical energy gaps
avgVEGf   = avg(VEGf)
avgVEGb   = avg(VEGb)

# Compute the Stokes reorganization energy. Note that the definitions of the
# initial and final states to compute avgVEGb were inverted from the
# definitions used to compute avgVEGf. That is, the initial state for the
# forward reaction was Heem #1 reduced and Heme #2 oxidized, whereas the
```

```
# initial state in the backward direction was Heme #1 oxidized and Heme #2
# reduced. Thus, for lambdast, we are adding a negative avgVEGb instead of
# subtracting a positive avgVEGb.
lambdast  = (avgVEGf + avgVEGb)/2

# Compute the variance reorganization energy
varVEGf   = stdev(VEGf)^2
varVEGb   = stdev(VEGb)^2
lambdavarf = varVEGf/(2*kb*T)
lambdavarb = varVEGb/(2*kb*T)

# Compute the ergodicity factor
xg = (lambdavarf + lambdavarb)/(2*lambdast)

printdata avgVEGf avgVEGb varVEGf varVEGb
printdata lambdast lambdavarf lambdavarb xg
writedata Reorg1280,1274.dat lambdast lambdavarf lambdavarb xg
run
```

### S1.5. Marcus Theory Rate Constants

The reaction and reorganization free energies and the electronic couplings were used to compute Marcus theory rate constants in the high temperature limit (Eq. S9). The rate constant is given by a Boltzmann activation factor (ρ) scaled by a tunneling probability (κ).[62] ρ describes the likelihood of thermal fluctuations bring the donor and acceptor to an energetically degenerate configuration, whereas κ gives the likelihood of an electron tunneling from the donor to the acceptor in that configuration.

$$k_{n,m} = \kappa\rho = \left(\frac{2\pi\langle H_{n,m}\rangle^2}{\hbar}\right)\left(\frac{e^{\frac{-\left(\Delta G^{\circ}_{n,m}+\lambda_{n,m}\right)^2}{4\lambda_{n,m}k_bT}}}{\sqrt{\left(4\pi\lambda_{n,m}k_bT\right)}}\right) \qquad (S9)$$

In Eq. S9, $\hbar$ is the reduced Plank constant; $\langle H_{m,n}\rangle^2$ is the squared-mean electronic coupling between site *m* and site *n*; $\Delta G_{m,n}$ and $\lambda_{m,n}$ are the reaction and reorganization free energies for electron transfer from site *m* to *n*; $k_b$ is the Boltzmann constant, and $T$ is absolute temperature. Depending on the regime of weak or strong coupling fluctuations, the squared-mean ($\langle H_{DA}\rangle^2$) or the mean-squared ($\langle H_{DA}^2\rangle$) electronic coupling, respectively, should be used in Eq. S9.[57] The coherence parameter, which is the ratio of these quantities (i.e., $R_{coh} = \frac{\langle H_{DA}\rangle^2}{\langle H_{DA}^2\rangle}$), characterizes the strength of coupling fluctuations. $R_{coh} \approx 1$ (Table S12), indicating that the weak coupling fluctuation regime was operative in OmcS, and justified the use of $\langle H_{n,m}\rangle^2$ in Eq. S9.

S1.6. Multi-Step Kinetic Modeling

S1.6.1. Single-Particle Diffusion on a one-dimensional periodic chain

A charge hopping current at a given applied bias is expressed in terms of a diffusion constant by Eq. S10.

$$\mathrm{I} = \left(\frac{Ae^2\rho D}{k_b TL}\right)\mathrm{V} = \left(\frac{e^2 N_{chg} D}{k_b TL^2}\right)\mathrm{V} \tag{S10}$$

Eq. S10 is derived from Ohm's law by expressing the resistance in terms of the cross-sectional area ($A$) and the length ($L$) of the conduction path, the mobile charge density ($\rho$) in the filament, and the diffusion constant ($D$) of the charge carriers. Because $\rho$ is the number of mobile charges ($N_{chg}$) per unit volume ($v$), and $v = LA$, the last expression on the right-hand-side was obtained by substituting $A\rho = A\left(\frac{N_{chg}}{v}\right) = \left(\frac{N_{chg}}{L}\right)$. Since the method used to compute $D$ (see below) neglects interactions between mobile charges, precedent[3] was followed in assuming every other heme was occupied, which provides an upper limit to the computed current.

To compute the diffusion coefficient, Derrida's analytical solution to the one-dimensional periodic chain was solved using the implemented by Jansson *et al.*[63, 64]

$$\mathrm{D} = \frac{\Delta x^2}{\left(\sum_{n=1}^{N} r_n\right)^2}\left(A\sum_{n=1}^{N} u_{n\sum_{i=1}^{N} i r_{n+1}} + N\sum_{n=1}^{N} k_{n+1,n} u_n r_n\right) - \mathrm{A}\Delta x^2\,\frac{N+2}{2} \tag{S10}$$

$\Delta x$ is the minimum edge-to-edge macrocycle distance[65] averaged over all heme pairs in an OmcS subunit (5.2 Å); $N$ is the number of hopping sites in a repeat unit of the periodic chain (the 6 hemes in a subunit of the filament); $A$, $u_n$, and $r_n$ are related to products of site-to-site rate ratios (Eqs. S11-S13), where the rate from site *m* to *n* ($k_{n,m}$) was obtained from non-adiabatic Marcus theory in the high temperature limit.

$$\mathrm{A} = \frac{N}{\sum_{n=1}^{N} r_n}\left(1 - \prod_{n=1}^{N} \frac{k_{n,n+1}}{k_{n+1,n}}\right) \qquad (S11)$$

$$u_n = \frac{1}{k_{n+1,n}}\left(1 + \sum_{i=1}^{N-1} \prod_{j=1}^{i} \frac{k_{n-j,n+1-j}}{k_{n+1-j,n-j}}\right) \qquad (S12)$$

$$r_n = \frac{1}{k_{n+1,n}}\left(1 + \sum_{i=1}^{N-1} \prod_{j=1}^{i} \frac{k_{n+j,n+j+1}}{k_{n+j+1,n+j}}\right) \qquad (S13)$$

#### S1.6.2. Multi-Particle steady state hopping on a one-dimensional chain

The steady-state flux model developed by Blumberger and co-workers[66] was also used to assess charge hopping conductivity. Electron injection and rejection rates (assumed to be the same) gave a maximal or protein-limited flux when set to $10^{10}$ electrons/s, as used in prior work[67] and confirmed in Figure S2. All other input parameters (i.e., Marcus theory rates) were identical to those used for the single-particle diffusion model. The program for the flux model was kindly provided by Xiuyun Jiang.

#### S1.6.3. Comparison of diffusive and stead-state flux electrical hopping currents

The maximum protein-limited electron flux through an OmcS trimer was converted into a current by multiplying by the electrical charge: $I = -eJ$, where $e$ = 1.602 × $10^{-19}$

C/electron and $J$ is the electron flux (electrons/s). This intrinsic current was realized (Figure S7) by setting the rate of electron injection and ejection (assumed to be equal) to $10^{10}$ electrons/s. Using the experimental resistance of OmcS, scaled for the trimer model (226 MΩ/subunit × 3 subunits = 678 MΩ),[1] the injection/ejection rate corresponded to a voltage of 1.1 V. Our intention was not to model heterogeneous electron transfer. Rather, this voltage was used to relate the results from both kinetic schemes: Eq S10 was evaluated with the voltage (1.1 V) and the length of the filament (L = $4.67 \times 10^{-7}$ cm/subunit * 3 subunits) used for the flux model. Both theoretical results were compared to what the current would be if this voltage was applied across an OmcS trimer having the experimentally measured conductivity; we refer to this current as "experimental" with quotation marks in the main text because it has not actually been measured. Of utmost importance, please note that the voltage is just a multiplicative factor used to set the protein-limited charge flux, the diffusion constant, and the experimental conductivity on an equal footing for comparison.

### S1.7. H-bond Analysis

The intra-protein H-bonding network in fully oxidized OmcS was examined in MD simulations at temperatures of 100, 125, 150, 175, 200, 225, 250, 270, 300, 325, 350, and 400 K. The analysis used both the H-bond plugin to Visual Molecular Dynamics (VMD)[68] implemented by J. C. Gumbart and Dong Luo, as well as the H-bond facility in CPPTRAJ.[61]

These programs were found to give essentially the same result (Table S17) if care was taken to use a consistent Donor-Hydrogen-Acceptor (D-H-A) angle. VMD identifies H-bonds with a D-H-A angle < 20°, whereas CPPTRAJ considers a H-bond to have an A-H-D angle > 135°; for comparability, 135° must be changed to 160° (the complementary angle to 20°). VMD also allows any heavy atom that satisfies the geometrical criteria to be considered a H-bond donor/acceptor, whereas CPPTRAJ follows the FON convention (i.e., a H-bond donor/acceptor is an F, O, or N atom). The latter point is the reason for the small differences in Table S17.

To compare the composition of the H-bonding network at two different temperatures, a bash script written and distributed by Emmett Leddin was used (https://emleddin.github.io/comp-chem-website/Analysisguide-hba.html). To compute the characteristic H-bonding Frequency (CHD), the difference CHF (ΔCHF), and the ΔCHF only consisting of H-bonds that exist at 300 K and some other temperature ($\Delta CHF_{shared}$), the Pandas and NumPy python libraries were used to create and analyze pivot tables.

**S2. Assessment of Convergence on Redox Potentials at 270 K**

There is no available experimental data with which to compare the computed $E°$ values at low temperature. More extensive sampling at 270 K was therefore performed to ensure the computational results were robust. Multiple trajectories for the oxidized and reduced states of each heme were generated by periodically randomizing the velocities. $E°$ values were compared between and over all these trajectories. A total of 1.2 μs of dynamics and ~7200 QM/MM single point calculations were used to compute the final $E°$ values at 270 K (Table S7; Figures S4-S6).

The cooling-induced shifts were considered converged because (Figure S4-S6): (1) The average $E°$ computed from the last pair of trajectories in the oxidized and reduced states agreed with the average over all trajectories within 0.031 V; (2) The $E°$ computed from the first pair and the last pair of trajectories from the two redox states typically agreed within 0.042 V, and (3) The vertical energy gaps before and after ≥100 ns of dynamics during which the velocities were randomized multiple times agreed within 0.060 V. Exceptions to the two latter observations only pertain to heme **#5**.

The $E°$ for heme **#5** became more negative by 0.117 V between the first pair and the last pair of trajectories from the two redox states. The larger negative shift in $E°$ for this heme primarily resulted from a decrease in the magnitude of the vertical electron affinity of the oxidized state by as much as 0.255 eV (Figure S5). To account for this fact, the oxidized state dynamics were propagated and sampled for QM/MM energy evaluations until (1) There was no longer a monotonic change in the vertical energy gap, and (2) The change in the vertical energy gap between the last two sets of trajectories was only 0.029 V. The relatively strong stabilization of the oxidized state for heme **#5** at 270 versus 300 K had an electrostatic origin, as explained in §2.3.1 of the main text.

Table S1. Comparison of the QM/MM@MD methodologies employed in the present and prior work[1]

| Present Work | Dahl *et al.* |
|---|---|
| **Initial Structure** | |
| PDB 6EF8[2] | PDB 6EF8[2] |
| **Oligomerization state of model** | |
| Trimer | Monomer[a] or Dimer |
| **MD unit cell dimensions** | |
| 102 x 101 x 193 | 90 x 90 x 140 Å |
| **Protonation state of ionizable residues** | |
| Standard at pH 7 | Standard for pH 7[b] |
| **Protein & cofactor force field** | |
| AMBER99SB | CHAMM36[c,d] |
| **Water & counterion force field** | |
| TIP3P | TIP3P |
| **Minimization** | |
| • 50,000 steps Trimer<br>• First 10,000 steps by steepest descent<br>• Remaining 40,000 by conjugate gradient<br>• Harmonic restraints with a force constant of 10 kcal/(mol*Å$^2$) on protein backbone and select heme atoms (AMBER mask syntax: @CA,C,O,N\|(:HEH,PRN)@FE,NA,NB, NC,ND,C3D,C2A,C3B,C2C,CA,CB) | Minimization + 2.5 ns relaxation of solvent at target temperature |
| **Thermalization** | |
| • 2.0 ns<br>• NVT ensemble<br>• 2 fs time step<br>• 0.3 K/ps from 0 to target temperature (100, 125, 150, 175, 200, 225, 250, 270, 300, 325, 350, and 400 K)<br>• Held at target temperature for remaining 1.0 ns<br>• Temperature scaling using Langevin dynamics with a 5 ps$^{-1}$ collision frequency<br>• Hydrogen atoms restrained with the SHAKE algorithm<br>• Harmonic restraints with a force constant of 1.0 kcal/(mol*Å$^2$) on protein backbone and select heme atoms (AMBER mask syntax: | • 3.5 ns<br>• NVT ensemble<br>• Temperature = 270 or 310 K<br>• Harmonic restraints of 0.1 kcal/(mol*Å$^2$)[e] to amino acid side chains; 1.0 kcal/(mol*Å$^2$) to protein backbone and hemes |

@CA,C,O,N|(:HEH,PRN)@FE,NA,NB,
NC,ND,C3D,C2A,C3B,C2C,CA,CB)

**Density Equilibration**

- Part I: 4.0 ns
  - NVT ensemble
  - 2 fs time step
  - Temperature maintained at target temperature using Langevin dynamics with a 2 $ps^{-1}$ collision frequency
  - Hydrogen atoms restrained with the SHAKE algorithm
  - Harmonic restraints with a force constant of 0.1 kca/(mol*$Å^2$) on protein backbone and select heme atoms (AMBER mask syntax: @CA,C,O,N|(:HEH,PRN)@FE, NA,NB,NC,ND,C3D,C2A,C3B,C 2C,CA,CB)
- Part II: 8.0 ns
  - NPT ensemble
  - 2 fs time step
  - 1.0 bar maintained using a Monte Carlo barostat with a pressure relaxation period of 1.0 ps
  - Temperature maintained at target value using Langevin dynamics with a 2 $ps^{-1}$ collision frequency
  - Hydrogen atoms restrained with the SHAKE algorithm Harmonic restraints with a force constant of 0.1 kcal/(mol*$Å^2$) on protein backbone and select heme atoms (AMBER mask syntax: @CA,C,O,N|(:HEH,PRN)@FE, NA,NB,NC,ND,C3D,C2A,C3B,C 2C,CA,CB)

**Production Stage**

- Redox Microstates:
  - All Oxidized:
    - 100 K: 73 ns
    - 125 K: 92 ns
    - 150 K: 92 ns

- 50 ns (presumably each microstate)
- Langevin thermostat at target temperature

| | |
|---|---|
| ▪ 175 K: 73 ns<br>▪ 200 K: 83 ns<br>▪ 225 K: 83 ns<br>▪ 250 K: 83 ns<br>▪ 270 K: 252 ns<br>▪ 300 K: 72 ns<br>▪ 325 K: 108 ns<br>▪ 350 K: 108 ns<br>▪ 375 K: 108 ns<br>▪ 400 K: 108 ns<br>o Single Heme Reduced (@300/@270 K)<br>▪ Heme #1: 72/180 ns<br>▪ Heme #2: 72/232 ns<br>▪ Heme #3: 72/180 ns<br>▪ Heme #4: 72/180 ns<br>▪ Heme #5: 72/180 ns<br>▪ Heme #6: 72/180ns<br>o For timings of constant pH, constant redox or constant redox and pH MD simulations, see Tables S2 and S3.<br>• NVT ensemble<br>• 2 fs time step<br>• Temperature maintained at target value using Langevin dynamics with a 2 $ps^{-1}$ collision frequency<br>• Hydrogen atoms restrained with the SHAKE algorithm | |
| **QM Region** | |
| A single heme macrocycle, the bonded Cys and His residues up to and including the $C_\beta$ atoms (saturated with a capping hydrogen atom in place of the linkage to $C_\alpha$), and all peripheral substituents, except the propionic acid groups. Capping hydrogen atoms satisfied the valances of the C2A and C3D ring atoms to which the CAA and CAD atoms of the propionic acid groups would have been attached, respectively. | Same as present work except the propionic acid groups were included. |
| **QM Functional** | |
| B3LYP | ωB97X-D |
| QM Basis Set | |
| • Double-ζ: Fe = LANL2DZ; H, C, N, S = 6-31G(d) | Fe: LANL2DZ; H, C, N, O, S = cc-pVDZ |

| | |
|---|---|
| • Triple-ζ: Fe = LANL2TZ; H, C, N, S = 6-311G(d) | |
| Number of Frames Analyzed | |
| • 300 K<br> o Triple-ζ = 916 (Table S5)<br> o Double-ζ = 1187 (Table S6)<br>• 270 K<br> o Double-ζ = 1874 (Table S7) | 175 (25 frames per redox microstate for 7 redox microstates) [f] |

[a]Though mentioned, no data was presented for the monomer, or the impact of oligomerization from monomer to dimer on any property.
[b]Claimed that standard protonation states were assigned because the experimental structure was solved at pH 10.5, but to be consistent with experiment, the protonation state of Tyr and Lys residues should have been checked.
[c]Not well documented in the original publication, the simulations were performed by loading CHARMM22 followed by CHARMM36 into NAMD, where the older version of the force field was needed for some bonded parameters in the heme group. This procedure is very risky because any parameters or dihedral multiplicities defined in CHARMM22 *but not* CHARMM36 would be retained by NAMD, thereby performing the simulations with an unvalidated admixture of the two generations of the forcefield. The handful of needed CHARMM22 parameters should have been simply appended to the CHARM36 parameter set.
[d]The heme parameterization used from Ref. [69] was for a histidine and methionine, not a bis-histidine ligated heme as in OmcS. The patch for the histidine ligation was simply applied twice without further validation. Some deficiencies in the heme force field were apparent in the simulations (i.e., the failure to reproduce the relative orientation of the axial ligands for Heme #4, which was achieved with the AMBER FF99SB force field in the present work).
[e]Wrongly given as kcal/mol in original publication
[f]This is for the electron-hopping regime, which is comparable to the work presented by us. Another 175 frames were examined for the hole-hopping regime.

Table S2. Length (in ns) of constant pH molecular dynamics simulations performed before and after a six-electron reduction of the hemes in the central subunit of the simulated filament

| Solution pH | Fully Oxidized State | Fully Reduced State |
|---|---|---|
| -8 | 26.5 | 18.70 |
| -6 | 27.0 | 18.70 |
| -4 | 27.2 | 18.50 |
| -2 | 26.2 | 18.30 |
| 0 | 27.2 | 18.20 |
| 2 | 26.8 | 18.30 |
| 4 | 26.9 | 18.60 |
| 6 | 22.8 | 19.70 |
| 8 | 29.2 | 19.70 |
| 10 | 27.5 | 18.40 |
| 12 | 28.9 | 19.80 |
| 14 | 29.9 | 19.40 |
| 16 | 24.8 | 21.60 |
| 18 | 32.9 | 22.60 |
| 20 | 33.1 | 20.40 |
| 22 | 22.1 | 22.80 |
| 24 | 26.7 | 22.50 |

Table S3. Length (in ns) of constant redox or constant redox and pH molecular dynamics simulations for the hemes in a trimeric assembly of OmcS titrated either separately or simultaneously at various solution potentials and pH 7

| Solution Potential | Individual Hemes Titrated[a] | | | | | | All Hemes Titrated[b] | All Hemes Titrated[a,c] | | |
|---|---|---|---|---|---|---|---|---|---|---|
| | #1 | #2 | #3 | #4 | #5 | #6 | | Rep. 1 | Rep. 2 | Rep. 3 |
| -0.030 | 33 | 41 | 42 | 58 | 39 | 50 | 72 | 43 | 14 | 13 |
| -0.090 | 33 | 44 | 42 | 58 | 33 | 54 | 55 | 35 | 20 | 25 |
| -0.150 | 35 | 40 | 38 | 58 | 33 | 51 | 66 | 35 | 18 | 13 |
| -0.210 | 37 | 36 | 38 | 50 | 35 | 46 | 52 | 33 | 18 | 11 |
| -0.270 | 37 | 38 | 43 | 57 | 35 | 49 | 51 | 34 | 23 | 16 |
| -0.330 | 43 | 39 | 44 | 58 | 35 | 58 | 52 | 36 | 16 | 24 |
| -0.390 | 43 | 39 | 43 | 59 | 55 | 55 | 60 | 44 | 30 | 14 |

[a]The protonation states of the propionic acid groups on all 18 hemes in the trimeric assembly were also titrated at a solution pH of 7

[b]All propionic acid groups were locked in the deprotonated state

[c]Three independent replicas (Rep.) of these simulations were performed

Table S4. Statistics for QM/MM vertical energy gaps and redox potentials computed with a triple-ζ basis set on configurations sampled from classical dynamics at 300 K

| Oxidized State Trajectory | | | | Reduced State Trajectory | | | | |
|---|---|---|---|---|---|---|---|---|
| # of Replicas | Time (ns) | # of Frames | -ΔVEA | # of Replicas | Time (ns) | # of Frames | ΔVIP | $E^{\circ}$ |
| | | | | Heme #1 | | | | |
| 1 | 36-72 | 100 | 3.4394 ±0.2617 | 1 | 36-72 | 107 | 5.072 ±0.2294 | -0.063 ±0.0347 |
| | | | | Heme #2 | | | | |
| 1 | 36-72 | 154 | 3.3495 ±0.2162 | 1 | 36-72 | 150 | 5.0663 ±0.2187 | -0.111 ±0.025 |
| | | | | Heme #3 | | | | |
| 1 | 36-72 | 131 | 3.6084 ±0.2059 | 1 | 36-72 | 130 | 4.8879 ±0.2198 | -0.071 ±0.026 |
| | | | | Heme #4 | | | | |
| 1 | 36-72 | 128 | 3.4472 ±0.2675 | 1 | 36-72 | 120 | 4.6957 ±0.2328 | -0.248 ±0.032 |
| | | | | Heme #5 | | | | |
| 1 | 36-72 | 139 | 3.382 ±0.2359 | 1 | 36-72 | 168 | 4.9026 ±0.2300 | -0.177 ±0.027 |
| | | | | Heme #6 | | | | |
| 1 | 36-72 | 112 | 3.4707 ±0.2712 | 1 | 36-72 | 87 | 4.9771 ±0.2383 | -0.095 ±0.036 |

Table S5. Statistics for QM/MM vertical energy gaps and redox potentials computed with a double-ζ basis set on configurations sampled from classical dynamics at 300 K

| Oxidized State Trajectory | | | | Reduced State Trajectory | | | | |
|---|---|---|---|---|---|---|---|---|
| # of Replicas | Time (ns) | # of Frames | -ΔVEA | # of Replicas | Time (ns) | # of Frames | ΔVIP | $E^{\circ}$ |
| Heme #1 | | | | | | | | |
| 1 | 36-72 | 174 | 3.3667 ±0.2649 | 1 | 36-72 | 178 | 5.0323 ±0.2475 | -0.119 ±0.027 |
| Heme #2 | | | | | | | | |
| 1 | 36-72 | 167 | 3.3306 ±0.2154 | 1 | 36-72 | 173 | 5.0467 ±0.2157 | -0.130 ±0.023 |
| Heme #3 | | | | | | | | |
| 1 | 36-72 | 158 | 3.5604 ±0.2033 | 1 | 36-72 | 161 | 4.8614 ±0.2082 | -0.108 ±0.023 |
| Heme #4 | | | | | | | | |
| 1 | 36-72 | 167 | 3.4035 ±0.2572 | 1 | 36-72 | 167 | 4.6920 ±0.2318 | -0.271 ±0.027 |
| Heme #5 | | | | | | | | |
| 1 | 36-72 | 174 | 3.3367 ±0.2281 | 1 | 36-72 | 175 | 4.8740 ±0.235 | -0.214 ±0.025 |
| Heme #6 | | | | | | | | |
| 1 | 36-72 | 159 | 3.4338 ±0.244 | 1 | 36-72 | 159 | 4.9475 ±0.2483 | -0.128 ±0.028 |

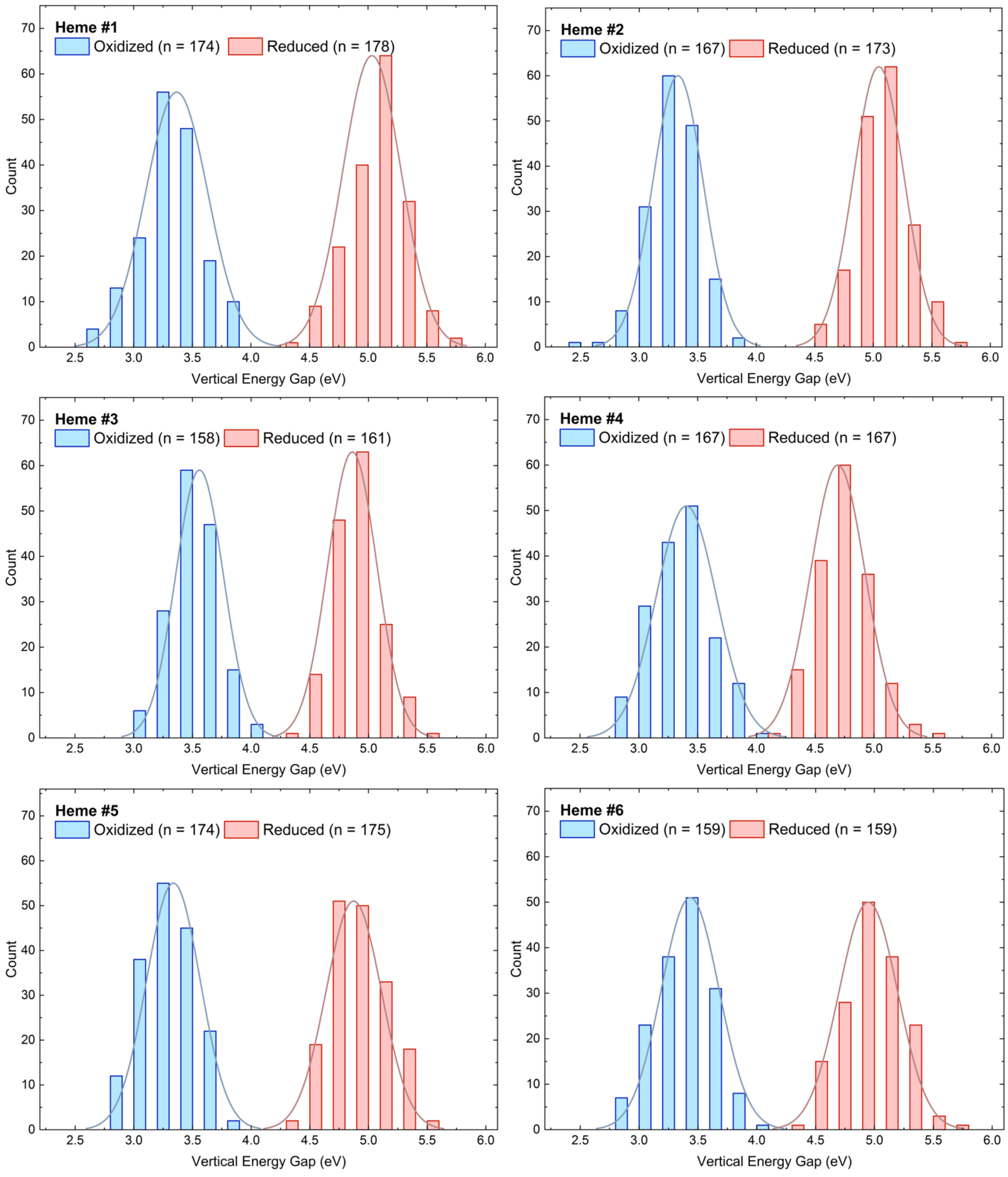

Figure S2. Distribution of vertical energy gaps for each heme in the central subunit of the trimeric OmcS system from trajectories in which the heme was oxidized (blue) or reduced (red) at 300 K. The vertical energy gaps were computed with B3LYP/[Fe = LANL2DZ; H, C, N, S = 6-31G(d)].

Table S6. Statistics for QM/MM vertical energy gaps and redox potentials computed with a double-ζ basis set on a sub-ensemble of configurations sampled from classical dynamics at 300 K

| Oxidized State Trajectory | | | | Reduced State Trajectory | | | | |
|---|---|---|---|---|---|---|---|---|
| # of Replicas | Time (ns) | # of Frames | -ΔVEA | # of Replicas | Time (ns) | # of Frames | ΔVIP | $E°$ |
| | | | | Heme #1 | | | | |
| 1 | 67-72 | 26 | 3.3757 ±0.0484 | 1 | 67-72 | 26 | 5.0466 ±0.0454 | -0.108 ±0.066 |
| | | | | Heme #2 | | | | |
| 1 | 67-72 | 26 | 3.3045 ±0.0406 | 1 | 67-72 | 26 | 4.9575 ±0.0374 | -0.188 ±0.055 |
| | | | | Heme #3 | | | | |
| 1 | 67-72 | 21 | 3.6012 ±0.0450 | 1 | 67-72 | 24 | 4.8036 ±0.0465 | -0.117 ±0.065 |
| | | | | Heme #4 | | | | |
| 1 | 67-72 | 25 | 3.4972 ±0.0463 | 1 | 67-72 | 23 | 4.6901 ±0.0425 | -0.225 ±0.063 |
| | | | | Heme #5 | | | | |
| 1 | 67-72 | 26 | 3.438 ±0.0398 | 1 | 67-72 | 25 | 4.8275 ±0.0431 | -0.186 ±0.059 |
| | | | | Heme #6 | | | | |
| 1 | 67-72 | 25 | 3.464 ±0.0349 | 1 | 67-72 | 24 | 4.9364 ±0.0394 | -0.119 ±0.053 |

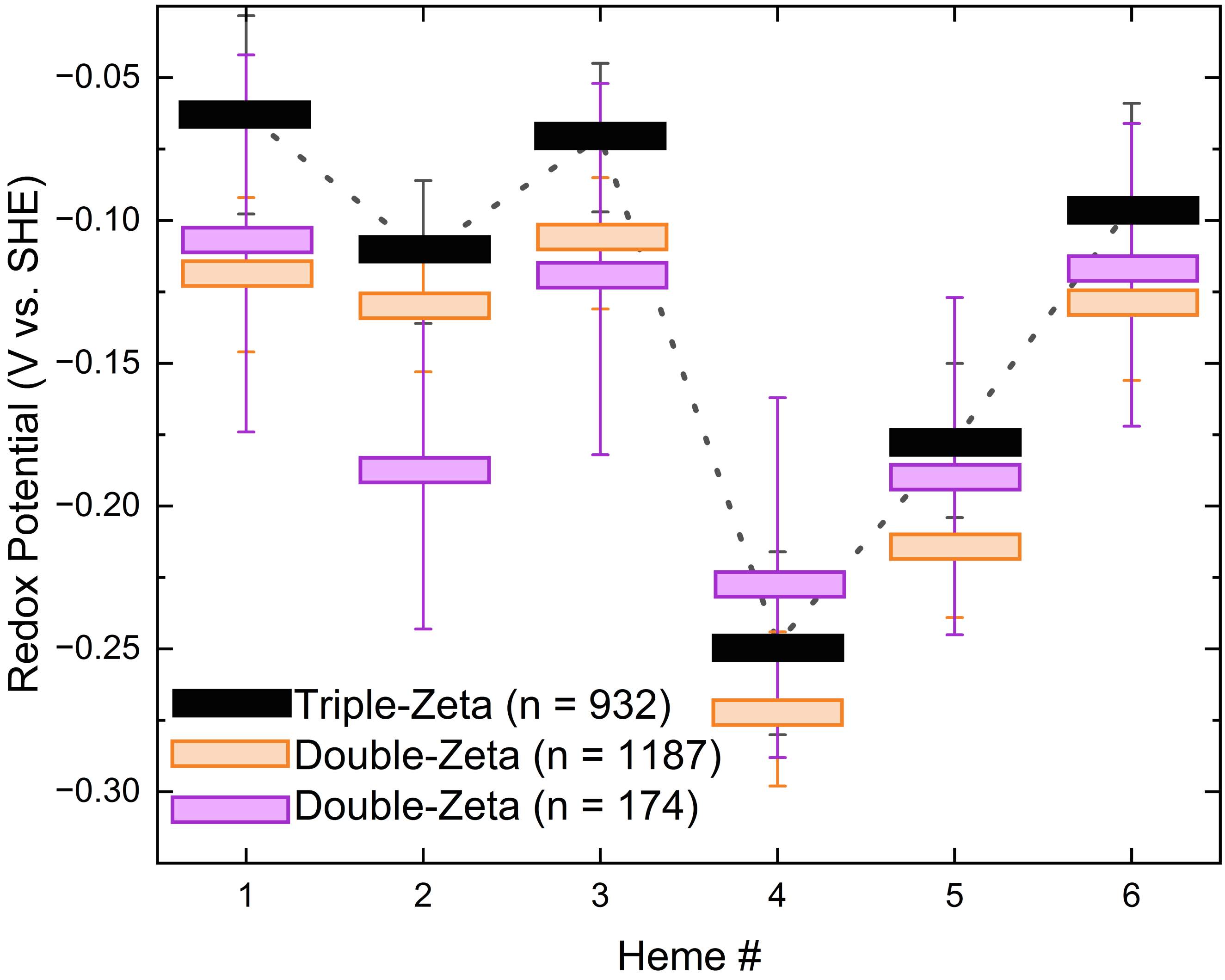

Figure S3. Comparison of computed redox potentials using a triple- *versus* double-ζ basis set, and the double-ζ basis set for the full set of examined snapshots or a subset used for mechanistic analyses. “n = …” indicates the number of frames analyzed for the oxidized and single-heme reduced states to compute the redox potentials.

Table S7. Statistics for QM/MM vertical energy gaps and redox potentials computed with a double-ζ basis set on configurations sampled from classical dynamics at 270 K

| Oxidized State Trajectory | | | | Reduced State Trajectory | | | | |
|---|---|---|---|---|---|---|---|---|
| # of Replicas | Time (ns) | # of Frames | -ΔVEA | # of Replicas | Time (ns) | # of Frames | ΔVIP | $E°$ |
| Heme #1 | | | | | | | | |
| 1 | 36-72 | 176 | 3.253 ±0.019 | 1 | 36-72 | 168 | 5.040 ±0.018 | |
| 2 | 108-144 | 76 | 3.262 ±0.025 | 2 | 108-144 | 75 | 5.052 ±0.027 | |
| 3 | 145-180 | 26 | 3.182 ±0.042 | 3 | 145-180 | 25 | 4.953 ±0.053 | |
| 4 | 180-216 | 47 | 3.255 ±0.031 | | | | | |
| Overall | | | 3.238 ±0.061 | | | | 5.015 ±0.062 | -0.192 ±0.087 |
| Heme #2 | | | | | | | | |
| 1 | 36-72 | 176 | 3.304 ±0.017 | 1 | 36-72 | 176 | 4.964 ±0.016 | |
| 2 | 108-144 | 76 | 3.322 ±0.026 | 2 | 108-144 | 74 | 4.985 ±0.027 | |
| 3 | 145-180 | 26 | 3.245 ±0.042 | 3 | 145-180 | 26 | 5.068 ±0.031 | |
| | | | | 4 | 181-216 | 26 | 5.159 ±0.031 | |
| | | | | 5 | 217-232 | 31 | 5.022 ±0.034 | |
| Overall | | | 3.290 ±0.052 | | | | 5.039 ±0.064 | -0.154 ±0.082 |
| Heme #3 | | | | | | | | |
| 1 | 36-72 | 180 | 3.531 ±0.014 | 1 | 36-72 | 174 | 4.825 ±0.015 | |
| 2 | 108-144 | 72 | 3.492 ±0.026 | 2 | 108-144 | 72 | 4.872 ±0.026 | |
| 3 | 145-180 | 25 | 3.474 ±0.032 | 3 | 145-180 | 26 | 4.806 ±0.033 | |
| Overall | | | 3.499 ±0.044 | | | | 4.835 ±0.045 | -0.152 ±0.063 |
| Heme #4 | | | | | | | | |
| 1 | 36-72 | 172 | 3.369 ±0.017 | 1 | 36-72 | 174 | 4.614 ±0.016 | |
| 2 | 108-144 | 72 | 3.330 ±0.026 | 2 | 108-144 | 71 | 4.577 ±0.031 | |

| | | | | | | | | |
|---|---|---|---|---|---|---|---|---|
| 3 | 145-180 | 25 | 3.378 ±0.047 | 3 | 145-180 | 26 | 4.592 ±0.044 | |
| Overall | | | 3.359 ±0.056 | | | | 4.594 ±0.056 | -0.342 ±0.079 |
| Heme #5 | | | | | | | | |
| 1 | 36-72 | 175 | 3.341 ±0.016 | 1 | 36-72 | 175 | 4.831 ±0.015 | |
| 2 | 108-144 | 71 | 3.216 ±0.029 | 2 | 108-144 | 74 | 4.818 ±0.024 | |
| 3 | 145-180 | 26 | 3.167 ±0.038 | 3 | 145-180 | 25 | 4.823 ±0.041 | |
| 4 | 180-216 | 51 | 3.086 ±0.033 | | | | | |
| 5 | 217-252 | 50 | 3.115 ±0.030 | | | | | |
| Overall | | | 3.185 ±0.067 | | | | 4.824 ±0.050 | -0.315 ±0.084 |
| Heme #6 | | | | | | | | |
| 1 | 36-72 | 172 | 3.304 ±0.017 | 1 | 36-72 | 176 | 4.975 ±0.015 | |
| 2 | 108-144 | 72 | 3.255 ±0.026 | 2 | 108-144 | 76 | 4.994 ±0.023 | |
| 3 | 145-180 | 25 | 3.195 ±0.050 | 3 | 145-180 | 24 | 4.941 ±0.038 | |
| 4 | 180-216 | 50 | 3.140 ±0.036 | | | | | |
| 5 | 217-252 | 50 | 3.267 ±0.028 | | | | | |
| Overall | | | 3.232 ±0.074 | | | | 4.970 ±0.047 | -0.218 ±0.088 |

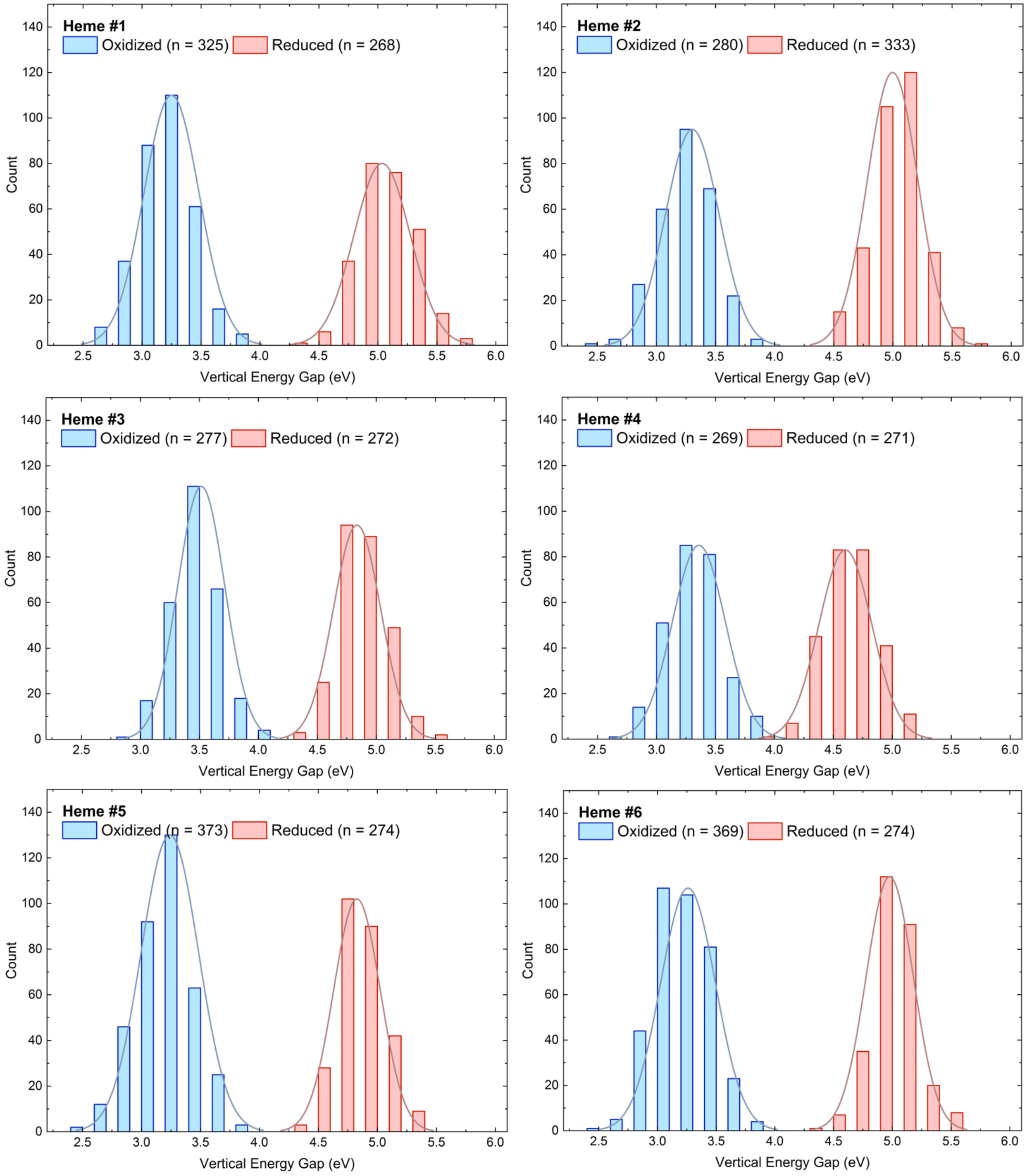

Figure S4. Distribution of vertical energy gaps for each heme in the central subunit of the trimeric OmcS assembly from trajectories in which the heme was oxidized (blue) or reduced (red) at 270 K. The vertical energy gaps were computed with B3LYP/[Fe = LANL2DZ; H, C, N, S = 6-31G(d)].

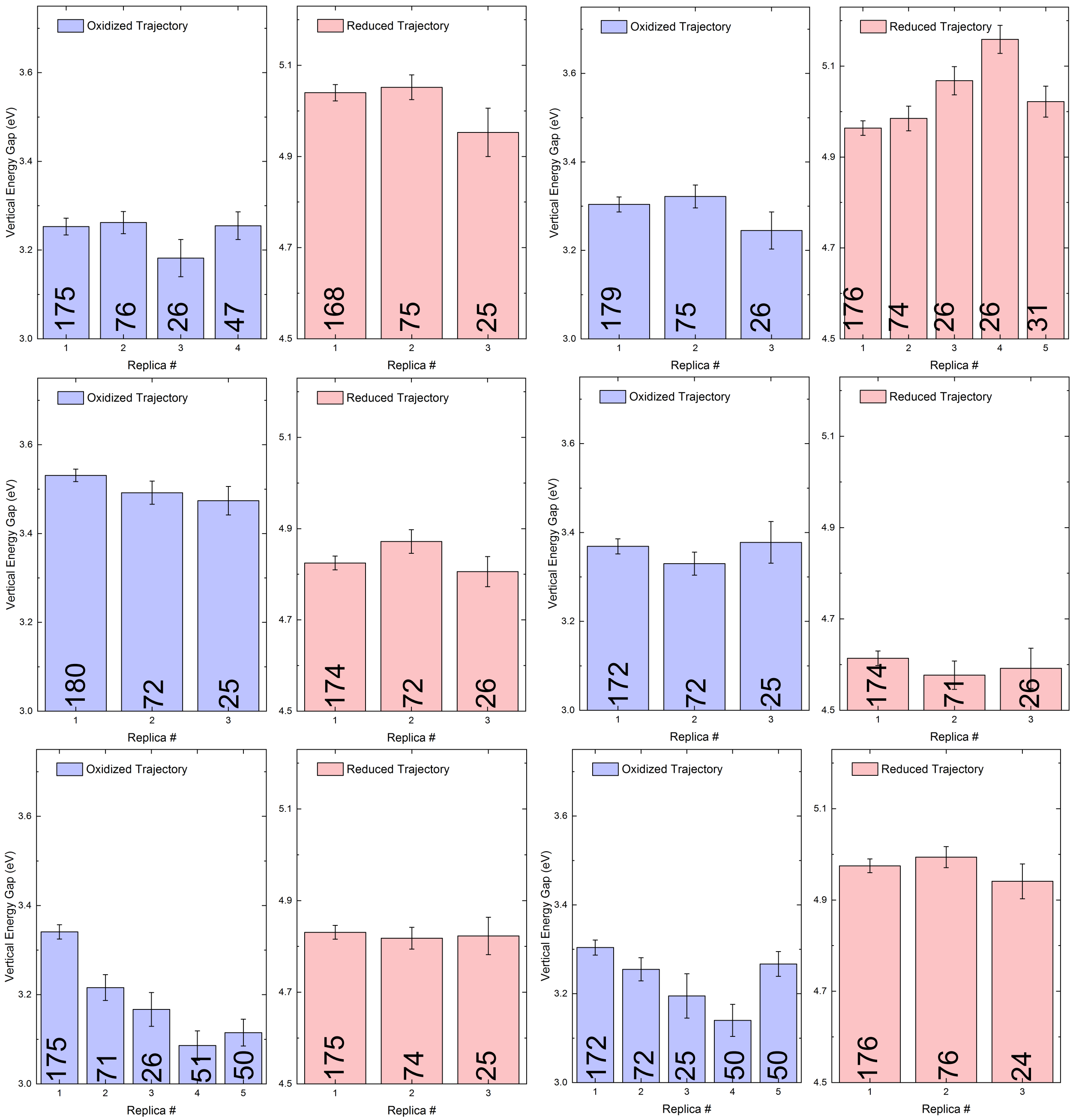

Figure S5. Trajectory-to-trajectory variation in vertical energy gaps computed at the DFT level on configurations sampled from classical dynamics. The vertical energy gaps were computed with B3LYP/[Fe = LANL2DZ; H, C, N, S = 6-31G(d)]. The number of frames sampled from each trajectory is indicated on the corresponding bar.

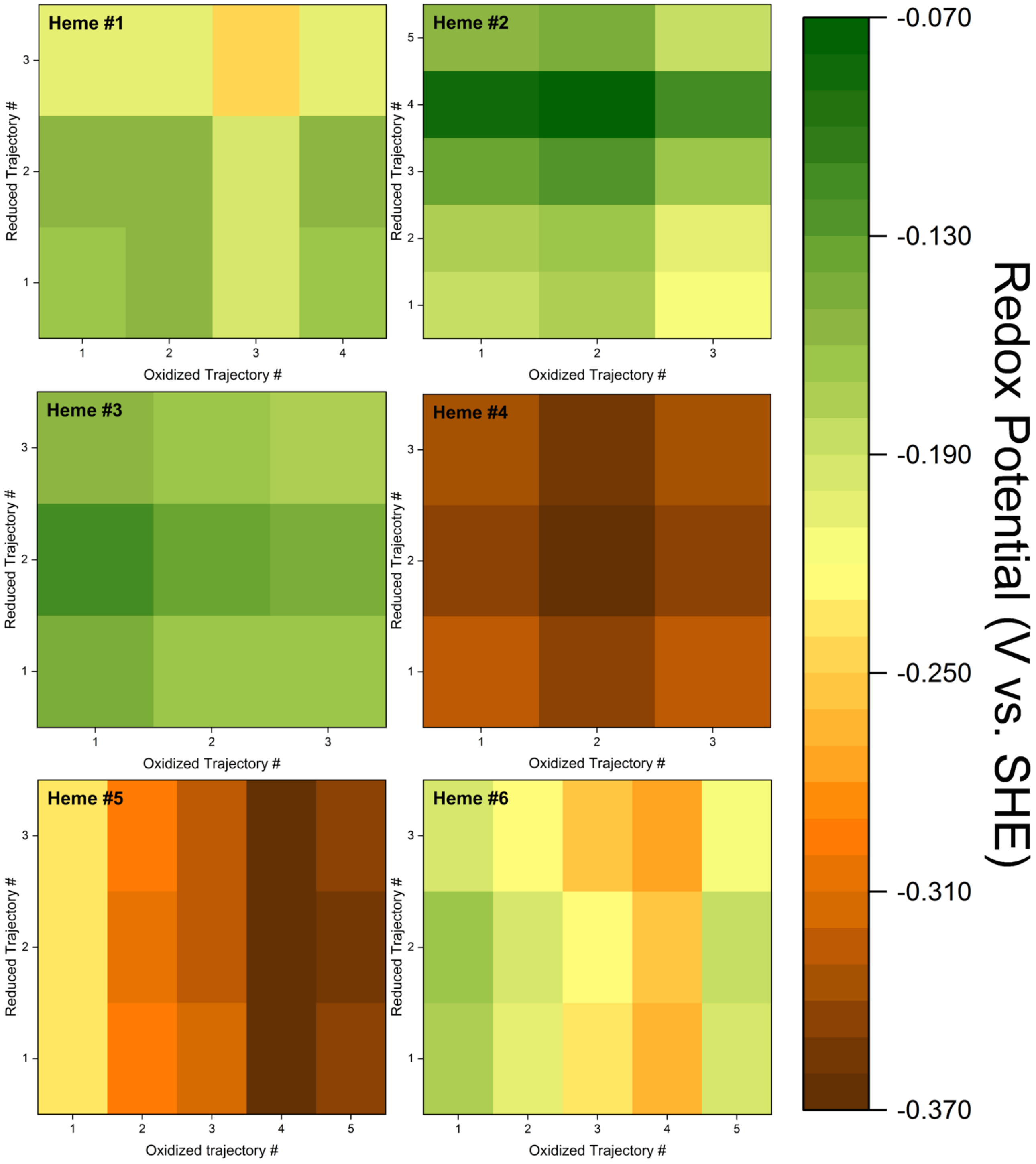

Figure S6. Distribution of redox potentials combinatorically computed from every pairing of an oxidized- and a reduced-state trajectory for each heme. The trajectories of a given redox state were propagated in chronological order starting from the last configuration of the previous simulation, but with randomized velocities.

Table S8. Influence of explicit and implicit water/ion models on heme redox potentials in OmcS

| Heme # | Explicit Water/Ions | Implicit Water/Ions (15 mM salt) | Implicit Water/Ions (150 mM salt) |
|---|---|---|---|
| 1 | -0.109 ± 0.013 | -0.124 ± 0.014 | -0.136 ± 0.009 |
| 2 | -0.184 ± 0.011 | -0.107 ± 0.008 | -0.109 ± 0.008 |
| 3 | -0.126 ± 0.012 | -0.134 ± 0.011 | -0.133 ± 0.009 |
| 4 | -0.225 ± 0.012 | -0.261 ± 0.013 | -0.274 ± 0.011 |
| 5 | -0.185 ± 0.011 | -0.262 ± 0.009 | -0.263 ± 0.008 |
| 6 | -0.121 ± 0.010 | -0.097 ± 0.012 | -0.080 ± 0.010 |

Table S9. Change in the vertical energy gap before and after torsionally restrained optimizations in vacuum of the heme groups from MD snapshots

| | Oxidized State Trajectory | | | Reduced Trajectory | | |
|---|---|---|---|---|---|---|
| Heme # | # of Frames | Before Opt. | After Opt. | # of frames | Before Opt. | After Opt. |
| 1 | 171 | 4.880 ±0.007 | 4.867 ±0.006 | 162 | 4.899 ±0.007 | 4.877 ±0.006 |
| 2 | 82 | 4.874 ±0.009 | 4.845 ±0.007 | 75 | 4.897 ±0.012 | 4.877 ±0.010 |
| 3 | 135 | 4.827 ±0.007 | 4.821 ±0.006 | 138 | 4.848 ±0.007 | 4.832 ±0.006 |
| 4 | 62 | 4.923 ±0.011 | 4.894 ±0.010 | 82 | 4.929 ±0.011 | 4.905 ±0.011 |
| 5 | 85 | 4.854 ±0.008 | 4.824 ±0.008 | 81 | 4.815 ±0.008 | 4.797 ±0.006 |
| 6 | 57 | 4.927 ± 0.014 | 4.905 ±0.013 | 58 | 4.928 ±0.015 | 4.918 ±0.014 |

Table S10. Comparison of Absolutely Localized Molecular Orbital Multi-State Density Functional Theory 2 (ALMO(MSDFT2)) Electronic Couplings (in meV) to Previously Reported Multireference Configuration Interaction with Quadruples (MRCI+Q) values[a]

| | Distance (Å) | | | |
|---|---|---|---|---|
| Compound | 5.0 | 4.5 | 4.0 | 3.5 |
| acetylene | 3.494 | 10.166 | 27.790 | 72.335 |
| | (56.600) | (114.800) | (231.800) | (460.700) |
| ethylene | 46.546 | 110.970 | 237.824 | 490.041 |
| | (68.500) | (137.600) | (270.800) | (519.200) |
| cyclopropene | 45.177 | 108.289 | 244.423 | 537.712 |
| | (54.000) | (118.400) | (254.000) | (536.600) |
| cyclobutadiene | 39.060 | 91.797 | 194.955 | 406.350 |
| | (62.200) | (121.700) | (239.100) | (462.700) |
| cyclopentadiene | 40.454 | 95.365 | 199.956 | 417.804 |
| | (53.400) | (114.300) | (234.400) | (465.800) |
| Furane | 36.745 | 86.854 | 185.060 | 393.420 |
| | (46.000) | (101.800) | (214.900) | (440.300) |
| Pyrrole | 39.810 | 93.430 | 195.800 | 409.200 |
| | (52.200) | (111.300) | (228.600) | (456.300) |
| Thiophene | 39.286 | 92.061 | 195.870 | 414.480 |
| | (54.400) | (106.500) | (218.900) | (449.000) |
| Imidazole | 36.296 | 85.691 | 182.483 | 387.462 |
| | (49.700) | (99.100) | (202.800) | (411.600) |

[a]Reference values from Table V in Ref. 56 are given in parentheses.

Table S11. Dependence of the average Heme-to-Heme electronic coupling, $\langle H_{DA} \rangle$, on the presence of the electrostatic environment and the inclusion (with/without) of the propionate groups in the QM region

| | | Environment within 40 Å | |
|---|---|---|---|
| Heme Pair | Vacuum | Without Propionates | With Propionates |
| 6'-1 | 5.59 | 7.01 | 5.32 |
| 1-2 | 1.11 | 1.13 | 0.91 |
| 2-3 | 4.98 | 5.91 | 4.83 |
| 3-4 | 2.91 | 2.63 | 1.58 |
| 4-5 | 7.90 | 9.76 | 6.17 |
| 5-6 | 1.02 | 1.05 | 0.55 |
| 6-1'' | 6.63 | 7.76 | 6.75 |

Table S12. Comparison of the root mean-squared and root squared-mean electronic couplings (in meV) computed with Absolutely Localized Molecular Orbital Multi-State Density Functional Theory 2 (ALMO(MSDFT2)) on molecular dynamics (MD)-generated configurations

| Heme Pair[a] | $\sqrt{\langle H_{DA}^2 \rangle}$ | $\sqrt{\langle H_{DA} \rangle^2}$ |
|---:|---:|---:|
| 6'-1 | 6.6204 | 5.5939 |
| 1-2 | 1.3550 | 1.1101 |
| 2-3 | 6.2155 | 4.9805 |
| 3-4 | 3.4731 | 2.9101 |
| 4-5 | 8.9531 | 7.9021 |
| 5-6 | 1.3963 | 1.0229 |
| 6-1'' | 7.5031 | 6.6282 |

[a]A single or double prime mark indicates the preceding or proceeding OmcS subunit, respectively.

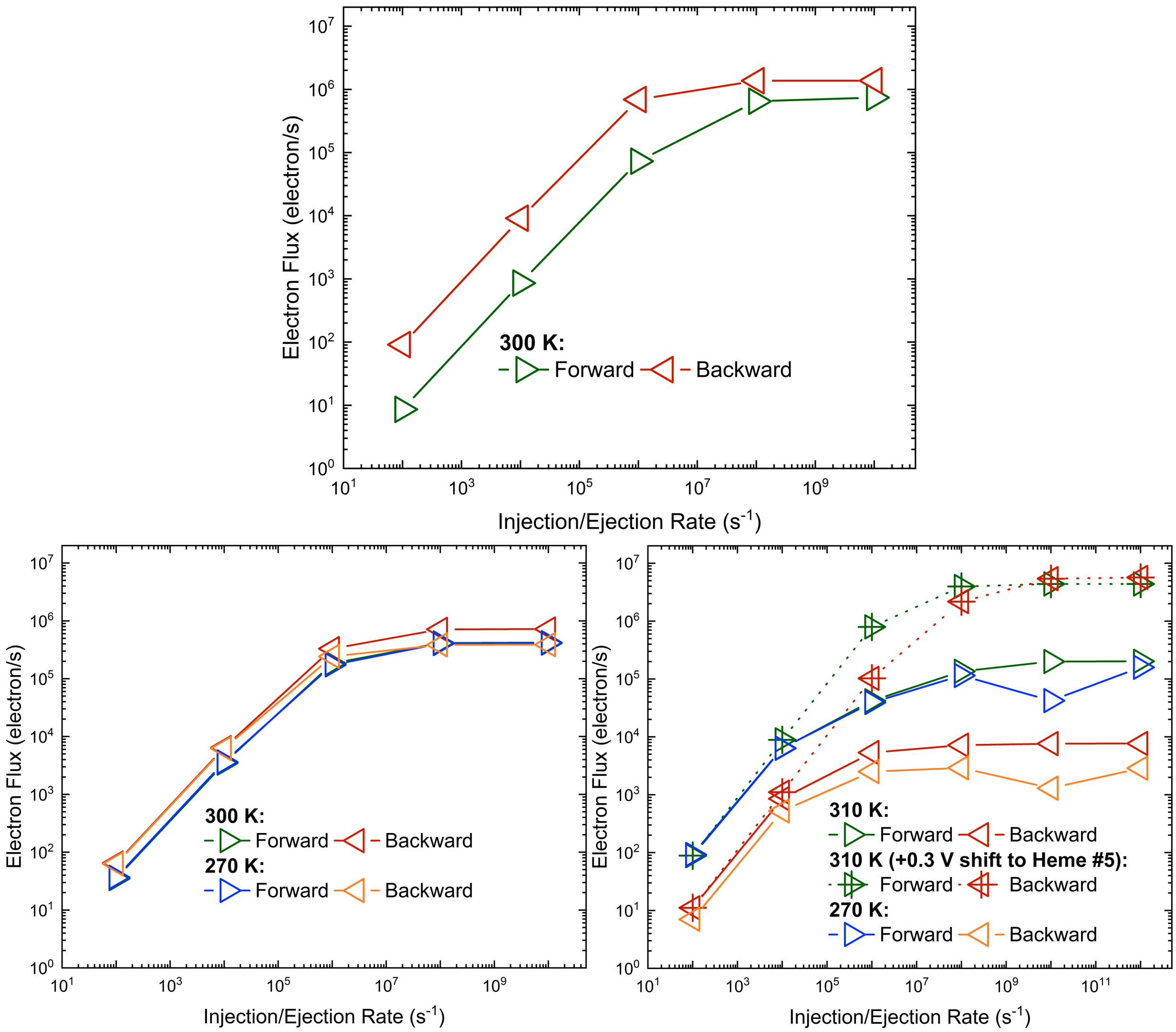

Figure S7. Variation of the charge-hopping flux through a trimeric assembly of OmcS as a function of the electron injection and ejection rates, which were assumed to be identical.

Table S13. Comparison of redox potentials computed at high (310 or 300 K) and low (270 K) temperatures by Dahl *et al.*[1] and in the present work

| Heme # | Dahl *et al.* 310 K | This Work @ 300 K Double-ζ basis | This Work @ 300 K Triple-ζ basis | Dahl *et al.* 270 K | This Work 270 K[a,b] |
|---|---|---|---|---|---|
| 1 | -0.191 ± 0.05 | -0.119 ± 0.027 | -0.063 ± 0.034 | -0.227 ± 0.06 | -0.193 ± 0.088 |
| 2 | -0.108 ± 0.04 | -0.130 ± 0.023 | -0.111 ± 0.027 | -0.200 ± 0.05 | -0.154 ± 0.082 |
| 3 | -0.126 ± 0.05 | -0.108 ± 0.023 | -0.071 ± 0.027 | -0.315 ± 0.04 | -0.152 ± 0.062 |
| 4 | -0.334 ± 0.04 | -0.271 ± 0.027 | -0.248 ± 0.036 | -0.221 ± 0.06 | -0.342 ± 0.080 |
| 5 | -0.521 ± 0.05 | -0.214 ± 0.025 | -0.177 ± 0.024 | -0.477 ± 0.04 | -0.315 ± 0.084 |
| 6 | -0.080 ± 0.06 | -0.128 ± 0.028 | -0.095 ± 0.031 | -0.107 ± 0.04 | -0.218 ± 0.088 |

[a]Computed with the double-ζ basis

[b]Mean ± standard-error-of-the-mean taken over all trajectories

Table S14. $E°$ (V vs. SHE) computed with constant redox or constant redox and pH molecular dynamics under various conditions of coupled/decoupled redox and protonation state changes at high (300 K) and low (270 K) temperatures

| | Case #1[a] | Case #2[b] | | Case #3[c] | | | |
|---|---|---|---|---|---|---|---|
| Heme # | @300 K | @300 K | @270 K | Rep. #1 @300 K | Rep. #2 @300 K | Rep. #3 @300 K | Avg. @300 K[d] |
| 1’ | | -0.189 | -0.209 | -0.186 | -0.176 | -0.181 | -0.181 |
| 2’ | | -0.287 | -0.284 | -0.286 | -0.253 | -0.279 | -0.273 |
| 3’ | | -0.202 | -0.205 | -0.202 | -0.207 | -0.193 | -0.200 |
| 4’ | | -0.296 | -0.332 | -0.284 | -0.291 | -0.275 | -0.283 |
| 5’ | | -0.163 | -0.178 | -0.125 | -0.126 | -0.124 | -0.125 |
| 6’ | | -0.301 | -0.301 | -0.258 | -0.264 | -0.266 | -0.263 |
| 1 | -0.105 | -0.113 | -0.128 | -0.077 | -0.106 | -0.095 | -0.093 |
| 2 | -0.202 | -0.290 | -0.287 | -0.241 | -0.250 | -0.265 | -0.252 |
| 3 | -0.177 | -0.188 | -0.220 | -0.188 | -0.187 | -0.206 | -0.194 |
| 4 | -0.233 | -0.321 | -0.298 | -0.298 | -0.316 | -0.304 | -0.306 |
| 5 | -0.120 | -0.156 | -0.139 | -0.120 | -0.133 | -0.135 | -0.129 |
| 6 | -0.219 | -0.320 | -0.297 | -0.276 | -0.295 | -0.287 | -0.286 |
| 1’’ | | -0.142 | -0.088 | -0.111 | -0.125 | -0.126 | -0.121 |
| 2’’ | | -0.285 | -0.290 | -0.227 | -0.257 | -0.271 | -0.252 |
| 3’’ | | -0.217 | -0.208 | -0.202 | -0.205 | -0.215 | -0.208 |
| 4’’ | | -0.317 | -0.318 | -0.277 | -0.270 | -0.299 | -0.282 |
| 5’’ | | -0.127 | -0.156 | -0.121 | -0.116 | -0.104 | -0.114 |
| 6’’ | | -0.263 | -0.262 | -0.230 | -0.251 | -0.193 | -0.224 |

[a]Case #1 = each of the central six heme and the propionic acid sidechains within the same OmcS subunit were titrated

[b]Case #2 = all hemes were titrated, and the propionic acid groups were locked in the deprotonated state

[c]Case #3 = all hemes and propionic acid groups were titrated

[d]Standard deviations were ≤0.03 V.

| Level of Theory | Nature of Titration | # of Hemes Titrated | Protonation state of Propionic Acid Groups |
|---|---|---|---|
| QM/MM@MD | Single | 6 | Deprotonated |
| CEMD | Single | 6 | Changeable |
| CEMD | Multiple | 18 | Deprotonated |
| C(E,pH)MD | Multiple | 18 | Changeable |

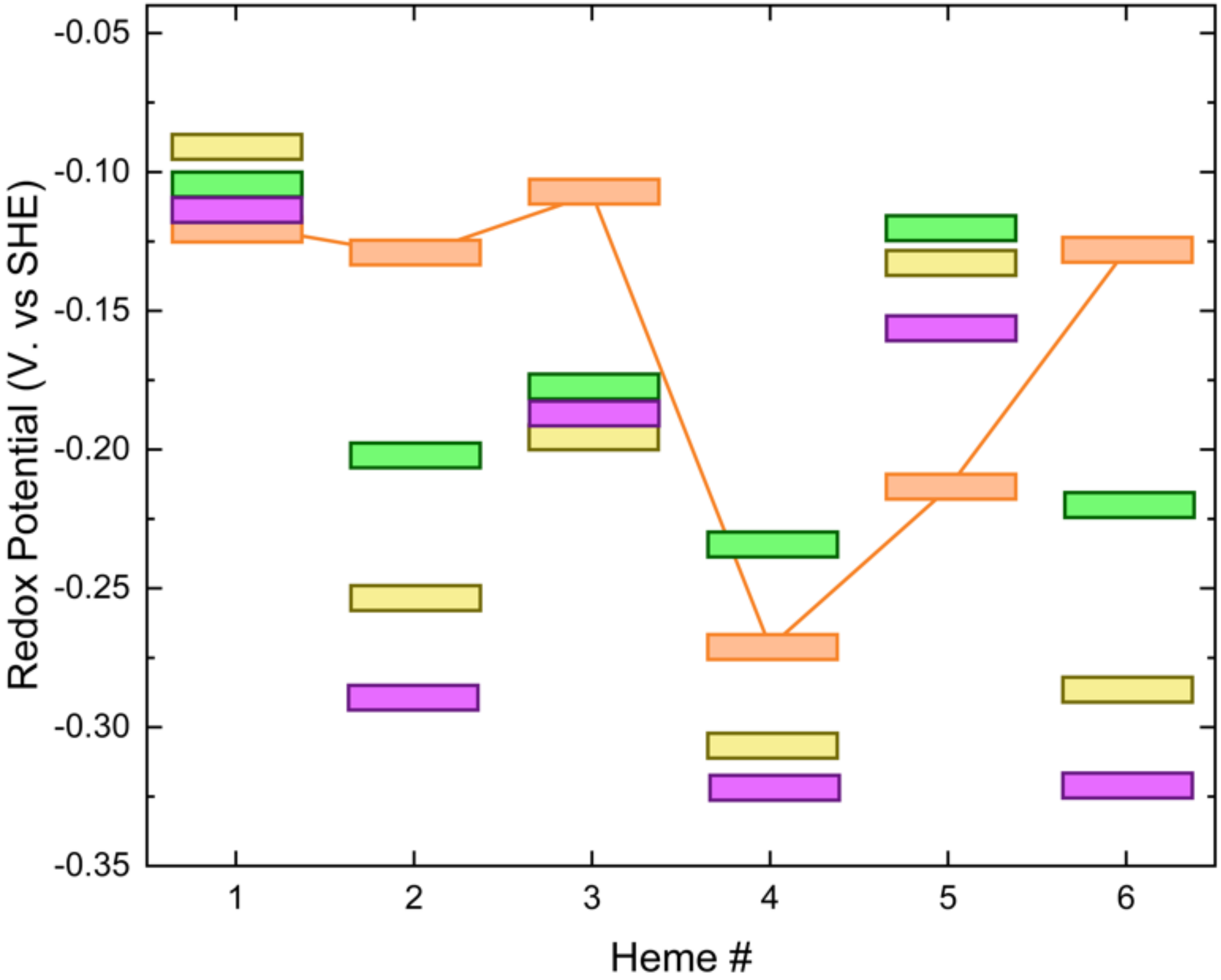

Figure S8. Comparison of $E^{\circ}s$ at 300 K computed with QM/MM@MD and CEMD or C(E,pH)MD methodologies under various conditions to assess the influence of interactions between redox or redox and protonation state changes on the $E^{\circ}$ values. The # and nature of titrated describes how many hemes were titrated and if they were titrated separately or simultaneously.

Table S15. Compariosn of heme redox potentials in various homogenious solvents and the heterogenous OmcS protein filament[a]

| | | Heme # | | | | | |
|---|---|---|---|---|---|---|---|
| Solvent Environment | ϵ(298 K) | #1 | #2 | #3 | #4 | #5 | #6 |
| *n*-Hexane | 1.88 | 0.053 ± 0.024 | 0.047 ± 0.027 | 0.011 ± 0.026 | 0.094 ± 0.030 | 0.033 ± 0.025 | 0.100 ± 0.027 |
| Cyclohexane | 2.02 | 0.029 ± 0.029 | 0.028 ± 0.039 | -0.043 ± 0.037 | 0.055 ± 0.027 | -0.011 ± 0.023 | 0.065 ± 0.025 |
| *o*-Xylene | 2.35 | | -0.085 ± 0.030 | **-0.127 ± 0.024** | -0.028 ± 0.029 | -0.100 ± 0.027 | -0.044 ± 0.025 |
| Dibutylether | 3.05 | **-0.109 ± 0.030** | -0.139 ± 0.030 | -0.183 ± 0.025 | -0.087 ± 0.031 | -0.144 ± 0.027 | -0.079 ± 0.037 |
| Diethylamine | 3.58 | | **-0.169 ± 0.031** | | | **-0.185 ± 0.020** | **-0.120 ± 0.018** |
| Diethylether | 4.24 | -0.198 ± 0.027 | -0.212 ± 0.027 | -0.263 ± 0.023 | -0.152 ± 0.035 | -0.215 ± 0.028 | -0.145 ± 0.030 |
| Diethylsulfide | 5.72 | -0.225 ± 0.036 | -0.257 ± 0.030 | -0.311 ± 0.024 | -0.207 ± 0.033 | -0.262 ± 0.026 | -0.190 ± 0.031 |
| Ethanethiol | 6.67 | | | | **-0.228 ± 0.033** | | |
| 2,6-Dimethylpyridine | 7.17 | -0.254 ± 0.031 | -0.279 ± 0.030 | -0.347 ± 0.023 | -0.234 ± 0.029 | -0.285 ± 0.028 | -0.212 ± 0.031 |
| Tetrahydrofuran | 7.43 | -0.265 ± 0.029 | -0.285 ± 0.030 | -0.342 ± 0.025 | -0.243 ± 0.028 | -0.285 ± 0.028 | -0.218 ± 0.031 |
| OmcS | | -0.109 ±0.067 | -0.184 ±0.055 | -0.126 ±0.060 | -0.225 ±0.061 | -0.185 ±0.058 | -0.121 ±0.052 |

[a]Redox potentials were computed for a common set of 26 conformers for each heme in each solvent environment (see Table S6). The homogeneous solvents were modeled with the polarizable continuum model, whereas snapshots from all-atom but non-polarizable MD with explicit water molecules was used for the protein context.

[b]Redox potentials in a homogeneous solvent for a particular heme that matched the potential of that heme in OmcS are emphasized in **bold** font

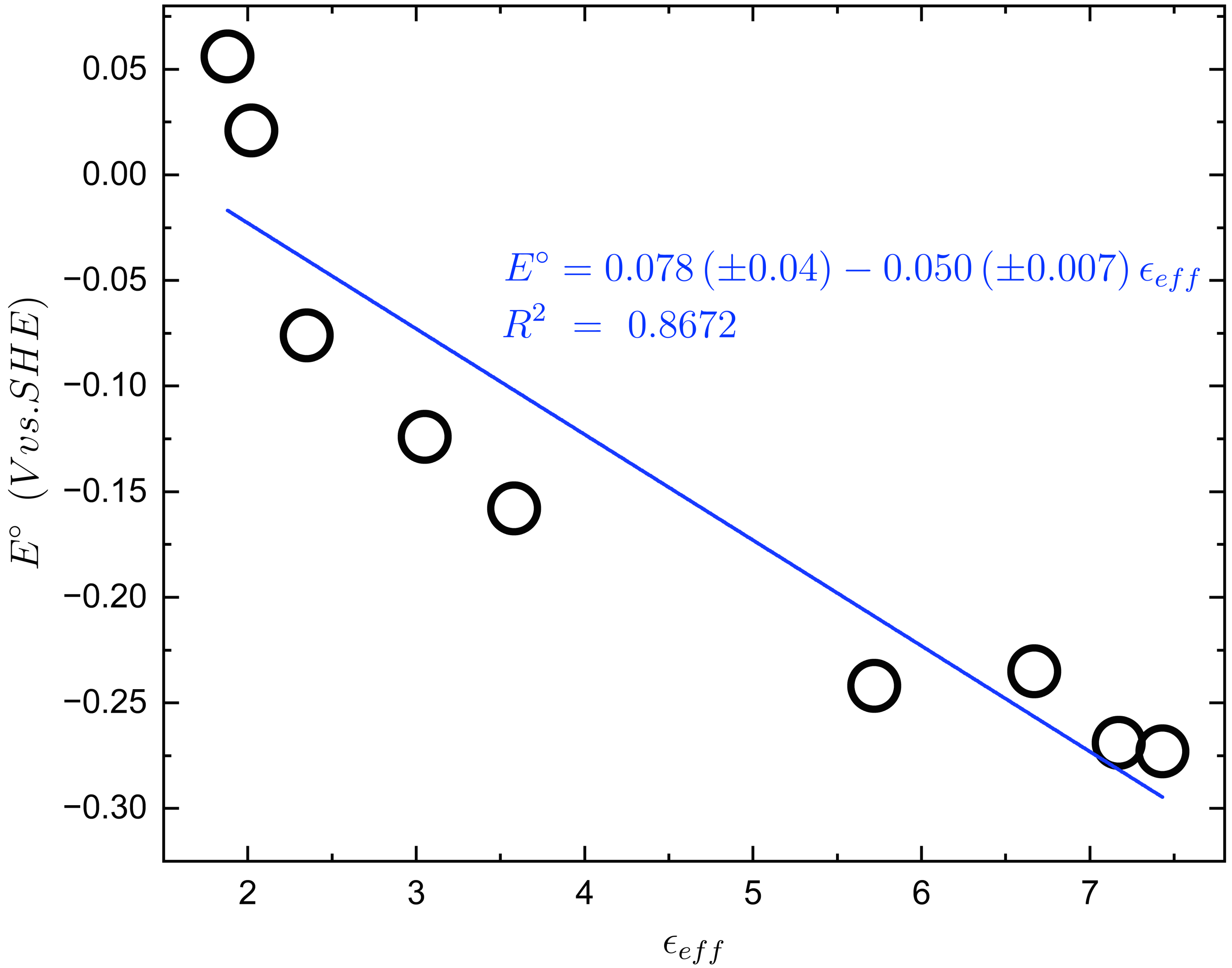

Figure 9. Correlation between heme redox potential and the static (or "effective" when assigned to a protein environment) dielectric constant of the surrounding medium. The redox potentials were computed for a sub-ensemble of representative configurations using B3LYP/double-ζ in conjunction with the polarizable continuum model for a variety of solvents. The redox potential at each effective dielectric is reported as an average of the potentials for the six hemes given in Table S15.

Table S16. Decomposition of redox-linked changes in electrostatic interaction energy for each heme at 300 and 270 K.

| | 300 K | | | | | |
|---|---|---|---|---|---|---|
| Heme # | 1 | 2 | 3 | 4 | 5 | 6 |
| Full Env. | -0.699 ± 0.063 | -0.705 ± 0.067 | -0.529 ± 0.040 | -0.659 ± 0.040 | -0.727 ± 0.037 | -0.593 ± 0.043 |
| Solvent | -0.675 ± 0.059 | -0.658 ± 0.054 | -0.677 ± 0.040 | -0.95 ± 0.042 | -1.119 ± 0.039 | -0.521 ± 0.043 |
| Protein | -0.024 ± 0.065 | -0.047 ± 0.058 | 0.148 ± 0.018 | 0.291 ± 0.028 | 0.392 ± 0.034 | -0.072 ± 0.047 |
| Non-Polar | -0.174 ± 0.037 | -0.338 ± 0.030 | -0.132 ± 0.027 | -0.251 ± 0.032 | -0.256 ± 0.019 | -0.616 ± 0.021 |
| Aromatic | 0.051 ± 0.006 | 0.263 ± 0.023 | 0.084 ± 0.008 | -0.092 ± 0.009 | 0.005 ± 0.006 | 0.025 ± 0.010 |
| Polar | 0.222 ± 0.022 | 0.108 ± 0.029 | 0.177 ± 0.020 | 0.120 ± 0.025 | -0.213 ± 0.030 | 0.064 ± 0.018 |
| Acidic | -1.174 ± 0.034 | -0.547 ± 0.012 | -0.173 ± 0.006 | -0.067 ± 0.009 | -0.106 ± 0.029 | -0.065 ± 0.010 |
| Basic | 1.376 ± 0.045 | 0.273 ± 0.017 | 0.419 ± 0.031 | 0.310 ± 0.018 | 0.528 ± 0.023 | 0.807 ± 0.036 |
| Other Hemes | 0.678 ± 0.013 | 0.432 ± 0.010 | 0.684 ± 0.009 | 0.483 ± 0.009 | 0.825 ± 0.011 | 0.442 ± 0.014 |
| Propionates | -1.653 ± 0.038 | -0.849 ± 0.021 | -1.738 ± 0.035 | -1.452 + 0.026 | -1.901 ± 0.023 | -1.177 ± 0.025 |
| | 270 K | | | | | |
| Heme # | 1 | 2 | 3 | 4 | 5 | 6 |
| Full Env. | -0.802 ± 0.017 | -0.750 ± 0.016 | -0.646 ± 0.012 | -0.796 ± 0.014 | -1.149 ± 0.018 | -0.958 ± 0.014 |
| Solvent | -1.039 ± 0.020 | -0.831 ± 0.016 | -0.741 ± 0.012 | -1.041 ± 0.014 | -1.278 ± 0.018 | -0.429 ± 0.016 |
| Protein | 0.237 ± 0.021 | 0.081 ± 0.018 | 0.095 ± 0.005 | 0.245 ± 0.009 | 0.128 ± 0.015 | -0.529 ± 0.018 |
| Non-Polar | -0.147 ± 0.008 | -0.350 ± 0.007 | -0.145 ± 0.009 | -0.307 ± 0.010 | -0.162 ± 0.009 | -0.503 ± 0.006 |
| Aromatic | 0.045 ± 0.002 | 0.297 ± 0.007 | 0.033 ± 0.003 | -0.089 ± 0.003 | -0.013 ± 0.002 | 0.132 ± 0.004 |
| Polar | 0.262 ± 0.007 | -0.016 ± 0.009 | 0.192 ± 0.006 | 0.058 ± 0.008 | -0.286 ± 0.008 | 0.147 ± 0.008 |
| Acidic | -1.166 ± 0.008 | -0.543 ± 0.005 | -0.182 ± 0.002 | -0.054 ± 0.003 | -0.349 ± 0.018 | -0.293 ± 0.007 |
| Basic | 0.979 ± 0.015 | 0.272 ± 0.007 | 0.393 ± 0.008 | 0.254 ± 0.006 | 0.549 ± 0.006 | 0.702 ± 0.009 |
| Other Hemes | 0.678 ± 0.004 | 0.437 ± 0.003 | 0.687 ± 0.003 | 0.484 ± 0.003 | 0.793 ± 0.003 | 0.508 ± 0.004 |

| | | | | | | |
|---|---|---|---|---|---|---|
| Propionates | -1.69<br>± 0.016 | -0.927<br>± 0.005 | -1.719<br>± 0.011 | -1.387<br>± 0.008 | -1.811<br>± 0.008 | -1.122<br>± 0.008 |

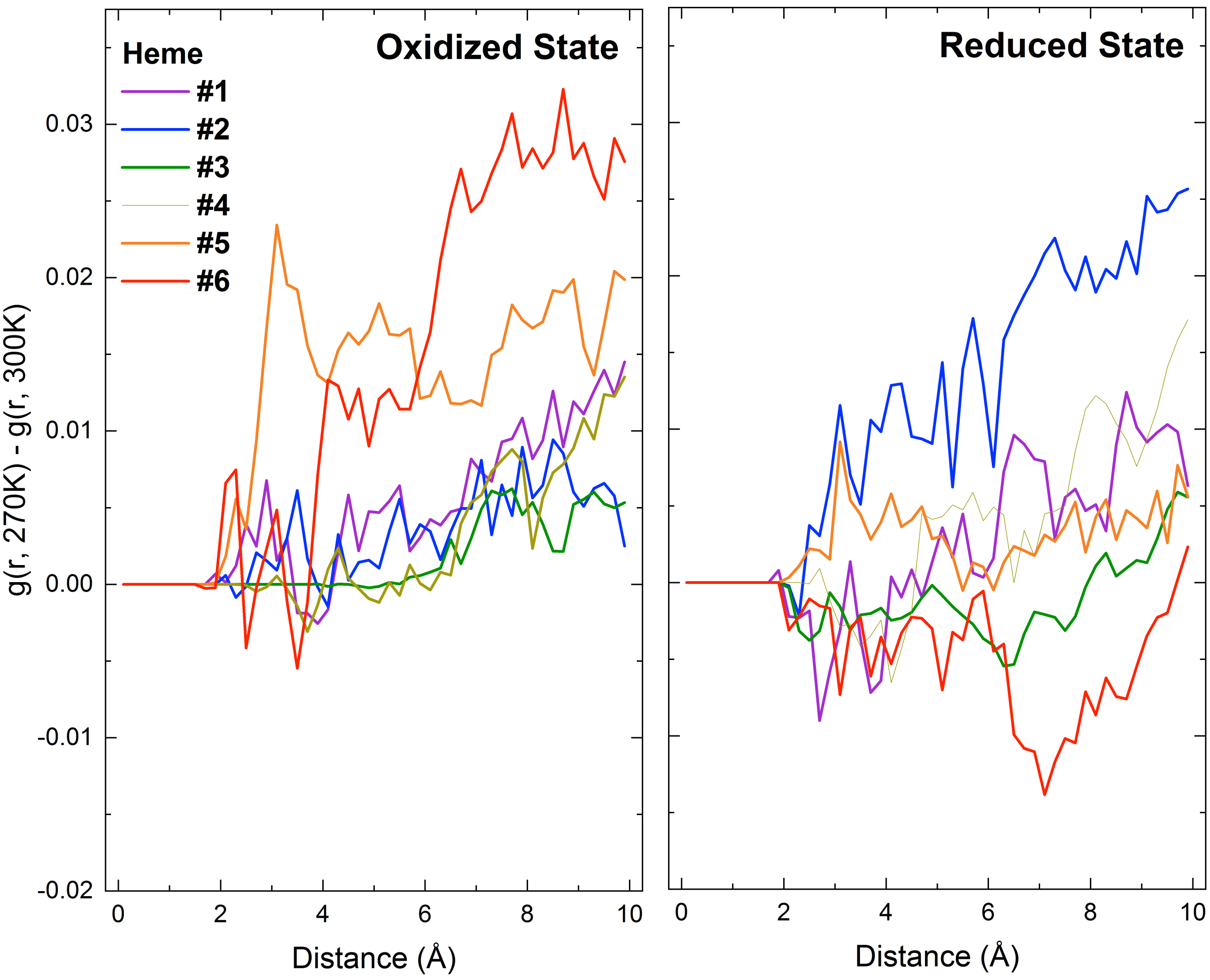

Figure S10. Cooling-induced changes in the normalized radial distribution functions (g(r,T)) for the solvent with respect to each heme in the (*left*) oxidized and (*right*) reduced states. g(r,T) is normalized to the density of pure water (0.033456 molecules/$Å^3$).

Table S17. Characterization of the H-bonding network as a function of temperature

| | VMD | | | | CPPTRAJ | | | | |
|---|---|---|---|---|---|---|---|---|---|
| | Intra-Protein | | | | | | | | Solute-Solvent |
| Temp. (K) | Total | Common to 300 K | Only present at 300 K | Not present at 300 K | Total | Common to 300 K | Only present at 300 K | Not present at 300 K | Total |
| 100 | 370.2 | 157.2 | 106.6 | 103.2 | 370.0 | 156.7 | 106.6 | 103.9 | 1119.3 |
| 125 | 365.9 | 162.3 | 97.2 | 86.9 | 365.4 | 162 | 97.2 | 87.1 | 1128.9 |
| 150 | 353.7 | 144.6 | 96.4 | 88.6 | 353.6 | 144.4 | 96.3 | 89 | 1137.1 |
| 175 | 343.1 | 138.9 | 86.2 | 67.2 | 342.9 | 138.3 | 86.5 | 67.6 | 1188.1 |
| 200 | 336.4 | 131.1 | 71.7 | 64.9 | 336.4 | 130.2 | 71.9 | 65.8 | 1121.8 |
| 225 | 324.3 | 108.7 | 59.6 | 52.1 | 324.3 | 107.9 | 59.7 | 52.9 | 1096.9 |
| 250 | 305.0 | 96.5 | 45 | 42.2 | 304.8 | 96.3 | 45.2 | 42.6 | 1039.9 |
| 270 | 278.0 | 56.6 | 24.9 | 13.4 | 277.9 | 56.9 | 24.9 | 13.4 | 974.4 |
| 300 | 279.3 | 0.0 | 0.0 | 0.0 | 279.2 | 0.0 | 0.0 | 0.0 | 911.9 |
| 325 | 272.6 | 69.6 | 18.7 | 32.8 | 272.5 | 69.1 | 19.3 | 32.8 | 818.9 |
| 350 | 260.1 | 75.3 | 27.2 | 35.2 | 260.0 | 74.4 | 28 | 35.4 | 747.4 |
| 375 | 249.8 | 76.7 | 31.5 | 39.7 | 249.7 | 76.2 | 32 | 39.9 | 663.2 |
| 400 | 242.7 | 85.7 | 29.0 | 38.7 | 242.5 | 85.6 | 29.2 | 38.8 | 595.0 |

Table S18. Analysis of the temperature dependence of the intra-protein H-bonding network in terms of the norms of various occupancy matrices

| Temp. (K) | CHF[a] | ΔCHF[b] | $\Delta CHF_{shared}$[c] | ($\Delta CHF - \Delta CHF_{shared}$)/ ΔCHF |
|---|---|---|---|---|
| 100 | 16.3 | 13.1 | 10.4 | 0.2 |
| 125 | 15.8 | 12.4 | 10.3 | 0.2 |
| 150 | 15.2 | 11.8 | 9.6 | 0.2 |
| 175 | 14.4 | 10.4 | 8.2 | 0.2 |
| 200 | 13.8 | 9.7 | 7.8 | 0.2 |
| 225 | 12.9 | 8.3 | 6.8 | 0.2 |
| 250 | 12.2 | 7.5 | 6.4 | 0.2 |
| 270 | 11.6 | 4.4 | 4.1 | 0.1 |
| 300[d] | 10.8 | 0.0 | 0.0 | |
| 325 | 10.1 | 5.6 | 4.6 | 0.2 |
| 350 | 9.5 | 6.0 | 5.2 | 0.1 |
| 375 | 8.9 | 6.4 | 5.4 | 0.2 |
| 400 | 8.5 | 6.5 | 5.5 | 0.2 |

[a]Characteristic H-bonding Frequency, which is the norm of a matrix of per residue donor-acceptor H-bonding occupancies.
[b]Difference Characteristic H-bonding Frequency, which is the norm of a matrix of differences in per-residue donor-acceptor H-bonding occupancies between two temperatures; 300 K was taken as the reference temperature.
[c]The ΔCHF filtered to only contain non-zero elements at both 300 K and some other temperature
[d]Some entries are left blank because 300 K was taken as the reference for those metrics.

Table S19. H-bonds within 10 Å of the central six hemes in a trimeric assembly of OmcS that had a change in occupancy > 10% at some other temperature relative to the occupancy at 300 K. AMBER99SB residue and atom names, as well as residue IDs particular to the simulation are given.

| Donor | Acceptor | Distance (Å) @ 300 K | Distance (Å) @ Other Temp. | % Difference in Occupancy |
|---|---|---|---|---|
| | | 100 K | | |
| HEH-1271_ND12 | ALA-334_O | 5.9 | 5.7 | 74.5 |
| HIO-335_N | GLY-83_O | 6.2 | 6.4 | 21.2 |
| HEH-1271_ND11 | PRO-129_O | 7.3 | 7.2 | 47.6 |
| TRP-238_NE1 | GLY-190_O | 7.4 | 7.6 | 25.1 |
| HIO-144_N | CYO-140_O | 8.2 | 7.8 | 25.4 |
| ARG-187_NH2 | PRN-1279_O1 | 8.0 | 8.1 | 31.3 |
| GLY-190_N | CYO-143_O | 8.1 | 8.2 | 60.8 |
| GLY-83_N | PRN-1278_O2 | 7.9 | 8.2 | 17.0 |
| ASN-241_N | TRP-238_O | 8.2 | 8.5 | 13.8 |
| TYR-186_OH | PRN-1279_O1 | 8.8 | 8.8 | 11.4 |
| ARG-344_NH2 | PRN-1279_O2 | 10.1 | 10.0 | 12.3 |
| THR-81_OG1 | PRN-1278_O1 | 10.4 | 10.6 | 11.6 |
| | | 125 K | | |
| HEH-1271_ND12 | ALA-334_O | 5.9 | 5.8 | 28.5 |
| HIO-335_N | GLY-83_O | 6.2 | 6.3 | 17.6 |
| HIO-110_N | HIO-335_O | 6.9 | 7.2 | 17.1 |
| HEH-1271_ND11 | PRO-129_O | 7.3 | 7.3 | 46.3 |
| TRP-238_NE1 | GLY-190_O | 7.4 | 7.3 | 22.9 |
| GLY-83_N | PRN-1278_O2 | 7.9 | 8.0 | 14.9 |
| GLY-190_N | CYO-143_O | 8.1 | 8.2 | 33.8 |
| ARG-187_NE | TYR-186_OH | 9.4 | 9.6 | 12.6 |
| TYR-231_OH | PRN-1279_O2 | 10.3 | 10.1 | 39.2 |
| ASP-145_N | ILE-188_O | 11.8 | 11.4 | 30.8 |
| | | 150 K | | |
| HIO-335_N | GLY-83_O | 6.2 | 6.2 | 46.4 |
| HIO-110_N | HIO-335_O | 6.9 | 7.1 | 17.2 |
| HEH-1271_ND11 | PRO-129_O | 7.3 | 7.2 | 42.0 |
| TRP-238_NE1 | GLY-190_O | 7.4 | 7.5 | 13.6 |
| GLY-190_N | CYO-143_O | 8.1 | 8.0 | 18.2 |
| HIO-144_N | CYO-140_O | 8.2 | 8.0 | 17.1 |
| ARG-187_NH2 | PRN-1279_O1 | 8.0 | 8.3 | 18.8 |
| LEU-189_N | TYR-186_O | 8.8 | 9.0 | 15.0 |
| ARG-187_NE | TYR-186_OH | 9.4 | 9.7 | 14.9 |

| | | | | |
|---|---|---|---|---|
| PHE-86_N | ILE-64_O | 9.7 | 9.7 | 58.5 |
| SER-142_N | HIE-139_O | 10.4 | 10.1 | 10.3 |
| THR-81_OG1 | PRN-1278_O1 | 10.4 | 10.4 | 19.3 |
| TYR-231_OH | PRN-1279_O2 | 10.3 | 10.4 | 42.0 |
| | | 175 K | | |
| HIO-335_N | GLY-83_O | 6.2 | 6.4 | 46.9 |
| TRP-238_NE1 | GLY-190_O | 7.4 | 7.1 | 23.7 |
| HEH-1271_ND11 | PRO-129_O | 7.3 | 7.4 | 25.2 |
| HIO-110_N | HIO-335_O | 6.9 | 7.4 | 31.1 |
| GLY-190_N | CYO-143_O | 8.1 | 7.7 | 27.1 |
| GLY-83_N | PRN-1278_O2 | 7.9 | 7.9 | 12.5 |
| ARG-333_N | MET-342_O | 8.9 | 9.2 | 22.2 |
| ARG-333_NH2 | THR-81_O | 9.8 | 9.9 | 17.4 |
| TYR-231_OH | PRN-1279_O2 | 10.3 | 9.9 | 11.5 |
| PHE-86_N | ILE-64_O | 9.7 | 10.0 | 35.0 |
| THR-81_OG1 | PRN-1278_O1 | 10.4 | 10.3 | 10.1 |
| | | 200 K | | |
| HIO-110_N | HIO-335_O | 6.9 | 7.0 | 10.7 |
| CYO-239_N | MET-235_O | 8.0 | 7.4 | 18.9 |
| TRP-238_NE1 | GLY-190_O | 7.4 | 7.4 | 13.6 |
| HEH-1271_ND11 | PRO-129_O | 7.3 | 7.5 | 19.9 |
| TRP-238_N | GLY-234_O | 8.2 | 7.8 | 41.3 |
| ARG-187_NH2 | PRN-1279_O1 | 8.0 | 8.3 | 17.2 |
| TYR-231_OH | PRN-1279_O2 | 10.3 | 10.3 | 39.3 |
| | | 225 K | | |
| HEH-1271_ND12 | ALA-334_O | 5.9 | 5.8 | 10.1 |
| HIO-335_N | GLY-83_O | 6.2 | 6.0 | 10.1 |
| HIO-110_N | HIO-335_O | 6.9 | 6.8 | 19.9 |
| CYO-239_N | MET-235_O | 8.0 | 7.5 | 56.4 |
| TRP-238_NE1 | GLY-190_O | 7.4 | 7.6 | 15.1 |
| GLY-190_N | CYO-143_O | 8.1 | 8.2 | 17.5 |
| HIO-144_N | CYO-140_O | 8.2 | 8.3 | 31.7 |
| ARG-187_NH2 | PRN-1279_O1 | 8.0 | 8.5 | 16.4 |
| THR-81_OG1 | PRN-1278_O1 | 10.4 | 10.3 | 18.8 |
| SER-142_OG | TYR-133_OH | 10.6 | 10.6 | 10.7 |
| TYR-231_OH | PRN-1279_O2 | 10.3 | 10.7 | 37.0 |
| ALA-240_N | SER-236_O | 10.7 | 10.9 | 12.9 |
| CYO-328_N | SER-236_OG | 10.7 | 10.9 | 12.2 |
| VAL-89_N | PHE-86_O | 11.2 | 10.9 | 12.3 |
| | | 250 K | | |

| | | | | |
|---|---|---|---|---|
| HEH-1271_ND11 | PRO-129_O | 7.3 | 7.2 | 16.3 |
| GLY-190_N | CYO-143_O | 8.1 | 7.9 | 10.4 |
| HIO-144_N | CYO-140_O | 8.2 | 8.2 | 16.1 |
| PHE-86_N | ILE-64_O | 9.7 | 9.9 | 10.2 |
| ARG-187_NH1 | PRN-1275_O1 | 10.0 | 10.0 | 14.1 |
| ARG-187_NH1 | PRN-1275_O2 | 10 | 10 | 37.6 |
| TYR-231_OH | PRN-1279_O2 | 10.3 | 10.1 | 38.0 |
| HIE-139_N | SER-142_OG | 10.4 | 10.4 | 20.9 |
| LEU-329_N | ASN-327_OD1 | 10.6 | 10.8 | 23.9 |
| 270 K | | | | |
| HIO-110_N | HIO-335_O | 6.9 | 6.8 | 14.2 |
| CYO-239_N | MET-235_O | 8 | 8 | 21.9 |
| GLY-83_N | PRN-1278_O2 | 7.9 | 8 | 10.6 |
| ARG-333_NH2 | THR-81_O | 9.8 | 9.8 | 14.8 |
| ARG-187_NH1 | PRN-1275_O1 | 10 | 10 | 18.0 |
| HIO-332_N | CYO-328_O | 10 | 10.2 | 12.0 |
| HIE-139_N | SER-142_OG | 10.4 | 10.4 | 14.5 |
| 325 K | | | | |
| HIO-332_N | CYO-328_O | 10 | 10.2 | 21.0 |
| CYO-328_N | SER-236_OG | 10.7 | 10.9 | 10.6 |
| 350 K | | | | |
| CYO-239_N | MET-235_O | 8 | 7.9 | 17.2 |
| ARG-333_NH2 | THR-81_O | 9.8 | 9.6 | 42.8 |
| PHE-86_N | ILE-64_O | 9.7 | 9.9 | 10.2 |
| ARG-187_NH1 | PRN-1275_O1 | 10 | 10.1 | 22.8 |
| ARG-187_NH1 | PRN-1275_O2 | 10 | 10.1 | 12.8 |
| LEU-329_N | ASN-327_OD1 | 10.6 | 10.6 | 20.2 |
| 375 K | | | | |
| HIO-110_N | HIO-335_O | 6.9 | 6.9 | 11.3 |
| TRP-238_NE1 | GLY-190_O | 7.4 | 7.3 | 10.1 |
| CYO-239_N | MET-235_O | 8 | 7.7 | 20.5 |
| TYR-186_OH | PRN-1279_O1 | 8.8 | 8.8 | 52.3 |
| ARG-333_NH2 | THR-81_O | 9.8 | 9.7 | 11.5 |
| HIE-139_N | SER-142_OG | 10.4 | 10.5 | 10.4 |
| 400 K | | | | |
| HIO-335_N | GLY-83_O | 6.2 | 6.2 | 10.5 |
| HEH-1271_ND11 | PRO-129_O | 7.3 | 7.3 | 11.4 |
| TYR-186_OH | PRN-1279_O1 | 8.8 | 9 | 45.0 |
| PHE-86_N | ILE-64_O | 9.7 | 10 | 11.4 |
| HIO-332_N | CYO-328_O | 10 | 10.2 | 15.7 |

| | | | | |
|---|---|---|---|---|
| ARG-187_NH1 | PRN-1275_O2 | 10 | 10.9 | 11.7 |

Table S20. H-bonds within 10 Å of the central six hemes in a trimeric assembly of OmcS that were unique to a 300 K simulation when compared to a simulation at another temperature, and for which the occupancy at the two temperatures differed by > 10%. AMBER99SB residue and atom names, as well as residue IDs particular to the simulation are given.

| Donor | Acceptor | Distance (Å) @ 300 K | Distance (Å) @ Other Temp. | % Difference in Occupancy |
|---|---|---|---|---|
| | | 100 K | | |
| HEH-1277_ND12 | GLY-84_O | 5.0 | 5.0 | 14.9 |
| ARG-333_N | MET-342_O | 8.9 | 8.8 | 12.6 |
| ARG-333_NH2 | THR-81_O | 9.8 | 9.8 | 54.0 |
| ARG-187_NH1 | PRN-1275_O1 | 10.0 | 9.9 | 23.9 |
| ARG-187_NH1 | PRN-1275_O2 | 10.0 | 9.9 | 13.8 |
| HIO-332_N | CYO-328_O | 10.0 | 9.1 | 23.6 |
| HIE-139_N | SER-142_OG | 10.4 | 10.0 | 30.9 |
| LEU-329_N | ASN-327_OD1 | 10.6 | 9.8 | 25.3 |
| ALA-240_N | SER-236_O | 10.7 | 11.4 | 15.1 |
| | | 125 K | | |
| HEH-1277_ND11 | ARG-187_O | 5.1 | 5.1 | 10.5 |
| HIO-144_N | CYO-140_O | 8.2 | 8.0 | 31.2 |
| ARG-333_NH2 | THR-81_O | 9.8 | 9.6 | 54.0 |
| ARG-187_NH1 | PRN-1275_O1 | 10.0 | 9.9 | 23.9 |
| ARG-187_NH1 | PRN-1275_O2 | 10.0 | 9.9 | 13.8 |
| HIO-332_N | CYO-328_O | 10.0 | 9.4 | 23.6 |
| HIE-139_N | SER-142_OG | 10.4 | 10.4 | 30.9 |
| LEU-329_N | ASN-327_OD1 | 10.6 | 10.2 | 25.3 |
| ALA-240_N | SER-236_O | 10.7 | 11.5 | 15.1 |
| | | 150 K | | |
| GLY-83_N | PRN-1278_O2 | 7.9 | 8.1 | 18.8 |
| ARG-333_N | MET-342_O | 8.9 | 8.9 | 12.6 |
| ARG-333_NH2 | THR-81_O | 9.8 | 10.2 | 54.0 |
| ARG-187_NH1 | PRN-1275_O1 | 10.0 | 10.2 | 23.9 |
| ARG-187_NH1 | PRN-1275_O2 | 10.0 | 10.2 | 13.8 |
| HIO-332_N | CYO-328_O | 10.0 | 9.5 | 23.6 |
| HIE-139_N | SER-142_OG | 10.4 | 10.1 | 30.9 |
| LEU-329_N | ASN-327_OD1 | 10.6 | 10.4 | 25.3 |
| ALA-240_N | SER-236_O | 10.7 | 11.5 | 15.1 |
| | | 175 K | | |
| HIO-144_N | CYO-140_O | 8.2 | 8.4 | 31.2 |
| ARG-187_NH1 | PRN-1275_O1 | 10.0 | 9.9 | 23.9 |
| ARG-187_NH1 | PRN-1275_O2 | 10.0 | 9.9 | 13.8 |
| HIO-332_N | CYO-328_O | 10.0 | 9.6 | 23.6 |
| HIE-139_N | SER-142_OG | 10.4 | 10.8 | 30.9 |
| LEU-329_N | ASN-327_OD1 | 10.6 | 10.4 | 25.3 |
| ALA-240_N | SER-236_O | 10.7 | 11.1 | 15.1 |

| | | | | |
|---|---|---|---|---|
| | | 200 K | | |
| ARG-333_N | MET-342_O | 8.9 | 8.5 | 12.6 |
| ARG-333_NH2 | THR-81_O | 9.8 | 8.7 | 54.0 |
| ARG-187_NH1 | PRN-1275_O1 | 10.0 | 10.3 | 23.9 |
| ARG-187_NH1 | PRN-1275_O2 | 10.0 | 10.3 | 13.8 |
| HIO-332_N | CYO-328_O | 10.0 | 9.7 | 23.6 |
| HIE-139_N | SER-142_OG | 10.4 | 10.6 | 30.9 |
| LEU-329_N | ASN-327_OD1 | 10.6 | 10.2 | 25.3 |
| ALA-240_N | SER-236_O | 10.7 | 11.2 | 15.1 |
| | | 225 K | | |
| ARG-333_N | MET-342_O | 8.9 | 8.8 | 12.6 |
| ARG-187_NH1 | PRN-1275_O1 | 10.0 | 10.5 | 23.9 |
| ARG-187_NH1 | PRN-1275_O2 | 10.0 | 10.5 | 13.8 |
| HIO-332_N | CYO-328_O | 10.0 | 9.6 | 23.6 |
| LEU-329_N | ASN-327_OD1 | 10.6 | 10.6 | 25.3 |
| | | 250 K | | |
| ARG-333_N | MET-342_O | 8.9 | 9.1 | 12.6 |
| ARG-333_NH2 | THR-81_O | 9.8 | 9.6 | 54.0 |
| HIO-332_N | CYO-328_O | 10.0 | 10.2 | 23.6 |
| ALA-240_N | SER-236_O | 10.7 | 10.8 | 15.1 |
| TYR-194_OH | SER-141_O | 11.5 | 10.9 | 64.2 |
| | | 270 K | | |
| | | 325 K | | |
| ARG-187_NH1 | PRN-1275_O1 | 10.0 | 10.1 | 23.9 |
| ARG-187_NH1 | PRN-1275_O2 | 10.0 | 10.1 | 13.8 |
| | | 350 K | | |
| | | 375 K | | |
| ARG-187_NH2 | PRN-1279_O1 | 8.0 | 8.6 | 54.3 |
| | | 400 K | | |
| ARG-187_NH2 | PRN-1279_O1 | 8.0 | 8.7 | 54.3 |
| ARG-333_N | MET-342_O | 8.9 | 9.7 | 12.6 |
| ARG-333_NH2 | THR-81_O | 9.8 | 9.7 | 54.0 |

Table S21. H-bonds within 10 Å of the central six hemes in a trimeric assembly of OmcS that were unique to a simulation at a temperature other than 300 K and which had an occupancy > 10%. AMBER99SB residue and atom names, as well as residue IDs particular to the simulation are given.

| Donor | Acceptor | Distance (Å) @ 300 K | Distance (Å) @ Other Temp. | % Difference in Occupancy |
|---|---|---|---|---|
| | | 100 K | | |
| ARG-333_NH1 | PRO-82_O | 9.4 | 9.4 | 58.4 |
| SER-142_OG | HIE-139_O | 10.4 | 10.0 | 88.6 |
| | | 125 K | | |
| HIO-243_N | CYO-239_O | 9.9 | 9.7 | 46.8 |
| SER-142_OG | HIE-139_O | 10.4 | 10.4 | 88.7 |
| | | 150 K | | |
| HIO-243_N | CYO-239_O | 9.9 | 9.6 | 12.9 |
| SER-142_OG | HIE-139_O | 10.4 | 10.1 | 84.9 |
| ARG-344_NH2 | LEU-329_O | 10.6 | 10.2 | 13.4 |
| | | 175 K | | |
| SER-142_OG | HIE-139_O | 10.4 | 10.8 | 92.1 |
| | | 200 K | | |
| ARG-333_NE | ALA-336_O | 9.2 | 8.2 | 17.1 |
| ARG-344_NH2 | LEU-329_O | 10.6 | 10.3 | 15.6 |
| SER-142_OG | HIE-139_O | 10.4 | 10.6 | 89.6 |
| | | 225 K | | |
| MET-235_N | ASN-327_OD1 | 10.2 | 10.1 | 10.4 |
| | | 250K | | |
| ARG-333_NH1 | PRO-82_O | 9.4 | 9.3 | 23.9 |
| SER-142_OG | HIE-139_O | 10.4 | 10.4 | 16.1 |
| | | 270 K | | |
| | | 325 K | | |
| | | 350 K | | |
| | | 375 K | | |
| | | 400 K | | |

Table S22. Heme-to-heme electron trasnfer free energies ($\Delta G_{mn}$) in eV

| | High Temperature[a] | | | Low Temperature[a] | |
|---|---|---|---|---|---|
| Heme Pair | Jiang *et al.* | Dahl *et al.* | This Work | Dahl *et al.* | This Work |
| 1-2 | -0.090 | -0.083 ±0.064 | 0.011 ±0.035 | -0.027 ±0.078 | -0.039 ±0.120 |
| 2-3 | 0.000 | 0.018 ±0.064 | -0.022 ±0.033 | 0.115 ±0.064 | -0.002 ±0.103 |
| 3-4 | 0.090 | 0.208 ±0.064 | 0.163 ±0.035 | -0.094 ±0.072 | 0.190 ±0.101 |
| 4-5 | -0.040 | 0.187 ±0.064 | -0.057 ±0.037 | 0.256 ±0.072 | -0.027 ±0.116 |
| 5-6 | -0.090 | -0.441 ±0.078 | -0.086 ±0.038 | -0.370 ±0.057 | -0.097 ±0.122 |
| 6-1' | 0.120 | -0.080 ±0.078 | -0.128 ±0.039 | 0.120 ±0.072 | -0.025 ±0.124 |

[a]300 K for this work and that by Jiang *et al.*[67]; 310 K for Dahl *et al.*[1]

Table S23. Heme-to-heme electron trasnfer reorganziation energies ($\lambda_{mn}$) in eV

| | High Temperature[a] | | | 270 K | |
|---|---|---|---|---|---|
| Heme Pair | Jiang *et al.*[b] | Dahl *et al.*[c] | This Work[d] | Dahl *et al.* | This Work |
| 1 → 2 | 0.86 | 0.61 | 0.95 | 0.70 | 0.87 |
| 2 → 3 | 0.64 | 0.58 | 0.45 | 0.56 | 0.48 |
| 3 → 4 | 0.71 | 0.66 | 0.60 | 0.55 | 0.57 |
| 4 → 5 | 0.60 | 0.83 | 0.66 | 0.78 | 0.63 |
| 5 → 6 | 0.78 | 0.82 | 0.63 | 0.88 | 0.65 |
| 6 → 1 | 0.66 | 0.68 | 0.68 | 0.79 | 0.56 |

[a]300 K for this work and that by Jiang *et al.*[67]; 310 K for Dahl *et al.*[1]

[b]Estimated from the solvent accessible surface area of the donor and acceptor hemes

[c]Reported as the reaction reorganization energy, which is the square of the Stokes reorganization energy divided by the variance reorganization energy.

[d]Reported as the Stokes reorganization energy. See Table 26 for the variance reorganization energy and how these two quantities compare.

Table S24. Change in the Stokes and variance reorganization energies as a function of the level of hydration[a]

| Heme Pair | $\lambda^{st}$ | | | $\lambda_f^{var}$ | | | $\lambda_b^{var}$ | | | $\chi_g = \frac{(\lambda_f^{var} + \lambda_b^{var})}{\lambda^{st}}$ | | |
|---|---|---|---|---|---|---|---|---|---|---|---|---|
| | Full | First | None | Full | First | None | Full | First | None | Full | First | None |
| 1-2 | 0.95 | 0.80 | 0.42 | 0.87 | 1.64 | 1.72 | 0.95 | 1.59 | 2.39 | 0.96 | 2.01 | 4.94 |
| 2-3 | 0.45 | 0.38 | 0.29 | 0.50 | 1.99 | 1.59 | 0.47 | 1.27 | 0.81 | 1.06 | 4.30 | 4.16 |
| 3-4 | 0.60 | 0.53 | 0.35 | 0.63 | 1.42 | 0.88 | 0.51 | 1.88 | 0.63 | 0.95 | 3.10 | 2.13 |
| 4-5 | 0.66 | 0.67 | 0.36 | 0.64 | 1.10 | 0.62 | 0.50 | 0.81 | 0.54 | 0.86 | 1.43 | 1.62 |
| 5-6 | 0.63 | 0.69 | 0.37 | 0.66 | 1.43 | 0.80 | 0.62 | 1.30 | 1.10 | 1.01 | 1.20 | 2.58 |
| 6-1' | 0.68 | 0.65 | 0.58 | 0.63 | 1.25 | 1.94 | 0.64 | 1.44 | 1.82 | 0.93 | 2.10 | 3.25 |

[a]The hydration state is indicated by ”Full”, “First”, or “None” which stand for a full complement, only the first, or no solvation shells around the protein.

Table S25. Heme-to-heme electron trasnfer couplings $(\langle \mathrm{H_{ji}} \rangle)$ in meV

| | High Temperature[a] | | | 270 K | |
|---|---|---|---|---|---|
| Heme Pair[b] | Jiang *et al.* | Dahl *et al.* | This Work | Dahl *et al.* | This Work |
| 1 → 2 (T) | 1.54 | 8.93 | 1.13 | 9.07 | 1.12 |
| 2 → 3 (S) | 9.35 | 15.9 | 5.91 | 14.9 | 7.14 |
| 3 → 4 (T) | 2.12 | 3.48 | 2.63 | 3.37 | 3.67 |
| 4 → 5 (S) | 6.07 | 12.4 | 9.76 | 12.7 | 8.97 |
| 5 → 6 (T) | 1.18 | 5.68 | 1.05 | 5.56 | 2.71 |
| 6 → 1 (S) | 3.94 | 8.76 | 7.76 | 8.77 | 7.85 |

[a]300 K for this work and that by Jiang *et al.*[67]; 310 K for Dahl *et al.*[1]

[b]Parenthetical T or S indicates, respectively, a T- or slip-stacked heme pair

Table S26. Heme-to-heme electron trasnfer Marcus rates ($k_{nm}$) in s$^{-1}$

| | High Temperature[a] | | | | | |
|---|---|---|---|---|---|---|
| | | $k_{forward}$ | | | $k_{backward}$ | |
| Heme Pair | Jiang *et al.* | Dahl *et al.* | This Work | Jiang *et al.* | Dahl *et al.* | This Work |
| 1 → 2 | 1.8E+06 | 2.5E+10 | 1.9E+06 | 5.1E+07 | 1.1E+09 | 2.9E+06 |
| 2 → 3 | 4.1E+09 | 1.7E+10 | 1.6E+10 | 3.5E+09 | 3.4E+10 | 7.0E+09 |
| 3 → 4 | 4.4E+08 | 5.7E+06 | 1.3E+07 | 1.6E+07 | 1.4E+10 | 7.1E+09 |
| 4 → 5 | 1.1E+09 | 2.4E+07 | 9.6E+09 | 4.8E+09 | 2.7E+10 | 1.1E+09 |
| 5 → 6 | 2.3E+06 | 1.2E+11 | 2.4E+08 | 7.0E+07 | 7.8E+03 | 8.5E+06 |
| 6 → 1” | 4.6E+09 | 2.9E+08 | 2.1E+09 | 4.1E+07 | 1.9E+10 | 1.5E+09 |

| | Low Temperature | | | |
|---|---|---|---|---|
| | $k_{forward}$ | | $k_{backward}$ | |
| Heme Pair | Dahl *et al.* | This Work | Dahl *et al.* | This Work |
| 1 → 2 | 1.6E+09 | 4.7E+06 | 4.8E+08 | 8.8E+05 |
| 2 → 3 | 8.4E+08 | 7.8E+09 | 1.2E+11 | 7.2E+09 |
| 3 → 4 | 4.8E+09 | 5.9E+06 | 8.5E+07 | 2.1E+10 |
| 4 → 5 | 1.2E+06 | 3.6E+09 | 7.1E+10 | 1.1E+09 |
| 5 → 6 | 2.4E+10 | 1.0E+09 | 3.0E+03[b] | 1.6E+07 |
| 6 → 1” | 2.0E+07 | 6.0E+09 | 3.5E+09 | 2.1E+09 |

[a]300 K for this work and that by Jiang *et al.*[67]; 310 K for Dahl *et al.*[1]

[b]This rate is incorrectly reported as 4.51 × $10^3$. All other rates were reproduciable.

Table 27. Diffusion constants and protein-limited charge hopping fluxes for an OmcS filament

| Temperature | Single-Particle Diffusion | Protein-Limited Flux Heme #1 → #6 | Heme #6 → #1 | Ref. for Used Marcus Rates |
|---|---|---|---|---|
| 310 K | $2.7 \times 10^{-11}$ | $2.0 \times 10^{5}$ | $7.6 \times 10^{3}$ | Ref. 1 |
| 310 K | $1.3 \times 10^{-9}$ | $2.2 \times 10^{6}$ | $4.1 \times 10^{5}$ | Ref. 1[a] |
| 310 K | $1.5 \times 10^{-8}$ | $4.4 \times 10^{6}$ | $5.4 \times 10^{6}$ | Ref. 1[b] |
| 300 K | $4.2 \times 10^{-9}$ | $1.4 \times 10^{6}$ | $7.4 \times 10^{5}$ | Ref. 67 |
| 300 K | $1.2 \times 10^{-8}$ | $5.8 \times 10^{5}$ | $7.0 \times 10^{5}$ | This Work |
| 300 K | $9.1 \times 10^{8}$ | $6.3 \times 10^{7}$ | $7.0 \times 10^{7}$ | This Work[c] |
| 300 K | $4.5 \times 10^{6}$ | $2.9 \times 10^{8}$ | $4.4 \times 10^{8}$ | This Work[d] |
| 270 K | $1.1 \times 10^{-10}$ | $1.6 \times 10^{5}$ | $3.0 \times 10^{3}$ | Ref. 1 |
| 270 K | $6.1 \times 10^{-9}$ | $4.1 \times 10^{5}$ | $3.9 \times 10^{5}$ | This Work |

[a]The forward and backward Marcus theory rates for electron transfer between hemes #4 and **#5** and **#5** and **#6** were re-computed after shifting the redox potential of heme #5 by +0.15 V. This shift is the amount by which the reported value exceeded experimental expectations and it is the largest error for any of the other hemes relative to the present work

[b]The forward and backward Marcus theory rates for electron transfer between hemes #4 and **#5** and **#5** and **#6** were re-computed after shifting the redox potential of heme #5 by +0.3 V, as suggested by the calculations reported in the present study. The potentials for all other hemes agreed with the present work within |0.16| V.

[c]Computed with reorganization energies corresponding to a system with only the first solvation shell (3411 water molecules and 12 $Na^+$ ions within 3.4 Å of the trimeric OmcS assembly).

[d]Computed with reorganization energies for a fully dehydrated system (no water/ions). Note that the fluxes are not quite protein-limited values; they increase by a factor of ~1.3 if the injection/ejection rate is increased from $10^{10}$ to $10^{12}$ or $10^{14}$ electrons/s. For consistency with the other fluxes, we report the values obtained with the same injection/ejection rate ($10^{10}$ electrons/s).

Table S28. Predicted charge hopping currents based on the experimentally measured conductivity, the computed diffusion coefficient, and the simulated protein-limited stead-state charge flux

| | V = 1.1 V<br>L = 1.4 × $10^{-6}$ cm | | V = 0.1 V<br>L = 3.0 × $10^{-5}$ cm |
|---|---|---|---|
| Experiment | 6.17 × $10^{3}$ | | 2.67 × $10^{1}$ |
| | Diffusion Model | Flux Model | Diffusion Model |
| Jiang et al (300 K) | 1.3 × $10^{-1}$ | 1.7 × $10^{-1}$ | 5.6 × $10^{-4}$ |
| Dahl et al. (310 K) | 8.4 × $10^{-4}$ | 1.7 × $10^{-2}$ | 3.7 × $10^{-7}$ |
| Dahl et al. (310 K)* | 4.0 × $10^{-2}$ | 2.1 × $10^{-1}$ | 1.7 × $10^{-4}$ |
| Dahl et al. (310 K)[b] | 4.6 × $10^{-1}$ | 7.8 × $10^{-1}$ | 2.0 × $10^{-3}$ |
| Dahl et al. (270 K) | 3.4 × $10^{-3}$ | 1.3 × $10^{-2}$ | 1.5 × $10^{-5}$ |
| This Work (300 K) | 3.5 × $10^{-1}$ | 1.0 × $10^{-1}$ | 1.5 × $10^{-3}$ |
| This Work (300 K)[c] | 2.8 × $10^{0}$ | 1.6 × $10^{1}$ | 1.2 × $10^{-2}$ |
| This Work (300 K)[d] | 1.4 × $10^{2}$ | 8.2 × $10^{1}$ | 6.0 × $10^{-1}$ |
| This Work (270 K) | 1.9 × $10^{-1}$ | 6.4 × $10^{-2}$ | 4.1 × $10^{-4}$ |

[a]The forward and backward Marcus theory rates for electron transfer between hemes #4 and **#5** and **#5** and **#6** were re-computed after shifting the redox potential of heme #5 by +0.15 V. This shift is the amount by which the reported value exceeded experimental expectations and it is the largest error for any of the other hemes relative to the present work

[b]The forward and backward Marcus theory rates for electron transfer between hemes #4 and **#5** and **#5** and **#6** were re-computed after shifting the redox potential of heme #5 by +0.3 V, as suggested by the calculations reported in the present study. The potentials for all other hemes agreed with the present work within |0.16| V.

[c]Computed with reorganization energies corresponding to a system with only the first solvation shell (3411 water molecules and 12 $Na^+$ ions within 3.4 Å of the trimeric OmcS assembly).

[d]Computed with reorganization energies for a fully dehydrated system (no water/ions). Note that the fluxes are not quite protein-limited values; they increase by a factor of ~1.3 if the injection/ejection rate is increased from $10^{10}$ to $10^{12}$ or $10^{14}$ electrons/s. For consistency with the other fluxes, we report the values obtained with the same injection/ejection rate ($10^{10}$ electrons/s).

Table S29. Computed p$K_a$s for the titratable residues in the central subunit of the simulated OmcS filament before and after the six-electron reduction of all the hemes in that subunit. AMBER residue names are used as well as the residue IDs from the simulaitons. Note that AS4-85, TYR-150, and TYR-194 were excluded from the analysis because of large uncertainties in the computed p$K_a$s for these residues.

| | Oxidized State | | Reduced State | |
|---|---|---|---|---|
| Residue | Hill Coefficient | p$K_a$ | Hill Coefficient | p$K_a$ |
| GL4-8 | 1.5 | 6.62 ± 0.01 | 1.6 | 6.95 ± 0.00 |
| GL4-10 | 0.2 | 8.90 ± 0.080 | 1.6 | 10.36 ± 0.02 |
| HIP-53 | 0.5 | 6.39 ± 0.02 | 0.9 | 6.45 ± 0.08 |
| AS4-56 | 0.5 | 5.03 ± 0.02 | 0.9 | 4.81 ± 0.01 |
| TYR-62 | 1.0 | 12.81 ± 0.01 | 0.7 | 13.40 ± 0.01 |
| HIP-63 | 0.3 | 5.39 ± 0.28 | 1.2 | 7.84 ± 0.00 |
| GL4-68 | 0.8 | 3.59 ± 0.01 | 1.0 | 3.06 ± 0.05 |
| AS4-70 | 0.6 | 5.23 ± 0.01 | 0.7 | 4.70 ± 0.01 |
| LYS-90 | 0.8 | 11.02 ± 0.01 | 0.6 | 10.77 ± 0.02 |
| LYS-91 | 1.2 | 10.51 ± 0.01 | 1.1 | 10.60 ± 0.00 |
| TYR-93 | 1.3 | 15.75 ± 0.02 | 0.4 | 15.97 ± 0.18 |
| GL4-104 | 0.4 | 4.24 ± 0.05 | 0.7 | 4.53 ± 0.08 |
| GL4-106 | 0.3 | 5.93 ± 0.30 | 0.6 | 7.59 ± 0.06 |
| LYS-108 | 0.9 | 7.58 ± 0.42 | 0.2 | 6.24 ± 0.30 |
| AS4-116 | 1.7 | 4.32 ± 0.04 | 0.9 | 4.49 ± 0.05 |
| TYR-117 | 0.9 | 14.78 ± 0.01 | 0.9 | 14.34 ± 0.04 |
| TYR-119 | 0.5 | 13.84 ± 0.19 | 0.7 | 13.64 ± 0.05 |
| AS4-122 | 0.3 | 0.44 ± 0.13 | 1.6 | -3.11 ± 0.38 |
| TYR-133 | 1.2 | 16.55 ± 0.00 | 0.8 | 17.13 ± 0.01 |
| HIP-139 | 0.9 | 7.08 ± 0.01 | 1.1 | 6.46 ± 0.02 |
| AS4-145 | 0.8 | 2.45 ± 0.27 | 1.8 | 3.52 ± 0.02 |
| LYS-149 | 0.4 | 7.33 ± 0.07 | 0.8 | 8.44 ± 0.02 |
| AS4-155 | 1.0 | 4.05 ± 0.00 | 1.0 | 4.16 ± 0.00 |
| LYS-166 | 0.8 | 8.87 ± 0.01 | 1.1 | 8.58 ± 0.06 |
| TYR-171 | 1.3 | 12.02 ± 0.00 | 1.6 | 13.06 ± 0.00 |
| AS4-176 | 0.8 | 3.29 ± 0.00 | 0.8 | 2.70 ± 0.04 |
| TYR-186 | 1.8 | 15.60 ± 0.33 | 1.6 | 15.68 ± 0.16 |
| LYS-197 | 0.7 | 9.60 ± 0.02 | 0.6 | 9.72 ± 0.03 |
| TYR-203 | 1.5 | 13.11 ± 0.00 | 1.4 | 13.11 ± 0.00 |
| TYR-218 | 0.8 | 10.78 ± 0.00 | 1.1 | 12.73 ± 0.02 |
| GL4-222 | 0.5 | 1.26 ± 0.05 | 1.2 | 2.43 ± 0.14 |
| TYR-231 | 1.2 | 15.23 ± 0.06 | 0.7 | 14.88 ± 0.01 |

| | | | | |
|---|---|---|---|---|
| GL4-237 | 0.6 | 4.36 ± 0.12 | 0.7 | 4.52 ± 0.01 |
| AS4-245 | 1.2 | 4.78 ± 0.06 | 0.7 | 5.37 ± 0.01 |
| HIP-247 | 1.1 | 2.63 ± 0.02 | 0.4 | 3.43 ± 0.02 |
| TYR-251 | 2.1 | 13.15 ± 0.03 | 1.5 | 13.58 ± 0.16 |
| LYS-264 | 0.9 | 9.58 ± 0.01 | 0.6 | 9.86 ± 0.03 |
| TYR-273 | 1.2 | 18.30 ± 0.07 | 1.6 | 16.08 ± 0.00 |
| TYR-276 | 0.7 | 11.52 ± 0.02 | 0.7 | 11.93 ± 0.02 |
| LYS-277 | 0.6 | 8.81 ± 0.00 | 0.6 | 9.15 ± 0.01 |
| LYS-278 | 0.8 | 9.34 ± 0.00 | 0.9 | 9.20 ± 0.01 |
| AS4-281 | 0.7 | 2.41 ± 0.02 | 0.8 | 4.32 ± 0.07 |
| TYR-290 | 0.2 | 14.48 ± 1.01 | 1.7 | 13.46 ± 0.24 |
| GL4-297 | 1.1 | 1.39 ± 0.10 | 1.1 | 2.46 ± 0.13 |
| GL4-298 | 0.7 | 2.46 ± 0.03 | 0.2 | 4.50 ± 0.73 |
| AS4-302 | 0.9 | 3.03 ± 0.00 | 0.8 | 2.89 ± 0.00 |
| TYR-303 | 1.8 | 13.21 ± 0.01 | 1.4 | 13.42 ± 0.00 |
| LYS-307 | 0.9 | 10.24 ± 0.01 | 0.7 | 10.62 ± 0.03 |
| HIP-309 | 0.8 | 5.13 ± 0.01 | 0.7 | 5.29 ± 0.01 |
| LYS-311 | 1.0 | 9.69 ± 0.01 | 0.6 | 9.71 ± 0.01 |
| AS4-313 | 1.0 | 1.84 ± 0.01 | 0.9 | 1.96 ± 0.00 |
| AS4-314 | 0.7 | 3.64 ± 0.01 | 0.8 | 3.78 ± 0.00 |
| AS4-321 | 0.8 | 2.50 ± 0.03 | 0.6 | 2.05 ± 0.00 |
| AS4-340 | 0.6 | 2.27 ± 0.07 | 0.2 | 3.82 ± 0.41 |
| TYR-349 | 1.4 | 11.91 ± 0.00 | 1.1 | 11.76 ± 0.02 |
| GL4-350 | 1.0 | 0.34 ± 0.05 | 0.2 | 2.45 ± 0.69 |
| AS4-356 | 1.1 | 3.04 ± 0.00 | 1.0 | 3.01 ± 0.00 |
| TYR-363 | 2.8 | 15.62 ± 1.10 | 2.3 | 15.18 ± 0.01 |
| AS4-366 | 0.8 | 4.64 ± 0.00 | 0.9 | 4.64 ± 0.00 |
| GL4-379 | 1.1 | 1.71 ± 0.04 | 0.9 | 1.01 ± 0.01 |
| TYR-384 | 1.5 | 15.44 ± 0.00 | 1.6 | 15.93 ± 0.00 |
| TYR-385 | 0.2 | 12.94 ± 0.50 | 0.2 | 10.20 ± 0.59 |
| AS4-390 | 0.6 | 2.53 ± 0.03 | 0.5 | 2.26 ± 0.03 |
| LYS-391 | 0.9 | 10.55 ± 0.01 | 0.8 | 10.47 ± 0.05 |
| TYR-395 | 1.1 | 14.09 ± 0.00 | 1.0 | 14.09 ± 0.00 |
| LYS-402 | 0.6 | 13.35 ± 0.02 | 0.6 | 12.72 ± 0.04 |
| LYS-406 | 0.6 | 7.81 ± 0.01 | 0.5 | 6.73 ± 0.04 |
| PRN-1269 | 1.0 | 4.22 ± 0.02 | 0.6 | 3.23 ± 0.02 |
| PRN-1270 | 0.2 | 7.36 ± 0.63 | 0.2 | 11.20 ± 0.77 |
| PRN-1272 | 0.3 | 4.12 ± 0.66 | 1.2 | 7.37 ± 0.13 |
| PRN-1273 | 1.3 | 4.51 ± 0.49 | 0.4 | 3.83 ± 0.08 |
| PRN-1275 | 2.2 | 4.66 ± 1.92 | 0.2 | 2.69 ± 0.69 |

| | | | | |
|---|---|---|---|---|
| PRN-1276 | 1.1 | 2.46 ± 0.03 | 0.9 | 2.67 ± 0.07 |
| PRN-1278 | 0.8 | 0.88 ± 0.01 | 0.7 | 0.92 ± 0.11 |
| PRN-1279 | 0.5 | 1.97 ± 0.11 | 0.5 | 3.88 ± 0.12 |
| PRN-1281 | 1.2 | 4.83 ± 0.05 | 0.6 | 3.68 ± 0.05 |
| PRN-1282 | 0.4 | 2.21 ± 0.21 | 0.6 | 5.05 ± 0.02 |
| PRN-1284 | 1.0 | 6.76 ± 0.02 | 1.0 | 6.94 ± 0.07 |
| PRN-1285 | 0.3 | 6.84 ± 0.23 | 1.1 | 8.62 ± 0.33 |

Table S30. Analysis of the extent of charge compoensation in OmcS as a function of pH for a six-electorn reduction (one electron for each of the central six heems in the simulated filament)

| pH | Total Charge, Oxidized State | Total Change, Reduced State | Net Charge | % of Expected Charge Change |
|---|---|---|---|---|
| 2.0 | 27.1 | 23.3 | -3.8 | 64.0 |
| 2.1 | 26.4 | 22.7 | -3.7 | 61.6 |
| 2.2 | 25.7 | 22.1 | -3.6 | 59.5 |
| 2.3 | 25.0 | 21.5 | -3.5 | 57.6 |
| 2.4 | 24.3 | 20.9 | -3.4 | 56.1 |
| 2.5 | 23.5 | 20.2 | -3.3 | 54.8 |
| 2.6 | 22.7 | 19.5 | -3.2 | 53.8 |
| 2.7 | 22.0 | 18.8 | -3.2 | 53.0 |
| 2.8 | 21.2 | 18.1 | -3.1 | 52.4 |
| 2.9 | 20.5 | 17.3 | -3.1 | 52.1 |
| 3.0 | 19.7 | 16.6 | -3.1 | 52.1 |
| 3.1 | 19.0 | 15.8 | -3.2 | 52.5 |
| 3.2 | 18.3 | 15.1 | -3.2 | 53.2 |
| 3.3 | 17.5 | 14.3 | -3.3 | 54.4 |
| 3.4 | 16.8 | 13.5 | -3.4 | 55.8 |
| 3.5 | 16.1 | 12.7 | -3.5 | 57.5 |
| 3.6 | 15.4 | 11.9 | -3.6 | 59.2 |
| 3.7 | 14.7 | 11.1 | -3.7 | 60.8 |
| 3.8 | 14.0 | 10.3 | -3.7 | 62.0 |
| 3.9 | 13.3 | 9.5 | -3.8 | 62.8 |
| 4.0 | 12.5 | 8.8 | -3.8 | 63.0 |
| 4.1 | 11.8 | 8.0 | -3.8 | 62.5 |
| 4.2 | 11.0 | 7.3 | -3.7 | 61.4 |
| 4.3 | 10.1 | 6.5 | -3.6 | 59.7 |
| 4.4 | 9.2 | 5.8 | -3.4 | 57.4 |
| 4.5 | 8.4 | 5.1 | -3.3 | 54.6 |
| 4.6 | 7.5 | 4.4 | -3.1 | 51.5 |
| 4.7 | 6.6 | 3.7 | -2.9 | 48.5 |
| 4.8 | 5.7 | 3.0 | -2.8 | 45.8 |
| 4.9 | 5.0 | 2.3 | -2.6 | 43.7 |
| 5.0 | 4.2 | 1.7 | -2.5 | 42.2 |
| 5.1 | 3.6 | 1.1 | -2.5 | 41.3 |
| 5.2 | 3.0 | 0.5 | -2.5 | 40.9 |
| 5.3 | 2.4 | 0.0 | -2.5 | 41.0 |
| 5.4 | 1.9 | -0.5 | -2.5 | 41.4 |

| | | | | |
|---|---|---|---|---|
| 5.5 | 1.5 | -1.0 | -2.5 | 41.9 |
| 5.6 | 1.0 | -1.5 | -2.6 | 42.6 |
| 5.7 | 0.6 | -2.0 | -2.6 | 43.4 |
| 5.8 | 0.2 | -2.4 | -2.7 | 44.2 |
| 5.9 | -0.2 | -2.9 | -2.7 | 45.0 |
| 6.0 | -0.6 | -3.3 | -2.8 | 45.8 |
| 6.1 | -0.9 | -3.7 | -2.8 | 46.6 |
| 6.2 | -1.3 | -4.2 | -2.8 | 47.2 |
| 6.3 | -1.7 | -4.6 | -2.9 | 47.8 |
| 6.4 | -2.2 | -5.1 | -2.9 | 48.4 |
| 6.5 | -2.6 | -5.5 | -2.9 | 48.9 |
| 6.6 | -3.0 | -6.0 | -3.0 | 49.5 |
| 6.7 | -3.5 | -6.5 | -3.0 | 50.2 |
| 6.8 | -3.9 | -7.0 | -3.1 | 51.1 |
| 6.9 | -4.4 | -7.5 | -3.1 | 52.3 |
| 7.0 | -4.8 | -8.0 | -3.2 | 53.6 |
| 7.1 | -5.2 | -8.5 | -3.3 | 55.0 |
| 7.2 | -5.6 | -9.0 | -3.4 | 56.4 |
| 7.3 | -6.0 | -9.5 | -3.5 | 57.8 |
| 7.4 | -6.4 | -9.9 | -3.6 | 59.2 |
| 7.5 | -6.8 | -10.4 | -3.6 | 60.5 |
| 7.6 | -7.1 | -10.8 | -3.7 | 61.8 |
| 7.7 | -7.5 | -11.3 | -3.8 | 63.2 |
| 7.8 | -7.8 | -11.7 | -3.9 | 64.5 |
| 7.9 | -8.2 | -12.1 | -4.0 | 66.0 |
| 8.0 | -8.5 | -12.5 | -4.1 | 67.5 |
| 8.1 | -8.8 | -13.0 | -4.2 | 69.2 |
| 8.2 | -9.1 | -13.4 | -4.3 | 70.9 |
| 8.3 | -9.5 | -13.8 | -4.4 | 72.8 |
| 8.4 | -9.8 | -14.3 | -4.5 | 74.7 |
| 8.5 | -10.1 | -14.7 | -4.6 | 76.6 |
| 8.6 | -10.5 | -15.2 | -4.7 | 78.5 |
| 8.7 | -10.8 | -15.6 | -4.8 | 80.2 |
| 8.8 | -11.2 | -16.1 | -4.9 | 81.8 |
| 8.9 | -11.5 | -16.5 | -5.0 | 83.1 |
| 9.0 | -11.9 | -16.9 | -5.0 | 84.1 |
| 9.1 | -12.3 | -17.4 | -5.1 | 84.7 |
| 9.2 | -12.7 | -17.8 | -5.1 | 85.0 |
| 9.3 | -13.1 | -18.2 | -5.1 | 85.0 |
| 9.4 | -13.5 | -18.6 | -5.1 | 84.7 |

| | | | | |
|---|---|---|---|---|
| 9.5 | -13.9 | -19.0 | -5.1 | 84.2 |
| 9.6 | -14.3 | -19.4 | -5.0 | 83.6 |
| 9.7 | -14.8 | -19.8 | -5.0 | 82.8 |
| 9.8 | -15.2 | -20.2 | -4.9 | 82.1 |
| 9.9 | -15.7 | -20.6 | -4.9 | 81.4 |
| 10.0 | -16.1 | -21.0 | -4.9 | 80.9 |
| 10.1 | -16.6 | -21.4 | -4.8 | 80.6 |
| 10.2 | -17.1 | -21.9 | -4.8 | 80.5 |
| 10.3 | -17.5 | -22.4 | -4.8 | 80.5 |
| 10.4 | -18.0 | -22.8 | -4.8 | 80.7 |
| 10.5 | -18.4 | -23.3 | -4.9 | 80.9 |
| 10.6 | -18.9 | -23.7 | -4.9 | 81.1 |
| 10.7 | -19.3 | -24.2 | -4.9 | 81.2 |
| 10.8 | -19.7 | -24.6 | -4.9 | 81.1 |
| 10.9 | -20.1 | -25.0 | -4.9 | 81.0 |
| 11.0 | -20.5 | -25.4 | -4.8 | 80.7 |
| 11.1 | -20.9 | -25.7 | -4.8 | 80.4 |
| 11.2 | -21.3 | -26.1 | -4.8 | 80.1 |
| 11.3 | -21.6 | -26.4 | -4.8 | 79.7 |
| 11.4 | -22.0 | -26.7 | -4.8 | 79.2 |
| 11.5 | -22.3 | -27.0 | -4.7 | 78.7 |
| 11.6 | -22.7 | -27.3 | -4.7 | 78.0 |
| 11.7 | -23.0 | -27.6 | -4.6 | 77.2 |
| 11.8 | -23.4 | -27.9 | -4.6 | 76.3 |
| 11.9 | -23.7 | -28.3 | -4.5 | 75.4 |
| 12.0 | -24.1 | -28.6 | -4.5 | 74.5 |
| 12.1 | -24.4 | -28.9 | -4.4 | 73.7 |
| 12.2 | -24.8 | -29.2 | -4.4 | 73.2 |
| 12.3 | -25.1 | -29.5 | -4.4 | 73.0 |
| 12.4 | -25.5 | -29.8 | -4.4 | 73.1 |
| 12.5 | -25.8 | -30.2 | -4.4 | 73.6 |
| 12.6 | -26.1 | -30.6 | -4.5 | 74.4 |
| 12.7 | -26.5 | -31.0 | -4.5 | 75.5 |
| 12.8 | -26.9 | -31.5 | -4.6 | 76.7 |
| 12.9 | -27.3 | -32.0 | -4.7 | 78.1 |

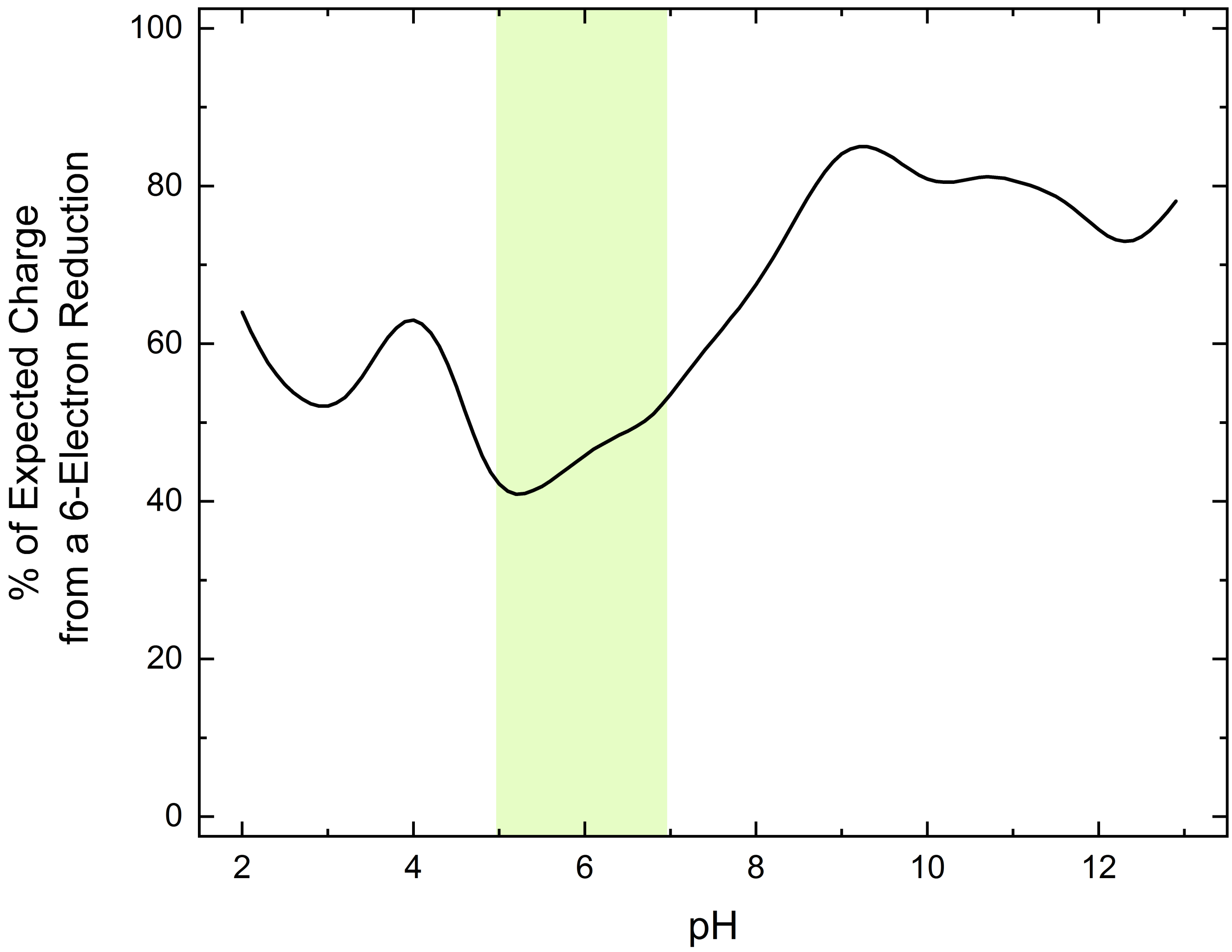

Figure S11. Percentage of the expected change in net charge upon a 6-electron reduction of the six hemes in the central subunit of a OmcS oligomer. The smaller the percentage, the greater the change in net charge is offset up the change in protonation states of pH-sensitive residues in the central subunit of the filament. Note that the greatest extent of this charge compensaiton occurs in the physiologically relevant pH range highlighted in green.

### References

1. Dahl, P. J.; Yi, S. M.; Gu, Y.; Acharya, A.; Shipps, C.; Neu, J.; O'Brien, J. P.; Morzan, U. N.; Chaudhuri, S.; Guberman-Pfeffer, M. J., A 300-fold conductivity increase in microbial cytochrome nanowires due to temperature-induced restructuring of hydrogen bonding networks. *Science Advances* **2022,** *8* (19), eabm7193.
2. Wang, F.; Gu, Y.; O'Brien, J. P.; Yi, S. M.; Yalcin, S. E.; Srikanth, V.; Shen, C.; Vu, D.; Ing, N. L.; Hochbaum, A. I.; Egelman, E. H.; Malvankar, N. S., Structure of Microbial Nanowires Reveals Stacked Hemes that Transport Electrons over Micrometers. *Cell* **2019,** *177* (2), 361-369.e10.
3. Case, D. A.; Cheatham III, T. E.; Darden, T.; Gohlke, H.; Luo, R.; Merz Jr, K. M.; Onufriev, A.; Simmerling, C.; Wang, B.; Woods, R. J., The Amber biomolecular simulation programs. *Journal of computational chemistry* **2005,** *26* (16), 1668-1688.
4. D.A. Case, H. M. A., K. Belfon, I.Y. Ben-Shalom, S.R. Brozell, D.S. Cerutti, T.E. Cheatham, III, V.W.D. Cruzeiro, T.A. Darden, R.E. Duke, G. Giambasu, M.K. Gilson, H. Gohlke, A.W. Goetz, R. Harris, S. Izadi, S.A. Izmailov, C. Jin, K. Kasavajhala, M.C. Kaymak, E. King, A. Kovalenko, T. Kurtzman, T.S. Lee, S. LeGrand, P. Li, C. Lin, J. Liu, T. Luchko, R. Luo, M. Machado, V. Man, M. Manathunga, K.M. Merz, Y. Miao, O. Mikhailovskii, G. Monard, H. Nguyen, K.A. O'Hearn, A. Onufriev, F. Pan, S. Pantano, R. Qi, A. Rahnamoun, D.R. Roe, A. Roitberg, C. Sagui, S. Schott-Verdugo, J. Shen, C.L. Simmerling, N.R. Skrynnikov, J. Smith, J. Swails, R.C. Walker, J. Wang, H. Wei, R.M. Wolf, X. Wu, Y. Xue, D.M. York, S. Zhao, and P.A. Kollman, Amber20. *University of California, San Francisco.* **2020**.
5. Hornak, V.; Abel, R.; Okur, A.; Strockbine, B.; Roitberg, A.; Simmerling, C., Comparison of multiple Amber force fields and development of improved protein backbone parameters. *Proteins* **2006,** *65* (3), 712-25.
6. Crespo, A.; Martí, M. A.; Kalko, S. G.; Morreale, A.; Orozco, M.; Gelpi, J. L.; Luque, F. J.; Estrin, D. A., Theoretical study of the truncated hemoglobin HbN: exploring the molecular basis of the NO detoxification mechanism. *Journal of the American Chemical Society* **2005,** *127* (12), 4433-4444.
7. Henriques, J.; Costa, P. J.; Calhorda, M. J.; Machuqueiro, M., Charge parametrization of the DvH-c3 heme group: validation using constant-(pH,E) molecular dynamics simulations. *J Phys Chem B* **2013,** *117* (1), 70-82.
8. Cruzeiro, V. W. D.; Amaral, M. S.; Roitberg, A. E., Redox potential replica exchange molecular dynamics at constant pH in AMBER: Implementation and validation. *J Chem Phys* **2018,** *149* (7), 072338.
9. Cruzeiro, V. W. D.; Feliciano, G. T.; Roitberg, A. E., Exploring Coupled Redox and pH Processes with a Force-Field-Based Approach: Applications to Five Different Systems. *J Am Chem Soc* **2020,** *142* (8), 3823-3835.
10. Jorgensen, W. L.; Chandrasekhar, J.; Madura, J. D.; Impey, R. W.; Klein, M. L., Comparison of simple potential functions for simulating liquid water. *The Journal of chemical physics* **1983,** *79* (2), 926-935.
11. Joung, I. S.; Cheatham III, T. E., Determination of alkali and halide monovalent ion parameters for use in explicitly solvated biomolecular simulations. *The journal of physical chemistry B* **2008,** *112* (30), 9020-9041.

12. Darden, T.; York, D.; Pedersen, L., Particle mesh Ewald: An N· log (N) method for Ewald sums in large systems. *The Journal of chemical physics* **1993,** *98* (12), 10089-10092.

13. Ryckaert, J.-P.; Ciccotti, G.; Berendsen, H. J., Numerical integration of the cartesian equations of motion of a system with constraints: molecular dynamics of n-alkanes. *Journal of computational physics* **1977,** *23* (3), 327-341.

14. Miyamoto, S.; Kollman, P. A., Settle: An analytical version of the SHAKE and RATTLE algorithm for rigid water models. *Journal of computational chemistry* **1992,** *13* (8), 952-962.

15. Gotz, A. W.; Williamson, M. J.; Xu, D.; Poole, D.; Le Grand, S.; Walker, R. C., Routine Microsecond Molecular Dynamics Simulations with AMBER on GPUs. 1. Generalized Born. *J Chem Theory Comput* **2012,** *8* (5), 1542-1555.

16. Jiang, X.; Burger, B.; Gajdos, F.; Bortolotti, C.; Futera, Z.; Breuer, M.; Blumberger, J., Kinetics of trifurcated electron flow in the decaheme bacterial proteins MtrC and MtrF. *Proceedings of the National Academy of Sciences* **2019,** *116* (9), 3425-3430.

17. Frisch, M. J.; Trucks, G. W.; Schlegel, H. B.; Scuseria, G. E.; Robb, M. A.; Cheeseman, J. R.; Scalmani, G.; Barone, V.; Petersson, G. A.; Nakatsuji, H.; Li, X.; Caricato, M.; Marenich, A. V.; Bloino, J.; Janesko, B. G.; Gomperts, R.; Mennucci, B.; Hratchian, H. P.; Ortiz, J. V.; Izmaylov, A. F.; Sonnenberg, J. L.; Williams; Ding, F.; Lipparini, F.; Egidi, F.; Goings, J.; Peng, B.; Petrone, A.; Henderson, T.; Ranasinghe, D.; Zakrzewski, V. G.; Gao, J.; Rega, N.; Zheng, G.; Liang, W.; Hada, M.; Ehara, M.; Toyota, K.; Fukuda, R.; Hasegawa, J.; Ishida, M.; Nakajima, T.; Honda, Y.; Kitao, O.; Nakai, H.; Vreven, T.; Throssell, K.; Montgomery Jr., J. A.; Peralta, J. E.; Ogliaro, F.; Bearpark, M. J.; Heyd, J. J.; Brothers, E. N.; Kudin, K. N.; Staroverov, V. N.; Keith, T. A.; Kobayashi, R.; Normand, J.; Raghavachari, K.; Rendell, A. P.; Burant, J. C.; Iyengar, S. S.; Tomasi, J.; Cossi, M.; Millam, J. M.; Klene, M.; Adamo, C.; Cammi, R.; Ochterski, J. W.; Martin, R. L.; Morokuma, K.; Farkas, O.; Foresman, J. B.; Fox, D. J. *Gaussian 16 Rev. A.03*, Wallingford, CT, 2016.

18. Shao, Y.; Gan, Z.; Epifanovsky, E.; Gilbert, A. T.; Wormit, M.; Kussmann, J.; Lange, A. W.; Behn, A.; Deng, J.; Feng, X., Advances in molecular quantum chemistry contained in the Q-Chem 4 program package. *Molecular Physics* **2015,** *113* (2), 184-215.

19. Jiang, X.; Futera, Z.; Blumberger, J., Ergodicity-breaking in thermal biological electron transfer? Cytochrome c revisited. *The Journal of Physical Chemistry B* **2019,** *123* (35), 7588-7598.

20. Wang, X.; He, X., An Ab Initio QM/MM Study of the Electrostatic Contribution to Catalysis in the Active Site of Ketosteroid Isomerase. *Molecules* **2018,** *23* (10), 2410.

21. Chen, C. G.; Nardi, A. N.; Amadei, A.; D'Abramo, M., Theoretical Modeling of Redox Potentials of Biomolecules. *Molecules* **2022,** *27* (3), 1077.

22. Guerard, J. J.; Tentscher, P. R.; Seijo, M.; Arey, J. S., Explicit solvent simulations of the aqueous oxidation potential and reorganization energy for neutral molecules: gas phase, linear solvent response, and non-linear response contributions. *Physical Chemistry Chemical Physics* **2015,** *17* (22), 14811-14826.

23. Ghosh, D.; Roy, A.; Seidel, R.; Winter, B.; Bradforth, S.; Krylov, A. I., First-principle protocol for calculating ionization energies and redox potentials of solvated molecules and ions: theory and application to aqueous phenol and phenolate. *The Journal of Physical Chemistry B* **2012,** *116* (24), 7269-7280.

24. Cruzeiro, V. W. D.; Amaral, M. S.; Roitberg, A. E., Redox potential replica exchange molecular dynamics at constant pH in AMBER: Implementation and validation. *The Journal of chemical physics* **2018,** *149* (7), 072338.

25. Karnaukh, E. A.; Bravaya, K. B., The redox potential of a heme cofactor in Nitrosomonas europaea cytochrome c peroxidase: A polarizable QM/MM study. *Physical Chemistry Chemical Physics* **2021,** *23* (31), 16506-16515.

26. Luo, R.; David, L.; Gilson, M. K., Accelerated Poisson–Boltzmann calculations for static and dynamic systems. *Journal of computational chemistry* **2002,** *23* (13), 1244-1253.

27. Lee, C.; Yang, W.; Parr, R., Density-functional exchange-energy approximation with correct asymptotic behaviour. *Phys. Rev. B* **1988,** *37* (2), 785-789.

28. Beck, A. D., Density-functional thermochemistry. III. The role of exact exchange. *J. Chem. Phys* **1993,** *98* (7), 5648-6.

29. Stephens, P. J.; Devlin, F. J.; Chabalowski, C. F.; Frisch, M. J., Ab initio calculation of vibrational absorption and circular dichroism spectra using density functional force fields. *The Journal of physical chemistry* **1994,** *98* (45), 11623-11627.

30. Hay, P. J.; Wadt, W. R., Ab initio effective core potentials for molecular calculations. Potentials for K to Au including the outermost core orbitals. *The Journal of chemical physics* **1985,** *82* (1), 299-310.

31. Ditchfield, R.; Hehre, W. J.; Pople, J. A., Self-consistent molecular-orbital methods. IX. An extended Gaussian-type basis for molecular-orbital studies of organic molecules. *The Journal of Chemical Physics* **1971,** *54* (2), 724-728.

32. Hehre, W. J.; Ditchfield, R.; Pople, J. A., Self—consistent molecular orbital methods. XII. Further extensions of Gaussian—type basis sets for use in molecular orbital studies of organic molecules. *The Journal of Chemical Physics* **1972,** *56* (5), 2257-2261.

33. Francl, M. M.; Pietro, W. J.; Hehre, W. J.; Binkley, J. S.; Gordon, M. S.; DeFrees, D. J.; Pople, J. A., Self-consistent molecular orbital methods. XXIII. A polarization-type basis set for second-row elements. *The Journal of Chemical Physics* **1982,** *77* (7), 3654-3665.

34. Gordon, M. S.; Binkley, J. S.; Pople, J. A.; Pietro, W. J.; Hehre, W. J., Self-consistent molecular-orbital methods. 22. Small split-valence basis sets for second-row elements. *Journal of the American Chemical Society* **1982,** *104* (10), 2797-2803.

35. Hariharan, P. C.; Pople, J. A., The influence of polarization functions on molecular orbital hydrogenation energies. *Theoretica chimica acta* **1973,** *28* (3), 213-222.

36. Roy, L. E.; Hay, P. J.; Martin, R. L., Revised basis sets for the LANL effective core potentials. *Journal of chemical theory and computation* **2008,** *4* (7), 1029-1031.

37. Krishnan, R.; Binkley, J. S.; Seeger, R.; Pople, J. A., Self-consistent molecular orbital methods. XX. A basis set for correlated wave functions. *The Journal of chemical physics* **1980,** *72* (1), 650-654.

38. McLean, A.; Chandler, G., Contracted Gaussian basis sets for molecular calculations. I. Second row atoms, Z= 11–18. *The Journal of chemical physics* **1980,** *72* (10), 5639-5648.

39. Scalmani, G.; Frisch, M. J., Continuous surface charge polarizable continuum models of solvation. I. General formalism. *The Journal of chemical physics* **2010,** *132* (11), 114110.

40. Smith, D. M.; Dupuis, M.; Vorpagel, E. R.; Straatsma, T., Characterization of electronic structure and properties of a bis (histidine) heme model complex. *Journal of the American Chemical Society* **2003,** *125* (9), 2711-2717.

41. Rovira, C.; Kunc, K.; Hutter, J.; Ballone, P.; Parrinello, M., Equilibrium geometries and electronic structure of iron− porphyrin complexes: A density functional study. *The Journal of Physical Chemistry A* **1997,** *101* (47), 8914-8925.

42. Kozlowski, P. M.; Spiro, T. G.; Bérces, A.; Zgierski, M. Z., Low-lying spin states of iron (II) porphine. *The Journal of Physical Chemistry B* **1998,** *102* (14), 2603-2608.

43. Smith, D. M.; Rosso, K. M.; Dupuis, M.; Valiev, M.; Straatsma, T., Electronic coupling between heme electron-transfer centers and its decay with distance depends strongly on relative orientation. *The Journal of Physical Chemistry B* **2006,** *110* (31), 15582-15588.

44. Johansson, M. P.; Sundholm, D.; Gerfen, G.; Wikström, M., The spin distribution in low-spin iron porphyrins. *Journal of the American Chemical Society* **2002,** *124* (39), 11771-11780.

45. Johansson, M. P.; Blomberg, M. R.; Sundholm, D.; Wikström, M., Change in electron and spin density upon electron transfer to haem. *Biochimica et Biophysica Acta (BBA)-Bioenergetics* **2002,** *1553* (3), 183-187.

46. McMahon, M. T.; DeDios, A. C.; Godbout, N.; Salzmann, R.; Laws, D. D.; Le, H.; Havlin, R. H.; Oldfield, E., An experimental and quantum chemical investigation of CO binding to heme proteins and model systems: a unified model based on 13C, 17O, and 57Fe nuclear magnetic resonance and 57Fe Mössbauer and infrared spectroscopies. *Journal of the American Chemical Society* **1998,** *120* (19), 4784-4797.

47. Zhang, Y.; Mao, J.; Oldfield, E., 57Fe Mössbauer isomer shifts of heme protein model systems: electronic structure calculations. *Journal of the American Chemical Society* **2002,** *124* (26), 7829-7839.

48. Havlin, R. H.; Godbout, N.; Salzmann, R.; Wojdelski, M.; Arnold, W.; Schulz, C. E.; Oldfield, E., An experimental and density functional theoretical investigation of iron-57 Mössbauer quadrupole splittings in organometallic and heme-model compounds: applications to carbonmonoxy-heme protein structure. *Journal of the American Chemical Society* **1998,** *120* (13), 3144-3151.

49. Kitagawa, S.; Morishima, I.; Yonezawa, T.; Sato, N., Photoelectron spectroscopic study on metallooctaethylporphyrins. *Inorganic Chemistry* **1979,** *18* (5), 1345-1349.

50. Liao, M.-S.; Scheiner, S., Electronic structure and bonding in metal porphyrins, metal=Fe, Co, Ni, Cu, Zn. *The Journal of Chemical Physics* **2002,** *117* (1), 205-219.

51. Listyarini, R. V.; Gesto, D. S.; Paiva, P.; Ramos, M. J.; Fernandes, P. A., Benchmark of Density Functionals for the Calculation of the Redox potential of Fe3+/Fe2+ within protein coordination shells. *Frontiers in chemistry* **2019,** *7*, 391.
52. Liao, M.-S.; Scheiner, S., Electronic structure and bonding in unligated and ligated FeII porphyrins. *The Journal of Chemical Physics* **2002,** *116* (9), 3635-3645.
53. Mao, Y.; Montoya-Castillo, A.; Markland, T. E., Accurate and efficient DFT-based diabatization for hole and electron transfer using absolutely localized molecular orbitals. *The Journal of Chemical Physics* **2019,** *151* (16), 164114.
54. Perdew, J. P.; Burke, K.; Ernzerhof, M., Generalized gradient approximation made simple. *Physical review letters* **1996,** *77* (18), 3865.
55. Weigend, F.; Ahlrichs, R., Balanced basis sets of split valence, triple zeta valence and quadruple zeta valence quality for H to Rn: Design and assessment of accuracy. *Physical Chemistry Chemical Physics* **2005,** *7* (18), 3297-3305.
56. Kubas, A.; Hoffmann, F.; Heck, A.; Oberhofer, H.; Elstner, M.; Blumberger, J., Electronic couplings for molecular charge transfer: Benchmarking CDFT, FODFT, and FODFTB against high-level ab initio calculations. *The Journal of chemical physics* **2014,** *140* (10), 104105.
57. Skourtis, S. S.; Waldeck, D. H.; Beratan, D. N., Fluctuations in biological and bioinspired electron-transfer reactions. *Annual review of physical chemistry* **2010,** *61*, 461-485.
58. Tipmanee, V.; Oberhofer, H.; Park, M.; Kim, K. S.; Blumberger, J., Prediction of reorganization free energies for biological electron transfer: A comparative study of Ru-modified cytochromes and a 4-helix bundle protein. *Journal of the American Chemical Society* **2010,** *132* (47), 17032-17040.
59. Sigfridsson, E.; Olsson, M. H.; Ryde, U., A comparison of the inner-sphere reorganization energies of cytochromes, iron− sulfur clusters, and blue copper proteins. *The Journal of Physical Chemistry B* **2001,** *105* (23), 5546-5552.
60. Matyushov, D. V., Protein electron transfer: is biology (thermo) dynamic? *Journal of Physics: Condensed Matter* **2015,** *27* (47), 473001.
61. Roe, D. R.; Cheatham III, T. E., PTRAJ and CPPTRAJ: software for processing and analysis of molecular dynamics trajectory data. *Journal of chemical theory and computation* **2013,** *9* (7), 3084-3095.
62. Skourtis, S. S., Reviewprobing protein electron transfer mechanisms from the molecular to the cellular length scales. *Peptide Science* **2013,** *100* (1), 82-92.
63. Jansson, F., Charge transport in disordered materials: simulations, theory, and numerical modeling of hopping transport and electron-hole recombination. **2011**.
64. Nenashev, A.; Jansson, F.; Baranovskii, S.; Österbacka, R.; Dvurechenskii, A.; Gebhard, F., Effect of electric field on diffusion in disordered materials. I. One-dimensional hopping transport. *Physical Review B* **2010,** *81* (11), 115203.
65. Moser, C. C.; Chobot, S. E.; Page, C. C.; Dutton, P. L., Distance metrics for heme protein electron tunneling. *Biochimica et Biophysica Acta (BBA)-Bioenergetics* **2008,** *1777* (7-8), 1032-1037.
66. Jiang, X.; Futera, Z.; Ali, M. E.; Gajdos, F.; von Rudorff, G. F.; Carof, A.; Breuer, M.; Blumberger, J., Cysteine linkages accelerate electron flow through tetra-

heme protein STC. *Journal of the American Chemical Society* **2017,** *139* (48), 17237-17240.
67. Jiang, X.; van Wonderen, J. H.; Butt, J. N.; Edwards, M. J.; Clarke, T. A.; Blumberger, J., Which multi-heme protein complex transfers electrons more efficiently? Comparing MtrCAB from Shewanella with OmcS from Geobacter. *The Journal of Physical Chemistry Letters* **2020,** *11* (21), 9421-9425.
68. Humphrey, W.; Dalke, A.; Schulten, K., VMD: visual molecular dynamics. *Journal of molecular graphics* **1996,** *14* (1), 33-38.
69. Autenrieth, F.; Tajkhorshid, E.; Baudry, J.; Luthey-Schulten, Z., Classical force field parameters for the heme prosthetic group of cytochrome c. *Journal of computational chemistry* **2004,** *25* (13), 1613-1622.
